# Smart Microscopy: Current Implementations and a Roadmap for Interoperability

**DOI:** 10.1101/2025.08.18.670881

**Authors:** Lucien Hinderling, Hannah S. Heil, Alfredo Rates, Philipp Seidel, Manuel Gunkel, Benedict Diederich, Thomas Guilbert, Rémy Torro, Otmane Bouchareb, Claire Demeautis, Célia Martin, Scott Brooks, Evangelos Sisamakis, Erwan Grandgirard, Jerome Mutterer, Harrison Oatman, Jared Toettcher, Andrii Rogov, Nelda Antonovaite, Karl Johansson, Johannes K. Ahnlinde, Oscar André, Philip Nordenfelt, Pontus Nordenfelt, Claudia Pfander, Jürgen Reymann, Talley Lambert, Marco R. Cosenza, Jan O. Korbel, Rainer Pepperkok, Lukas C. Kapitein, Olivier Pertz, Nils Norlin, Aliaksandr Halavatyi, Rafael Camacho

## Abstract

Smart microscopy is transforming life sciences by automating experimental imaging workflows and enabling real-time adaptation based on feedback from images and other data streams. This shift increases throughput, improves reproducibility, and expands the functional capabilities of microscopes. However, the current landscape is highly fragmented. Academic researchers often develop custom solutions for specific scientific needs, while industry offerings are typically proprietary and tied to specific hardware. This diversity, while fostering innovation, also creates major challenges in interoperability, reproducibility, and standardization, which slows progress and adaption. This article presents a collaborative effort between academic and industry leaders to survey the current state of smart microscopy, highlight representative implementations, and identify common technical and organizational barriers. We propose a framework for greater interoperability based on shared standards, modular software design, and community-driven development. Our goal is to support collaboration across the field and lay the groundwork for a more connected, reusable, and accessible smart microscopy ecosystem. We conclude with a call to action for researchers, hardware developers, and institutions to join in building an open, interoperable foundation that will unlock the full potential of smart microscopy in life science research.

## Introduction

In many areas of science, understanding complex natural phenomena increasingly relies on our ability to visualize them, sometimes down to the molecular level. In particular, microscopy and advanced imaging techniques have become crucial tools in biomedical research, enabling scientists to explore and document intricate biological structures and their dynamics across scales, from single biomolecules to entire animals. While some research questions require highly customised microscopy setups, a majority of imaging experiments are conducted using versatile, multipurpose imaging devices. The principles of smart microscopy apply broadly across imaging modalities, but this work focuses primarily on developments in light microscopy. Related efforts in fields such as electron microscopy face similar challenges in automation, interoperability, and data management [3], yet are beyond the scope of this paper.

Microscopy image acquisition workflows include positioning of the sample inside the microscope, finding the imaging target, and setting the acquisition parameters. These routine steps are often performed manually. While manual operation gives experimenters control over the acquisition process, it limits scalability and introduces potential bias, for example, by favoring fields of view that may not represent the full complexity of the sample. Automated microscopy has been actively developed for several decades in order to increase the throughput and has often focused on screening applications in which images of many samples are acquired with identical settings, enabling unbiased and systematic comparison of underlying specimens.

More sophisticated imaging experiments may require additional decisions being made during the acquisition process. For example, high-resolution imaging of a specific phenotype often begins with identifying the object of interest in a low-resolution overview, followed by switching to a high-resolution imaging modality. Similar requirements ap ply to different kinds of (photo-)manipulation experiments, in which the response of a sample is studied after actuating a specific area with a light stimulus.

**Fig. 1.**
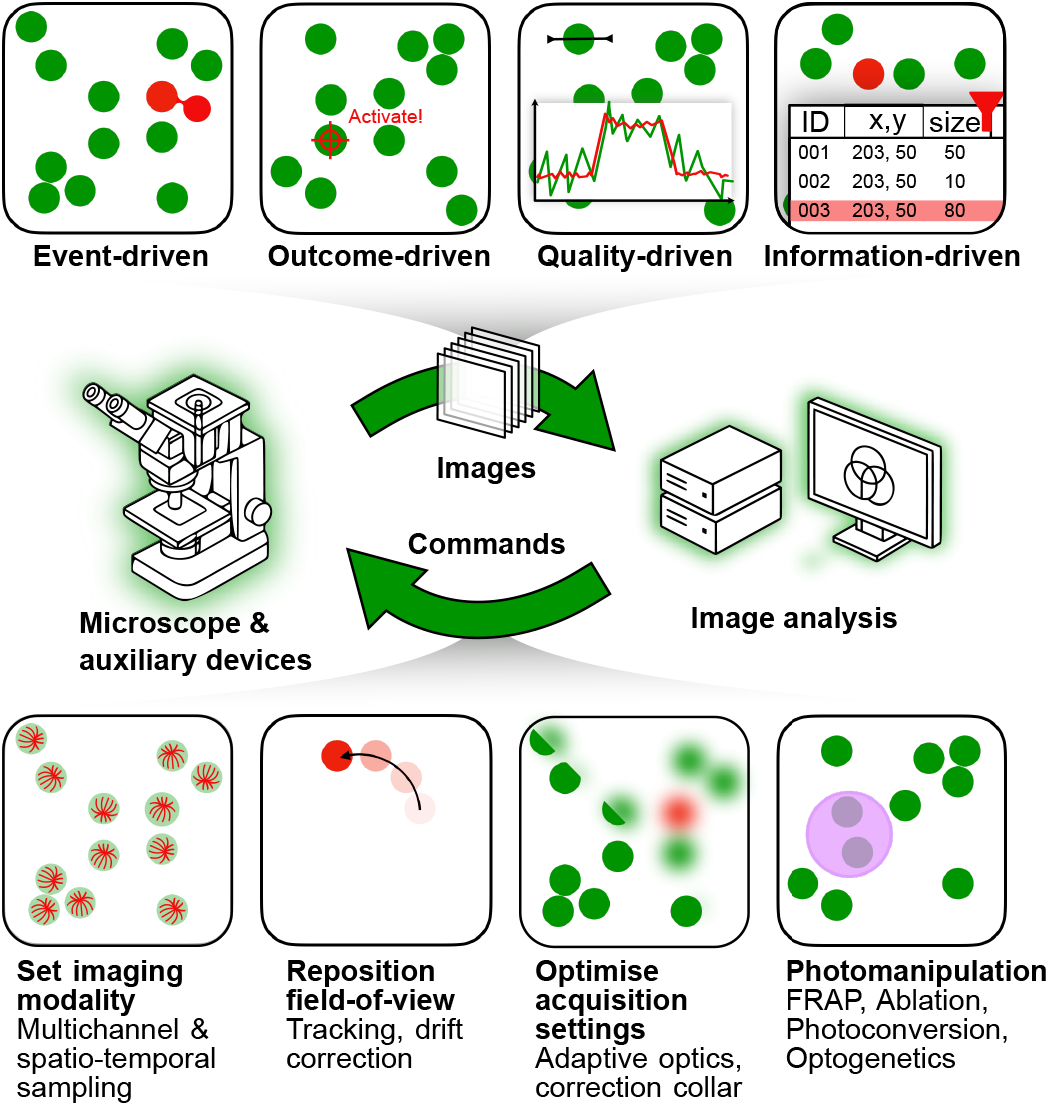
Categories and capabilities of smart microscopy systems integrating real-time image analysis and feedback control. **Top:** Smart microscopy workflows can be classified based on the driving logic behind decision-making: **Event-driven** (reacting to rare biological events), **Outcome-driven** (using feedback-control to steer biological systems toward a desired state), **Quality-driven** (optimizing signal quality or imaging metrics), and **Information-driven** (guided by models that predict which measurements/perturbations will yield the most informative data). **Middle:** Central feedback loop between the microscope and an image analysis system, which continuously exchanges images and commands to guide acquisition dynamically. **Bottom:** Key control actions enabled by smart microscopy: adjusting imaging modality (e.g. switching from brightfield to fluorescence, adjusting sampling rate), repositioning the field of view (e.g. tracking, drift correction), optimizing acquisition settings (e.g. adaptive optics), and performing photomanipulation (e.g. FRAP, ablation, optogenetics).

Advanced automation enables acquisition of large, information-rich datasets by on-the-fly processing of acquired data to trigger changes in the imaging or perturbation settings, ensuring that even the acquisition of large datasets can be reproduced, leaving only sample variability. Using live data to steer the acquisition or sample perturbation broadens the capabilities of microscopes to new types of experiments. This approach has the potential to revolutionise how scientists interact with and use microscopes in their research.

Different terms have been used to describe this emerging paradigm of running complex microscopy experiments without user supervision: smart microscopy, adaptive feedback microscopy, intelligent imaging, to name but a few. Throughout this document, we use the term **smart microscopy**. Despite promising technological advances and demonstrated impact on research [4, 5], the current landscape is fragmented, marked by diverse but often isolated solutions, as reflected in the wide range of naming conventions. Academic researchers have developed custom implementations based on either commercially available or self-built microscopes. While those successfully address specific scientific questions, they can often not be adapted to other experiments or microscope hardware systems.

As a response to these developments, many commercial microscope manufacturers started to create their own implementations, often integrated into existing proprietary software solutions and tailored to their hardware. While these efforts have enabled researchers to access ready-to-use smart microscopy solutions, many implementations still offer limited flexibility, particularly when it comes to extending and customizing experimental workflows or controlling devices not supported by the proprietary software. This article is the result of a collaborative effort between academic research groups and industry developers in the field of smart microscopy, brought together by the Smart Microscopy Working Group (SMWG)^1^ of EuroBioImaging^2^. The SMWG aims to coordinate community efforts toward standardization and development of interoperable smart microscopy techniques. We believe that such initiatives are essential for creating more unified and accessible solutions for future automated microscopy workflows.

Here, we provide an overview and categorisation of current smart microscopy implementations and applications. We compiled use cases and implementations from academic labs and microscope manufacturers, and present them in an accessible online resource designed to guide laboratories interested in adopting smart microscopy approaches. From this collection, we identify common themes, divergences, and challenges, and use these insights to propose a roadmap toward interoperability.

### Frequently Asked Questions

#### 1. What is smart microscopy?

Smart microscopy (also known as data-driven or adaptive-feedback microscopy) refers to imaging systems that can automatically adapt their behavior during acquisition. By combining real-time data/image analysis with fully motorized and computer-controlled microscopes, smart microscopy enables dynamic workflows: adjusting exposure, changing magnification, tracking the sample or triggering stimulation based on detected behavior. This approach can extend beyond the microscope, integrating it with environmental control, fluidics or sample handling systems, and also storage systems or downstream analysis pipelines.

#### 2. How do I get started?

Start small: choose one task you’d like to automate, find and zoom in on mitotic cells, or avoid photo-bleaching by adjusting illumination. Visit the SMWG website smartmicroscopy.github.io to e.g. find ready-made workflows, tutorials, tools and code examples. You don’t need to reinvent the wheel, many use cases have already been solved and shared by the community.

#### 3. Do I need to be a programmer?

Not necessarily. Many commercial platforms and some open-source tools offer ready to use workflows or scripting through user-friendly GUIs. If you want more flexibility, basic programming skills (often Python) can unlock more powerful workflows, but it’s not a barrier to entry.

#### 4. What if my microscope is from [X brand]?

Many commercial microscopes support advanced workflow control via macro programming, APIs or external scripting. A large number of systems are also compatible with µManager, an open-source device control framework supported by many academic smart microscopy tools. You can check hardware compatibility and software support in our overview tables (see Tables 2 and 3).

**Table 1.**
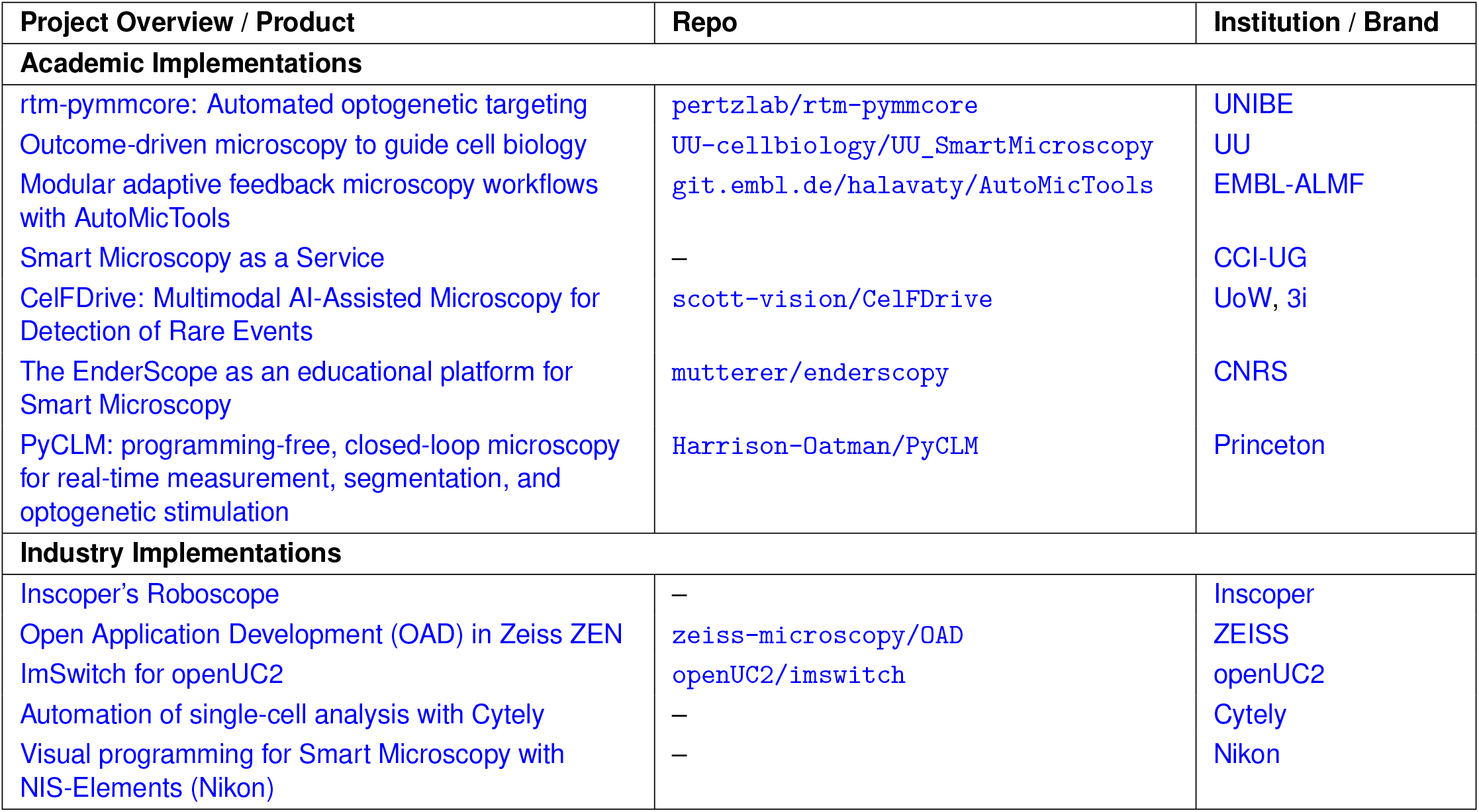
Smart Microscopy Implementations. Overview of the dataset compiling community-contributed smart microscopy implementations from academia and industry. Each entry includes a detailed description of the system, application examples, technical specifications, advantages and drawbacks, and links to code or repositories. While the supplementary information contains the full workflows at the time of publication, the [SMWG website](https://smartmicroscopy.github.io/implementations.html) provides the most current version.

**Table 2.**
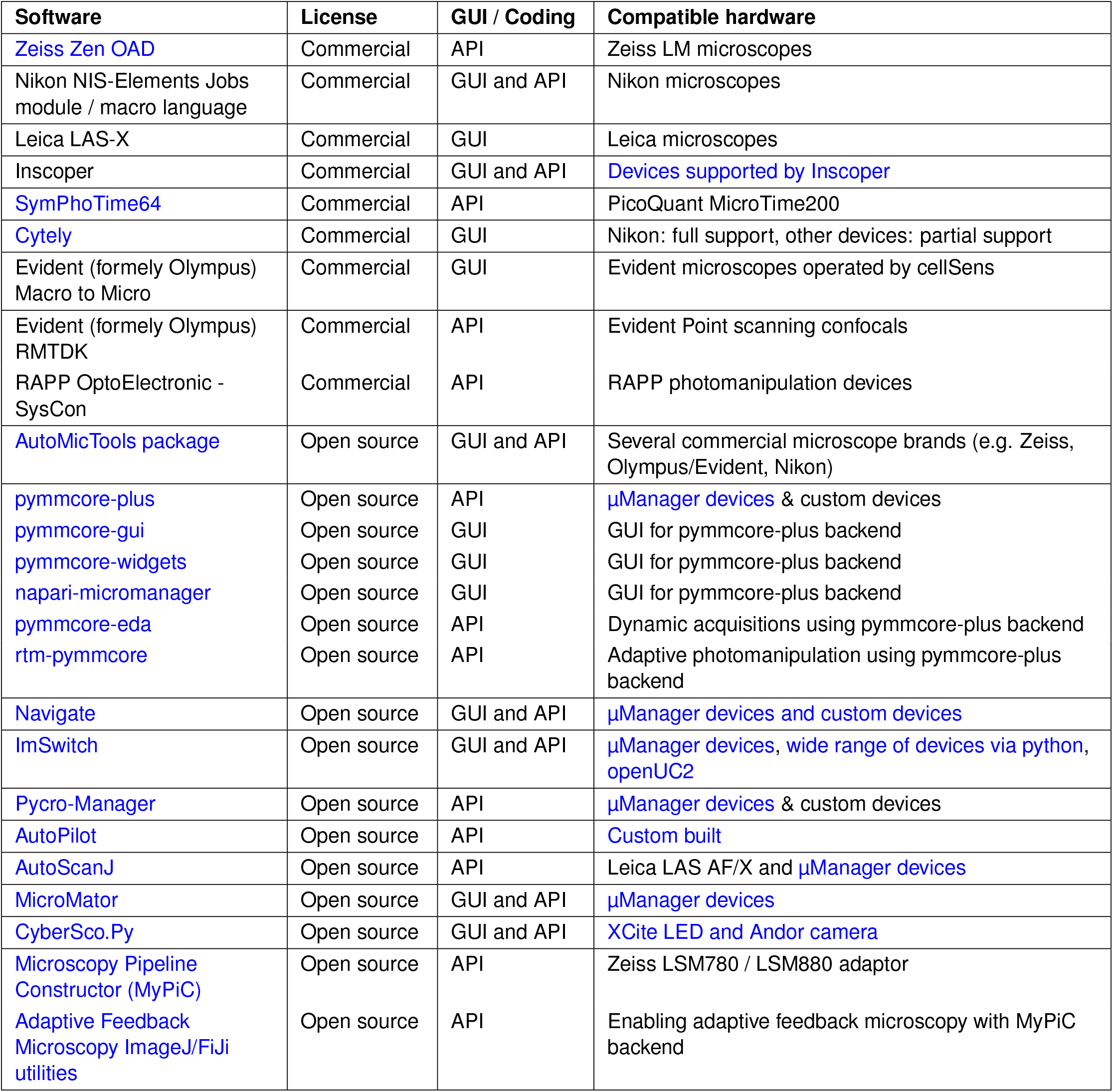
Software compatibility matrix. Comparison of microscopy control software with respect to openness, interface, integration, customization, and hardware compatibility. This feature matrix reflects the authors’ knowledge at the time of publication. For the latest software modules and features, see the [online version](https://smartmicroscopy.github.io/implementations.html).

**Table 3.**
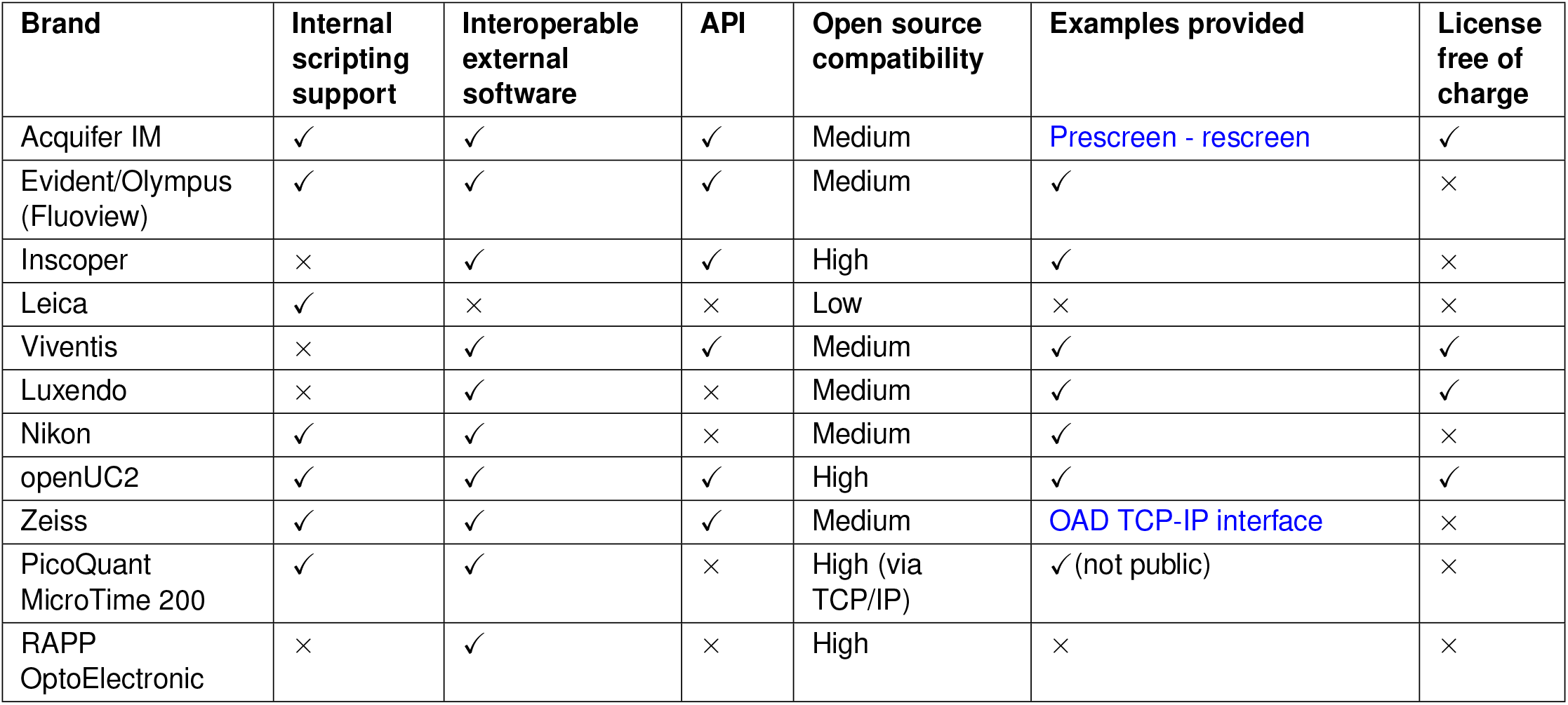
Hardware compatibility matrix. Comparison of microscopy hardware with respect to scripting support, interoperability, API availability, open source compatibility, documentation, and licensing. This table reflects the authors’ knowledge at the time of publication. For the latest updates on new hardware and features, see the [online version](https://smartmicroscopy.github.io/implementations.html).

| Brand | Internal scripting support | Interoperable external software | API | Open source compatibility | Examples provided | License free of charge |
| --- | --- | --- | --- | --- | --- | --- |
| Acquifer IM | ✓ | ✓ | ✓ | Medium | Prescreen - rescreen | ✓ |
| Evident/Olympus (Fluoview) | ✓ | ✓ | ✓ | Medium | ✓ | × |
| Inscoper | × | ✓ | ✓ | High | ✓ | × |
| Leica | ✓ | × | × | Low | × | × |
| Viventis | × | ✓ | ✓ | Medium | ✓ | ✓ |
| Luxendo | × | ✓ | × | Medium | ✓ | ✓ |
| Nikon | ✓ | ✓ | × | Medium | ✓ | × |
| openUC2 | ✓ | ✓ | ✓ | High | ✓ | ✓ |
| Zeiss | ✓ | ✓ | ✓ | Medium | OAD TCP-IP interface | × |
| PicoQuant MicroTime 200 | ✓ | ✓ | × | High (via TCP/IP) | ✓ (not public) | × |
| RAPP OptoElectronic | × | ✓ | × | High | × | × |

#### 5. Where can I learn more?

Join the SMWG, and check out these recent reviews:

- *The rise of data-driven microscopy powered by machine learning* Morgado et al. [1]: a concise primer on integrating machine learning with real-time acquisition.
- *Microscopes are coming for your job*, Pinkard and Waller [2]: forward-looking perspective on language-guided, fully autonomous microscopes.

Extensive literature and news can also be found at smartmicroscopy.org.

### Historical development

Innovations in optics and electronics over the past several decades have laid the groundwork for today’s automated microscopes. Key innovations, such as motorized stages introduced in the 1960s [6] and digital cameras and autofocus systems introduced in the 1980s enabled computer controlled acquisition, marking a first shift towards automation [7]. First efforts toward smart microscopy began with analog electronic circuits built for specimen tracking [8], soon digital approaches for single-cell segmentation followed [7, 9]. Innovations in image processing [10, 11], sample handling [12], and sample identification [13] further boosted automation capabilities. A pivotal turning point came with the introduction of computational microscopy, which enabled automated feedback-based control of imaging parameters and hardware components. By synchronising components like light sources and imaging stages, these systems enabled real-time adjustments, allowing for high-throughput data acquisition of dynamic phenomena with unprecedented precision and consistency [14, 15].

In the early 21st century, two key technical developments fueled the development of smart microscopy. First, lightsheet microscopy underwent rapid advancement. The massive datasets these microscopes acquire spurred the need for specialized algorithms for data handling, image processing, and most importantly, automation [16–19]. Second, artificial intelligence (AI), expanding across diverse fields, reached the microscopy community.

The integration of AI brought transformative solutions, not only for data and image processing through computer vision and data mining tools [20–24], but also by enabling a new quality of automatization, moving closer to the realization of the vision of smart microscopes as self-driving imaging robots that autonomously detect new biological principles [2, 25, 26].

### Current landscape and applications

Smart microscopy systems are fundamentally *data-driven*, meaning they adapt imaging decisions based on real-time experimental data or prior knowledge. These systems employ a wide range of strategies, tailored to specific experimental goals and constraints. Approaches range from passive observation (e.g. adjusting imaging parameters to minimize perturbations such as photobleaching or phytotoxicity) to active interventions that manipulate the biological system (e.g. using techniques like photoactivation, optogenetics, or microfluidics). Systems may respond to past or live events, or proactively attempt to predict and guide future outcomes. To provide an overview of current smart microscopy applications, we group them according to the *intent* behind their decision-making logic into four categories: **Event-driven, Outcome-driven, Quality-driven**, and **Information-driven**, as illustrated in Fig. 1. These categories are not mutually exclusive or fundamentally distinct frameworks; rather, they reflect different experimental priorities. In practice, many systems span multiple categories or integrate elements from more than one approach.

**Fig. 2.**
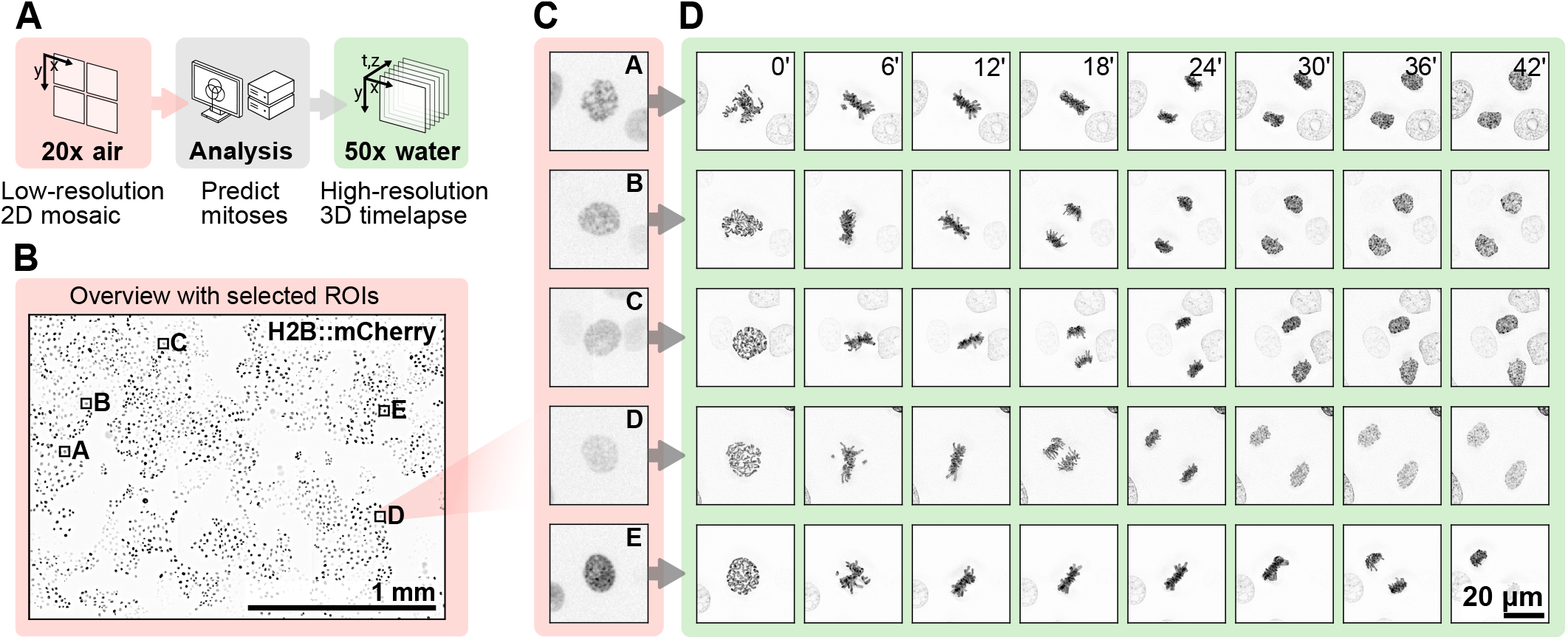
Example workflow of event-driven acquisition: automatic detection of mitotic events enables targeted high-resolution imaging of rare events. The smart acquisition workflow was completed entirely using Zeiss ZEN GUI tools, without the need for any custom coding. **A:** A low-resolution mosaic is acquired across the sample. A deep learning model identifies LLC-PK1 cells expressing H2B-mCherry in prometaphase, which are selected for high-resolution follow-up imaging. **B:** Overview image with detected regions of interest (ROIs) overlaid (20x air objective, 0.5x tubelens; background-subtracted). Cells express fluorescent histone H2B marker. **C:** Cropped regions from the overview image showing detected cells in prometaphase. **D:** High-resolution time-lapse imaging of selected cells undergoing mitosis (50x water auto-immersion objective, 2x tubelens; maximum projection of deconvolved z-stack).

### Event-driven

Event-driven imaging adapts acquisition in response to detected biological events, focusing data collection on relevant events while minimizing photodamage [27]. A representative example is the automated detection of mitotic cells, which triggers a switch to high-resolution imaging only during cell division (Fig. 2) [28]. This workflow, initially developed in research settings, is now becoming more widely accessible as commercial platforms begin to implement similar capabilities, reflecting a broader trend toward the integration of smart microscopy into routine imaging. Other research applications of event-driven imaging combine real-time analysis with dynamic adjustment of frame rate, illumination intensity, scanning mode, and many other parameters. Structured illumination microscopy (SIM) and lattice light-sheet microscopy (smartLLSM), enhanced with neural networks, switch between low-frequency background monitoring and fast, high-resolution imaging during events such as mitochondrial division, cell division, or immune synapse formation [29, 30]. Similarly, event-triggered STED nanoscopy captures vesicle trafficking and protein recruitment at high speed only when these processes are detected [31]. Event-driven control has also been applied to Brillouin microscopy, where a self-driving system uses deep learning to predict the onset of protein aggregation and trigger multimodal imaging, enabling realtime biomechanical characterization of aggregates in living cells while minimizing phototoxicity [32].At the population level, event-driven imaging can target phenotypes of interest across large samples [33]. One study used real-time characterization to identify migratory or infected cells and trigger high-resolution imaging only for those targets, capturing both rare and common events while preserving population context and reducing user bias [4]. These examples show how event-driven frameworks focus acquisition on relevant biological activity across scales, improving efficiency while maintaining image quality.

### Outcome-driven

Outcome-driven methods use the biological response of the specimen to dynamically adapt perturbation settings, forming a feedback-control loop between the microscope hardware and the biology under observation [35]. These approaches can leverage sensitivity to temperature or magnetism, but are more commonly used with light-based perturbations, such as optogenetics (Fig. 3). In optogenetics, cells are genetically engineered to express light-sensitive proteins, enabling precise control of cellular processes using light. When integrated with smart microscopy workflows, optogenetics enables the automatic targeting of subcellular structures like focal adhesions of just a few microns in size [36].

An illustrative example of outcome-driven microscopy is closed-loop optogenetic control of cell migration: real-time image analysis continuously monitors the position and orientation of migrating cells. When a cell begins to drift off its intended trajectory, the system triggers localized optogenetic stimulation to steer it back on course, essentially “nudging” the cell in real time to follow a predefined path [35].This approach can also be scaled to the tissue level, simultaneously directing the motility of many cells to form defined patterns like a “tissue printer” [37] or to investigate collective cell migration dynamics [38]. Another study demonstrated the use of optogenetics to manipulate cellular organelles, enabling precise control over organelle positioning and membrane trafficking. This allowed researchers to perturb organelle behavior at both global and localized scales without disrupting the overall state of the cell [39].

**Fig. 3.**
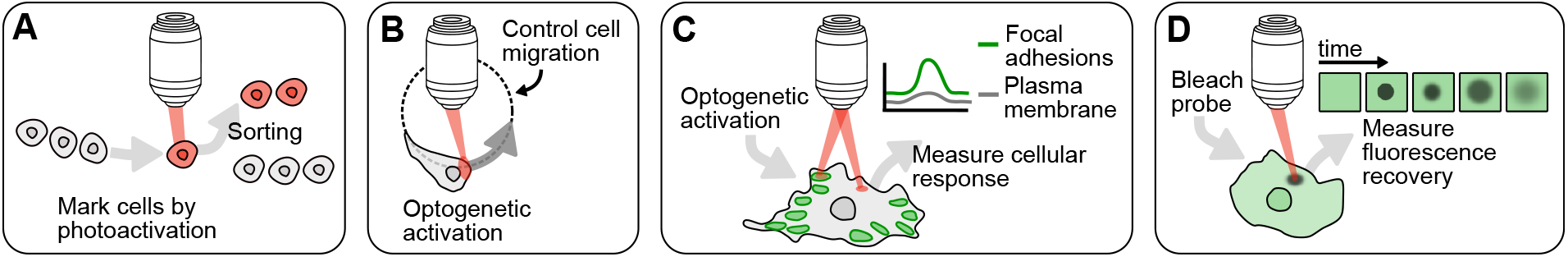
Examples of image-guided optogenetic control and photomanipulation. **A:** Cells are selectively marked based on spatial or phenotypic criteria via photoactivation and are subsequently sorted. **B:** Local optogenetic stimulation enables control of cellular migration. **C:** Automatic optogenetic targeting of subcellular structures (here focal adhesions and plasma membrane) to measure subcellular signaling heterogeneity. **D:** Similarly, image-guided feedback control in FRAP enables automated targeting and bleaching of dynamic subcellular structures, increasing experimental throughput [34].

Another application is the dynamic regulation of a cell’s signaling or gene expression state. Many cell signaling pathways are highly dynamic, with pulses or oscillations of pathway activity observed over time that are variable between cells [40–42]. By combining real-time measurements of signaling biosensors with optogenetic stimuli, signaling activity can be “clamped” at a constant level or made to follow a predefined dynamic trajectory, suppressing fluctuations and driving cells to a desired signaling state [43]. Engineered light-inducible transcription systems, such a light-responsive T7 RNA polymerases (Opto-T7RNAP) [44] and light-responsive two-component signaling systems [45, 46], enable precise control of gene expression in Escherichia coli. These systems offer and can be rapidly deactivated by removing the light or changing the stimulus wavelength, allowing tight temporal control over gene for the study of both synthetic and natural gene circuits. Dynamic gene regulation also enables feedback-driven control of protein production in response to changing environmental conditions: computer-controlled optogenetic platforms can be used to maintain target expression levels in bacterial cultures, despite fluctuations in factors such as nutrient availability or temperature. By modulating metabolic enzyme expression in real time, such systems can stabilize growth rates, highlighting their potential for applications in biotechnological production [47]. A closely related field is cybergenetics, which works towards engineered, computer-controlled biological systems [48–50].

These methods showcase how biological feedback can be harnessed to adaptively regulate experimental parameters, expanding control over complex cellular systems. They hold promise not only for fundamental research, but even in industrial biotechnology, where outcome-driven control could enhance process efficiency [51].

### Quality-driven

Quality-driven imaging in smart microscopy includes a range of strategies that adjust imaging parameters dynamically to maintain optimal image quality while minimizing damage to live or sensitive samples. These methods begin with straightforward mechanical adjustments and extend to sophisticated optical corrections and real-time computational feedback.

A simple yet effective example is the automated adjustment of the correction collar in confocal microscopy to compensate for variations in coverslip thickness. By analyzing axial image sharpness and intensity, the system fine-tunes the collar to optimize imaging across different samples and coverslips without manual intervention [52]. Adaptive optics offers another level of correction, addressing optical aberrations that vary within biological tissues. Using a guide star and direct wavefront sensing, this method enables adaptive correction with rapid update rates that are normally in the range of tens of milliseconds. This correction restores diffraction-limited imaging across large volumes, allowing detailed visualisation of neuronal structures and subcellular activity deep within the zebrafish brain [53, 54]. Tools like *AutoPilot* enhance quality by continuously aligning the light-sheet and detection planes in response to the specimen’s shape. This realtime geometric adaptation enables high-resolution, stable imaging in moving or developing organisms such as zebrafish and *Drosophila* embryos [18].

Quality-driven approaches can also significantly reduce light exposure during imaging. In adaptive scanning fluorescence microscopy, the system first maps a minimal subset of the sample (just 0.1%) to define the tissue surface of a developing *Drosophila* embryo. It then applies one of two targeted scanning strategies: focusing on a thin layer around the surface or imaging only specific structures such as adherens junctions. This method achieved up to an 80-fold reduction in light dose compared to traditional full-volume scans, enabling extended live imaging without compromising resolution [55]. At the computational side, deep learning-based image restoration techniques, such as Content-Aware Image Restoration (CARE), improve quality in low-light or undersampled conditions by enhancing fine details post-acquisition [56]. Recent approaches integrate such models directly into the imaging loop: one method combines real-time denoising with pixelwise uncertainty estimation to identify poor-quality regions and trigger targeted rescans. This strategy reduces light exposure and acquisition time by up to 16-fold while maintaining high detail [57]. This points to a new role for image restoration, not just in enhancing data but also in guiding real-time acquisition decisions. These quality-driven methods in smart microscopy highlight how adaptive control and computational feedback enable high-resolution imaging in large volumes and deep inside tissues.

### Information-driven

Information-driven approaches use statistical or machine learning models to guide imaging decisions. In this context, a model refers to a mathematical or computational representation of the biological system or process being studied, which can be used to make predictions, estimate uncertainty, and determine the most informative next steps. The aim is to acquire only new or relevant information in each image, reducing redundancy and increasing experimental efficiency. For example, a recent study applied this strategy to cell migration by combining broad population screening with model-informed targeting of extreme behaviors—specifically, the fastest- and slowest-moving cells. The system first performed a data-independent acquisition, capturing low-resolution, time-lapse images of approximately 25’000 cells to track movement across the population. It then used migration statistics, such as cell speed and directional persistence, to build a representation of cell behavior and identify the most distinctive subpopulations. Based on this model, the microscope selectively performed high-resolution imaging of only the most informative cells, minimizing unnecessary data collection without introducing human bias [4].

In active learning, such models are updated in real time to identify where additional data is most valuable, typically in regions of high uncertainty or where competing hypotheses could be distinguished. The system then prioritizes sampling accordingly. This contrasts with event- or quality-driven methods, which respond to immediate observations rather than supporting long-term inference. Similar strategies are already in use across other fields. In materials science, Gaussian process models guide scanning probe and electron microscopy by selecting perturbations that reduce model uncertainty [58, 59]. In systems biology, Bayesian inference and information theory are used to design experiments that maximize information gain and improve model identifiability and robustness [60]. While not yet common in biological research, these approaches provide a powerful framework for integrating modeling and decision-making into imaging workflows. Recent work has proposed systems in which microscopes use internal models to plan, test, and refine experimental hypotheses [2]. These approaches blur the line between model and experiment by incorporating a dynamic internal representation of the cell’s state to drive action [61]. As these systems advance, they may enable a new level of hypothesis-driven exploration in biological systems.

### Implementation strategies

The diversity of smart microscopy applications makes it difficult for a single software to cover all use cases. As a result, many research labs and imaging facilities rely on custom software and hardware solutions tailored to specific experiments, which are often difficult to reuse or adapt. While the field of smart microscopy was initially driven by these implementations, several commercial systems with smart microscopy features have since become available, making advanced imaging capabilities accessible to a broader user base. In this paper, we present a systematic collection of smart microscopy implementations, gathered from both academic institutions and commercial products contributed by members of the SMWG, spanning a variety of research applications and hardware configurations. We have created an online repository^3^ that compares different solutions, helping labs identify hardware and software that align with their research needs, available equipment, and coding expertise. This collection includes links to relevant code repositories and workflow examples to facilitate adoption. It is version-controlled and community-editable, allowing for expansion as new resources become available. The current documentation is included in the supplementary information (overview provided in Table 1). In the following sections, we summarize key implementation strategies among the collected examples and identify common challenges.

### Commercial implementations

Microscope manufacturers are increasingly integrating image analysis workflows with feedback control, as summarized in Table 3: **Hardware compatibility matrix**. These systems can dynamically adjust imaging parameters, such as increasing frame rates or switching to super-resolution modes upon detecting rare events like mitosis. They also optimize acquisition settings in real-time, for example, by increasing brightness when an image is too dark (e.g. Inscoper/Zeiss). For example Leica’s DMi8 and SP8 confocal microscopes, Zeiss’s AxioObserver and LSM 900/980 optionally equipped with AiryScan, Nikon’s Ti2 inverted and A1R HD25 confocal systems all integrate automated focus control, motorized stages, and software-based (often AI-assisted) feedback for live-cell imaging. These features improve resolution, minimize photobleaching, and enhance multi-wavelength time-lapse imaging. Similarly, Evident’s (formely Olympus’s) FV3000 confocal and VS120 slide scanner offer real-time image analysis and automated scanning, increasing throughput for applications like high-content screening. Modern microscopes include dedicated software for hardware control, visualization, and analysis, often featuring automation via scripting or graphical interfaces. This makes custom workflows accessible even to users without programming experience. Platforms like Inscoper’s Roboscope and Zeiss’s deep learning framework allow researchers to train their own neural networks for smart microscopy workflows, fine-tuned to their sample, without programming. To address the computational demands of image processing, some, such as Zeiss, incorporate GPU acceleration or offer cloud-based processing solutions.

**Fig. 4.**
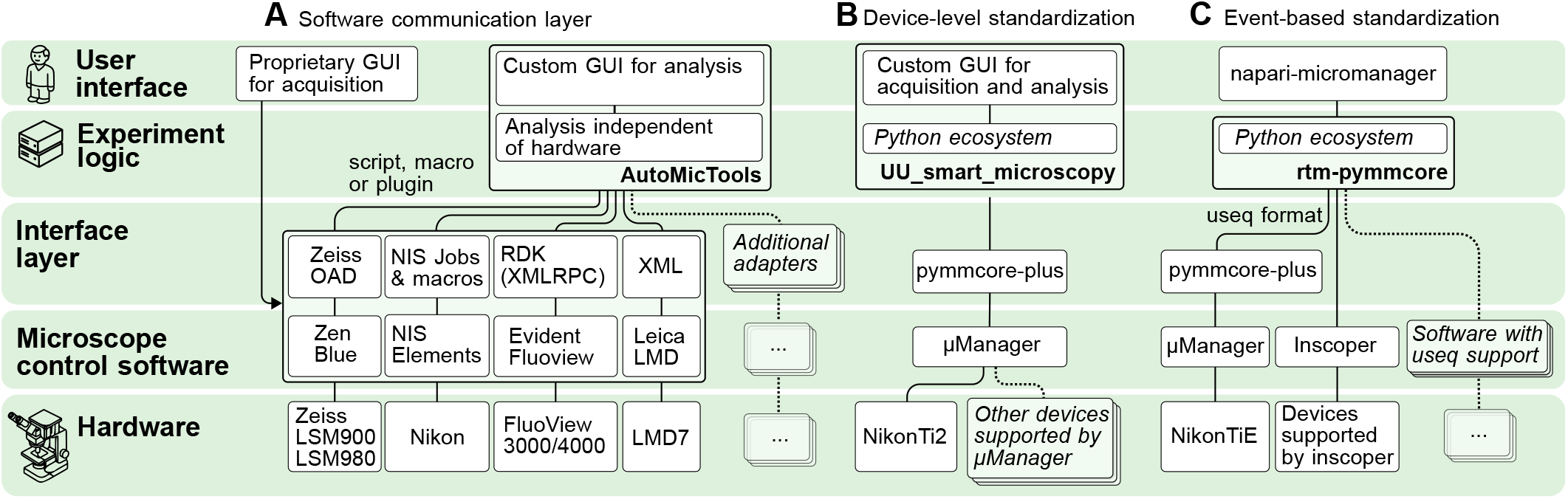
Strategies for hardware abstraction that allow smart microscopy workflows to run across different microscope systems, illustrated with example implementations collected on the SMWG website. **A: Software communication layers:** Image analysis and experiment logic are implemented in a platform-independent manner, while platform-specific adaptors control acquisition through proprietary microscope software via macros, APIs, or other interfaces. Custom GUIs allow users to configure analysis, while the vendor-provided software manages hardware and acquisition settings. By developing additional adaptors, these workflows can be extended to support microscope systems from other vendors. Example implementation: [AutoMicTools](https://smartmicroscopy.github.io/implementations/academic/embl.html). **B: Device-level standardization** (e.g. µManager-based workflows) bypasses proprietary GUIs and provides a unified API for direct device control across manufacturers. This API abstracts vendor-specific differences, enabling consistent control of a growing collection of supported hardware. Example implementation: [UU_smart_microscopy](https://smartmicroscopy.github.io/implementations/academic/uu.html). **C: Event-based standardization** decouples experimental design from hardware by describing acquisition events (e.g. acquire frame at *x, y, t* with channel *c*) in a structured format (e.g. useq-schema). Control software interprets these definitions and translates them into device-specific commands, enabling the use of vendor-specific features and optimizations during execution. Example implementation: [rtm-pymmcore](https://smartmicroscopy.github.io/implementations/academic/unibe.html).

To support external control and customization, some manufacturers provide APIs, allowing microscopes to be operated via Python bindings, REST APIs, or TCP/IP. However, only some companies provide free API access (like Luxendo and openUC2), while many still charge additional licence fees for this functionality. An important consideration when offering these interfaces is the accessibility of the scripting environment, for example the choice of programming language plays a critical role in usability. If users must learn a proprietary or unfamiliar language, or if the initial user base is small, the learning curve can discourage adoption. PicoQuant encountered this challenge with their SymPhoTime 64 software, which allowed users to create fully customized scripts for FLIM. Despite the flexibility this feature offered, less than 1% of users took advantage of it because the scripting language was unfamiliar. Lowering the entry barrier by using familiar and intuitive tools encourages more researchers to engage with advanced automation and customization features. PicoQuant has taken these lessons into account when designing their new system, Luminosa, and is considering support for widely used languages like Python as a priority for future development. Another important limitation of many provided APIs is the restricted availability of required commands and functions. Basic operations such as moving the stage or focus to a pre-defined position or switching between pre-defined acquisition settings are usually supported. However, more advanced features such as the programmatic definition of arbitrary ROI shapes for photomanipulation are often missing.

Recognizing the need for greater flexibility, an emerging trend is the shift from monolithic microscope control software to modular architectures. Some platforms, such as openUC2, support a “bring-your-own-control-software” model, allowing users to operate microscopes with tools like ImSwitch [62, 63], µManager [64, 65], pure serial interfaces, or Arkitekt [66]. This trend is also reflected in companies like Cytely, which offer software-only solutions for smart microscope control and data analysis. By decoupling software from hardware, such approaches highlight the growing importance of flexible, interoperable software infrastructure in modern microscopy. In parallel, support for open image formats, such as OME-TIFF or OMEZARR, adopted by openUC2, Inscoper, PicoQuant, and Zeiss [67], is further enabling integration with external tools and workflows.

### Academic implementations

In the collected examples, Python is the dominant programming language for implementing feedback control in academic settings. Its popularity stems from a versatile ecosystem that supports data analysis, deep learning, experiment logic, and GUI development within a single framework. Some solutions rely on vendor-provided software, controlling them either through built-in macro environments or via external APIs, with the latter offering greater flexibility. Unlike industry-provided graphical interfaces designed for general use, academic labs and facilities often create experiment-specific interfaces tailored to their unique workflows. Table 2: **Software compatibility matrix** provides an overview of available microscope control software.

A key challenge faced by academic implementations is expanding beyond a specific hardware platform. Due to the lack of standardized microscope hardware interfaces, each vendor provides distinct API functions. As a result, code written for one system cannot be easily transferred to another, leading to the need for intermediary software layers that abstract hardware-specific details and enable interoperability. These abstraction layers help decouple experiment logic from device-specific implementations, allowing to develop portable, reusable workflows. To address this problem, different labs have developed a variety of solutions, each operating at a different level of abstraction within the software stack (Fig. 4):

- **Software communication layers** act as an interface between hardware-agnostic analysis and decision-making tools and vendor-specific (often proprietary) microscope control software, accessed via macros, scripts, or plugins. In this setup, the analysis component can be reused across different microscope brands, while acquisition is coordinated by an additional adaptor program tailored to the specific control software. Data and notifications are exchanged between the analysis tool and the acquisition script via a predefined protocol. This approach, e.g. implemented by `AutoMicTools` and **MyPic** [68], allows to leverage vendor-specific capabilities. However, maintaining the adaptors requires continuous effort to support new devices and evolving APIs.
- **Device-level standardization**, exemplified by software like `µManager` [64, 65], provides a unified API for direct device control that abstracts over vendor-specific implementations. This enables consistent control of hardware such as Thorlabs and Nikon stages using the same code. It removes licensing constraints and promotes open access, but relies on the availability of open-source drivers and may limit device-specific features that fall outside the standard. This approach is mainly limited to self-built or commercial camera-based microscopes; low-level control of other microscope types (e.g. laser scanning confocals) is often restricted by companies.
- **Event-based standardization** uses structured formats, such as `useq-schema` [69], to describe acquisition events (e.g. acquire a frame at x, y, t with channel c) independently of the underlying hardware. Instead of issuing direct commands, `useq-schema` represents image acquisition as metadata, allowing different control software to interpret and translate these events into device-specific instructions. This approach delegates low-level implementation to hardware manufacturers, but requires their support of the standard to ensure interoperability.

Another commonality of the collected academic workflows is the heavy reliance on bioimage analysis methods for tasks such as segmentation [70, 71] and tracking [72, 73]. Because biological systems vary, these methods often need to be quickly customized. Modular, interchangeable software components make this possible. The Python ecosystem supports this flexibility with a rich set of libraries easily integrated into control workflows. Modularity supports rapid development and reuse of experiment logic across diverse imaging applications.

### Challenges

Smart microscopy not only increases data throughput but also generates **diverse data types** beyond traditional images, such as tracking data, segmentation masks, and biosensor quantification. This diversity necessitates flexible processing pipelines, robust metadata standards, visualisation tools, and scalable storage formats. The challenge is to develop computational solutions that can rapidly analyze these heterogeneous data streams while maintaining interoperability across different systems. The specificity of hardware setups, the **lack of standardized microscope control** APIs and metadata formats, and vendor lock-in make it difficult for academic labs to scale smart microscopy workflows across platforms. Incompatible software, closed firmware, and proprietary file formats complicate automation and integration. These barriers hinder reproducibility and force researchers to recreate similar solutions for each unique setup, slowing progress and increasing technical debt.

Despite facing similar challenges, research labs often develop solutions independently, leading to **redundancy**. A major obstacle to collaboration is the academic reward system, which prioritizes novelty and publications over contributions to shared tools. This discourages researchers from investing time in improving existing software when such work cannot easily lead to publishable outcomes. These structural disincentives fragment the field, undermining both collaboration and the robustness of methodological development.

On the industry side, companies have strong incentives to promote complete hardware-software ecosystems designed for broad usability. Ease-of-use is prioritized, often through pre-configured, integrated solutions rather than open systems that allow user-side customization. This model encourages users to purchase full microscope packages, including software contracts, locking them into proprietary ecosystems which **discourages adoption of open standards**. However, the development and use of open standards can shift demand toward modular, interoperable systems, pushing vendors to prioritize compatibility.

A significant challenge in smart microscopy lies in **balancing accessibility with complexity**. GUIs simplify adoption but limit flexibility, while coding-based approaches offer powerful customization but require programming skills many users lack. This divide seems to run parallel to the boundary between academia and industry: industry favors general, easy-to-use solutions, while academic labs tend to develop specialized tools tailored to narrow research questions. This split reinforces a broader challenge: tools that are accessible are often closed, while open tools remain underused due to technical barriers. Bridging this gap will require coordinated efforts across academia, industry, and open-source communities to develop systems that are both approachable and adaptable, without sacrificing interoperability or scientific depth.

### Towards interoperable smart microscopy

This section outlines the core components and community strategies needed to realize an interoperable smart microscopy ecosystem, as illustrated in Fig. 5.

### Community aspects

Smart microscopy is a rapidly evolving field with diverse expectations and requirements, making it difficult to establish a unified framework. Even basic terminology is not standardized, which creates barriers to collaboration and slows progress. To address this, our working group has taken a practical approach by organizing regular discussions and systematically collecting workflows that define core smart microscopy tasks such as: image acquisition → target detection → photoactivation. By comparing how such workflows are implemented across different hardware and software solutions, we aim to establish a shared vocabulary/terminology that improves communication and fosters interoperability.

One of the biggest obstacles to collaboration in smart microscopy is the fragmentation of its community. Since smart microscopy has been seen as a tool rather than a discipline so far, researchers working on these technologies often belong to a variety of disciplines and rarely find each other at the same conferences or publish in the same journals. While dedicated meetings on smart microscopy do exist, many contributors primarily identify with their biological subfields. As a result, new techniques are often published as part of broader biological studies, making them hard to discover, evaluate and adopt across disciplines. By systematically collecting and organizing approaches independent of specific biological questions, we aim to make smart microscopy methods more accessible to researchers from different backgrounds. Technical details and implementation examples are available on the SMWG website^4^. In addition, the SmartMicroscopy website^5,6^ hosts a curated, regularly updated list of publications related to smart microscopy. The site is maintained by SMWG members and welcomes suggestions from the community.

Building a strong community is critical at this stage. Our online repository and regular meetings not only serve as technical resources but also help connect labs working on similar challenges. By sharing insights and avoiding redundant development efforts, researchers can accelerate progress and make their solutions more adaptable for others. Public platforms such as image.sc^7^ and µForum^8^ demonstrate the value of open discussion spaces for troubleshooting, protocol sharing, and code reuse. However, company representation on image.sc is notably limited, as the forum’s moderation guidelines prioritize open-source projects and community-driven organizations.

A key part of this effort is thus strengthening ties between academia and industry. Academic laboratories often require flexibility to develop custom solutions, whereas industry tends to prioritize ease of use and seamless integration. These different priorities, however, are not necessarily in conflict. Through structured dialogue, researchers and industry partners can identify common needs and work toward more adaptable smart microscopy systems. Dedicated forums, such as the SMWG, provide a space where both communities can share challenges, present solutions and align priorities. By fostering open exchange, these initiatives help bridge the gap and promote greater interoperability.

Teaching smart microscopy differs from other disciplines due to its inherently interdisciplinary nature. Unlike traditional microscopy, many smart microscopy approaches require technical skills in programming and data analysis, which can create barriers for life science researchers without a computational background. Because smart microscopy projects involve diverse research questions and hardware setups, it is challenging to create standardized teaching materials. Effective training needs flexible approaches that combine biological topics with computational skills, preparing students to adapt what they’ve learned to their own hardware setups and specific research questions. In addition, economic accessibility is essential to building an inclusive smart microscopy community. Open-source projects like UC2 [74] or the Enderscope [75, 76] exemplify this by offering affordable, modular microscopes either 3D-printed and operated via smartphones or based or a 3D printer’s motorized 3 axis stage, while remaining fully compatible with advanced smart microscopy workflows. The modular teaching kits from openUC2 aim to lower the entry barrier with their open-source smart microscopy boxes offering motorized stage movement, fluorescence and open interfaces, where students first learn how to build and then to automate a complex microscopy system without the fear of damaging expensive hardware. Experimenting with custom written Python workflows on the Enderscope provides students with hands-on smart microscopy experience that they can easily transpose later on high end systems. Encouraging adoption of open and accessible platforms for education supports broader participation and collaboration.

Another cost-effective path is to retrofit existing equipment with modern control software. Many microscopy facilities rely on older systems that are optically and mechanically reliable but lack automation capabilities. These setups can often be upgraded using software tools; µManager [64, 65], for instance, supports a wide range of legacy devices and enables retrofitting systems with programmatic control. Python interfaces for µManager, such as pymmcore-plus [77] and pycro-manager [78], allow users to implement adaptive acquisition (see e.g. [27]), custom analyses, and feedback loops using modern software tools on older hardware – without the need for expensive upgrades.

**Fig. 5.**
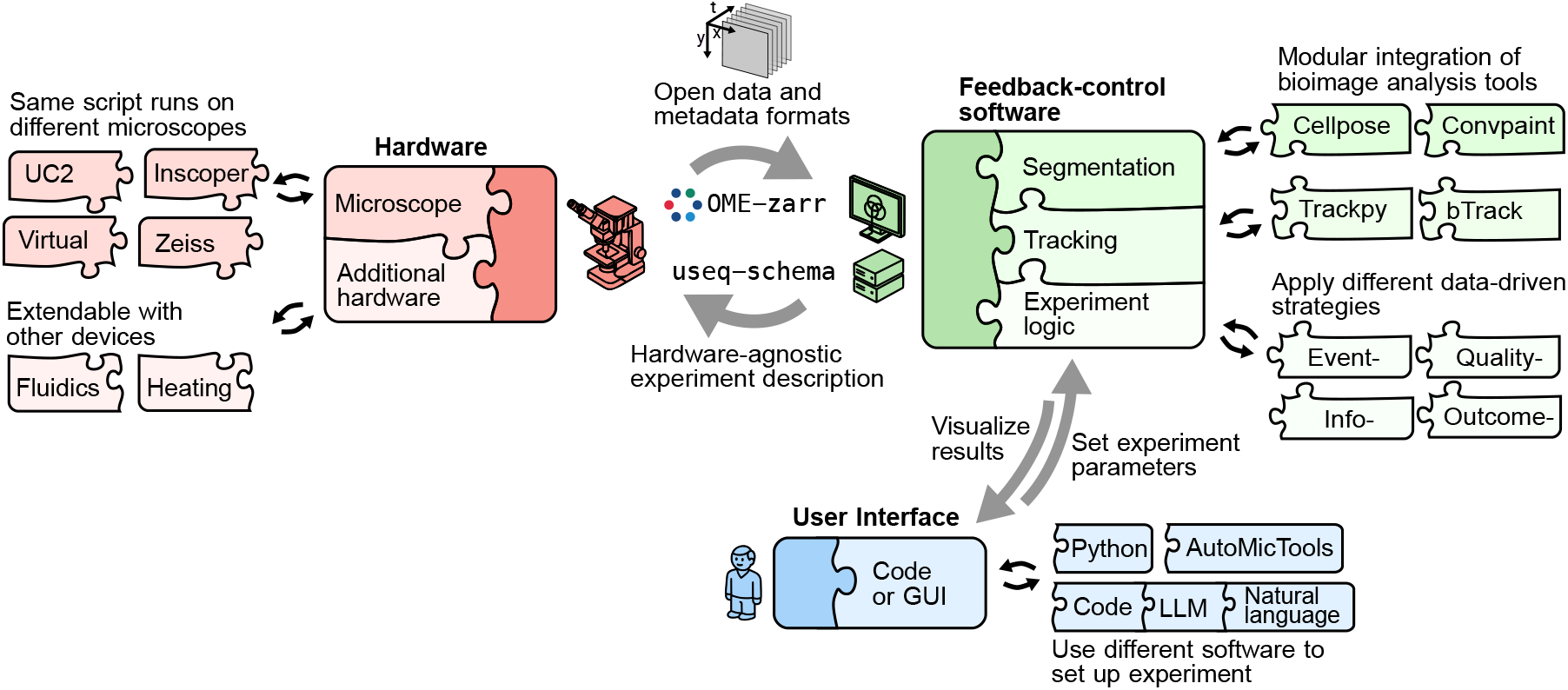
Interoperable smart microscopy ecosystem. Concept of a modular architecture for smart microscopy, where standardized experiment descriptions (e.g. useq-schema) and open data formats (e.g. OME-Zarr) allow integration of diverse microscopes, analysis tools, and user interfaces. Core components such as segmentation [70, 71], tracking [72, 73], and experiment logic are decoupled from specific hardware, enabling reuse across platforms. The system supports multiple input modalities (code, GUI, or natural language) and can be extended with additional devices like fluidics or environmental control modules. This structure enables flexible, feedback-driven acquisition strategies and cross-platform reproducibility.

### Formalizing workflows for interoperability

Smart microscopy development often starts with a specific biological question in mind. While this focus can lead to highly optimized solutions, the resulting tools are usually tightly coupled to a particular experiment or hardware configuration. As a results, they can be difficult to adapt or reuse in other research settings. This fragmentation limits interoperability and hinders collaboration across labs.

To address this, the SMWG is developing reference workflows based on collected implementation examples. These workflows describe experiments in terms of their objectives, acquisition logic, and expected outputs. Rather than being bound to a particular setup, these structured descriptions serve as adaptable blueprints for implementing software and defining hardware communication layers. By abstracting hardware-specific interactions within a common software infrastructure, they facilitate easier integration, reuse, and cross-platform compatibility. For example, a reference workflow for optogenetic stimulation and live imaging of cell migration might include: a description of the event-driven feedback loop (e.g. apply localized stimulation when a cell changes direction or slows below a threshold speed), along with general specifications for acquisition timing, resolution, and the structure of the expected output (e.g. image and tracking data).

These reference workflows can be implemented across diverse microscope platforms and software environments, providing a common benchmark for testing compatibility, evaluating performance, and identifying integration gaps. The SMWG will also provide reference implementations demonstrating how each workflow can be realized using different software and hardware configurations. By curating, documenting, and publishing these workflows and examples, the SMWG aims to establish a shared reference that fosters modular system development, promotes collaboration, and reduces duplication of effort across the community.

#### Metadata standardization and logging

Standardized metadata, interfaces, and communication protocols are essential for formalizing interoperability between different software and hardware systems in smart microscopy. On the acquisition side, structured metadata schemas like useq-schema [69] help researchers define experiments independently from the underlying hardware. This separation simplifies the sharing and adaptation of imaging protocols across laboratories. On the data side, OME-Zarr [67] is emerging as a key standard for handling the diverse and complex datasets produced by smart microscopy. Unlike traditional microscopy, which primarily generates image stacks, smart microscopy additionally outputs various data types including tracking data, segmentation masks, and biosensor measurements. OME-Zarr seamlessly integrates different data types, enabling efficient storage, cloud-based sharing, and real-time visualization. Enforcing well-defined metadata conventions allows downstream image processing and analysis tools to be reused across platforms without time-consuming conversions.

Community initiatives play a critical role in advancing metadata standards. The QUAREP-LiMi (Quality Assessment and Reproducibility for Instruments & Images in Light Microscopy) consortium promotes practical metadata standards such as the NBO Microscopy Metadata model [79], facilitating metadata capture and integration through user-friendly tools and educational resources. The NBO-Q standard, developed collaboratively by the 4D Nucleome project, BioImaging North America, OME, and QUAREP-LiMi, extends the OME Data Model with a tiered structure that adapts to varying experimental complexities. Tools like the Micro-Meta App [80] enable researchers to efficiently collect and structure metadata, enhancing reproducibility and interoperability across imaging platforms.

**Fig. 6.**
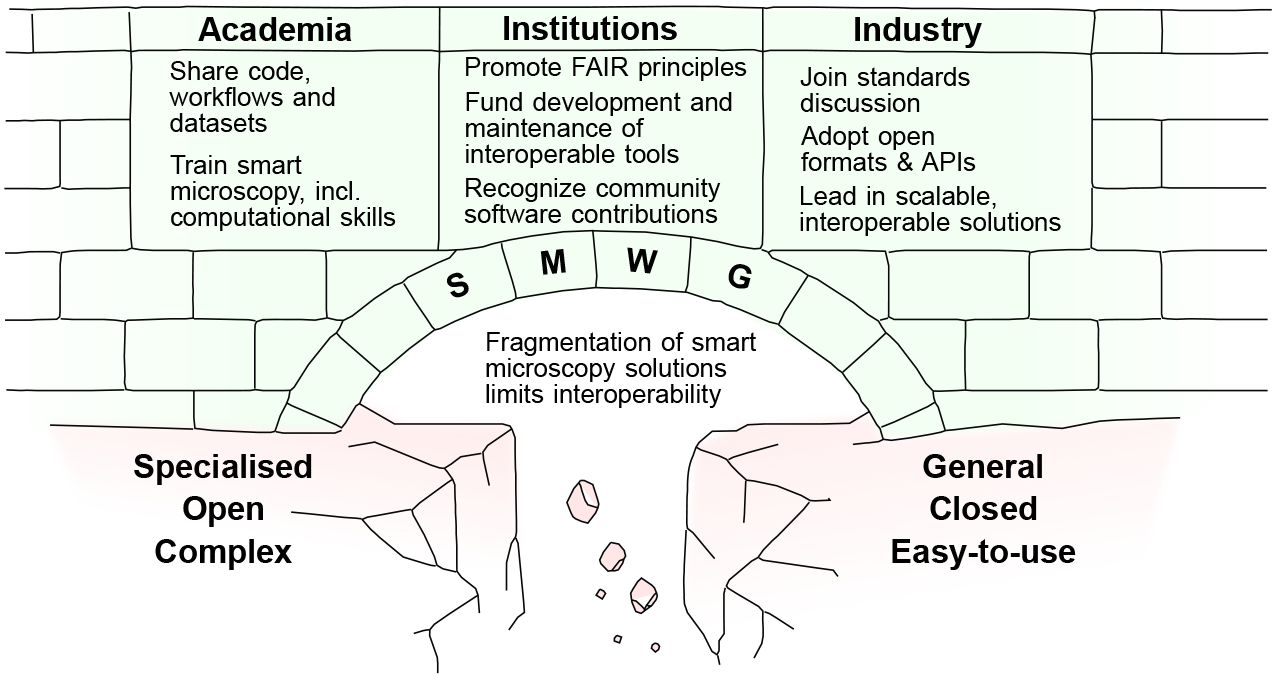
Shared participation of academia, institutions and industry is required to bridge between different smart microscopy solutions.

However, achieving widespread adoption of metadata standards faces significant challenges, especially due to varying levels of support among commercial hardware vendors. Companies must balance innovation and unique capabilities with standardized, interoperable designs. Differences in hardware architecture often make direct metadata mappings difficult. Consequently, the focus is shifting away from strict uniformity toward practical interoperability. Formats such as OME-Zarr represent a middle ground, enabling different systems to collaborate through shared conventions rather than identical internal implementations.

Transparent, machine-readable logging of actions taken during acquisition and analysis is another critical component of smart microscopy. Detailed logging complements structured metadata to ensure reproducibility, traceability, and efficient automated processing. Traditionally, metadata (system states and acquisition parameters) and workflow descriptions (human-readable experimental intent) have been distinct. However, as microscopy becomes increasingly smart, these boundaries may blur: Experimental intent, system states, and execution could become closely integrated, requiring metadata structures to evolve accordingly.

#### In silico testing of smart microscopy workflows

To fully leverage standardized workflows and data formats, smart microscopy systems require testing environments where researchers can evaluate and refine their approaches before conducting experiments on physical microscopes. Virtual microscopes play a crucial role in this process by providing a controlled setting to test imaging strategies, feedback loops, and analysis algorithms. To be truly effective, they must go beyond simple image simulations and accurately model both microscope behavior and how the sample responds to perturbations, such as changes in the field of view with stage movement or reactions to photostimulation. When combined with standardized experiment descriptions, virtual microscopes enable systematic verification of experiments across different hardware and software platforms. They also make it possible to write tests for complete smart microscopy experiments as part of the software development cycle, ensuring the same level of rigor expected from any complex codebase.

#### Using LLMs to translate workflows and serve as user interface

As large language models (LLMs) continue to advance and demonstrate utility in fields such as robotics [81], a key question emerges: how might these models be effectively leveraged in smart microscopy [2]?

In this context, LLMs could serve as intermediaries between user intent and system execution. A researcher might describe an experimental protocol in natural language, which the system could then translate into structured, hardware-specific commands. This translation layer would enable protocol portability across microscope platforms, reduce the friction of adapting workflows across labs, and lower the barrier for users without extensive programming experience.

LLMs also offer a novel way to resolve the longstanding trade-off between graphical interfaces and the flexibility of scripting. Rather than designing static, one-sizefits-all UIs, future systems could use LLMs to dynamically generate task-specific controls or code templates tailored to the user’s current experiment. Recent work has demonstrated this approach in bioimage analysis, where LLMs have been used to create on-the-fly graphical interfaces and analysis workflows based on natural language input [82].

Embedding language-based interaction into smart microscopy software architectures could lead to systems that are more accessible, adaptable, and user-centered. Looking ahead, it may be useful to consider what it means for a system to be “LLM-ready”, that is, designed to effectively integrate with LLMs to enable intelligent, interactive control. Key prerequisites include structured, machinereadable metadata and detailed logs of user actions — both of which can serve as training data for systems that progressively learn from expert interactions. This opens the door to semi-autonomous agents capable of assisting with experiment design, optimizing acquisition parameters, detecting anomalies, or automating routine tasks.

Realizing these capabilities in scientific workflows will require careful attention to transparency, validation, and reproducibility to maintain scientific rigor and user trust.

While technical and ethical challenges remain, the integration of LLMs into smart microscopy systems represents a promising direction for making complex automation workflows widely usable.

### Conclusion

Smart microscopy holds immense potential to accelerate discovery by combining advanced imaging, automation, and AI. Yet this potential remains limited by fragmentation across platforms, tools, and institutions. Achieving broad impact will require a shared commitment to interoperability across technical, cultural, and organizational dimensions (Fig. 6).

Key challenges include the absence of standardized workflows and metadata schemas, the prevalence of proprietary solutions that inhibit integration, and misaligned incentives between academia, industry, and infrastructure providers. These barriers slow development, hinder reproducibility, and limit reuse of valuable tools and data.

Over the next year, the Smart Microscopy Working Group (SMWG) will focus on three priorities: (1) defining and publishing baseline interoperability specifications; (2) curating an open, evolving collection of shared implementations, datasets, and workflows; and (3) organizing community events to facilitate dialogue and alignment across academia, industry, and research infrastructures/institutions. These efforts aim to transform fragmented innovations into coherent, and reusable ecosystem for smart microscopy. To achieve this, we invite all stakeholders to take action:

- **Academic labs and imaging facilities:** Share workflows, datasets, and modular code to public platforms such as the SMWG repository; integrate computational methods into training programs.
- **Industry partners:** Provide accessible APIs; participate in the development of shared specifications; adopt open formats like OME-Zarr and useq-schema.
- **Institutions (publishers, funders, infrastructure providers):** Incentivize FAIR practices; formally recognize contributions to open-source software and shared workflows; invest in the long-term sustainability of shared tools and platforms.

The SMWG is committed to enabling open collaboration, supporting shared technical resources, and coordinating efforts to define common standards. Participation is flexible: you can join a discussion, present a project, submit a resource, or simply stay informed. Every contribution, large or small, strengthens the foundation for scalable, open smart microscopy.

Advancing interoperability is a shared responsibility. It is a critical step toward unlocking the full potential of smart microscopy as a robust, widely adopted tool for discovery. This is both a technical and a cultural challenge, but one we can meet together.

## Acknowledgments

We thank Euro-BioImaging for support of the SMWG. We thank EMBL for support of the SWMG. This work has been supported by Uniscientia fellowship 187-2021 and SNF Sinergia grant CRSII5_183550 to OP. RC acknowledges support from the Swedish Foundation for Strategic Research (SSF) through the RIF-2 program and the Swedish National Microscopy Infrastructure, NMI (VR-RFI 2023-00163). HSH acknowledges the support by the European Union’s Horizon 2020 research and innovation program under the Marie Sklodowska-Curie grant agreement no. 101204120 (https://doi.org/10.3030/101204120). NN thanks the Swedish research Council (2023-05450), the IngaBritt and Arne Lundberg foundation, the Crafoord foundation, the Seydlitz foundation, and the Magnus Bergvall foundation for support. PN acknowledges the support of the Berta Kamprad Foundation, the Swedish Cancer Foundation (Cancerfonden), Swedish Research Council (Vetenskapsrådet, 2019-02355 & 2023-02387) and the the Knut and Alice Wallenberg Foundations. AR and LK acknowledge the support from the Netherlands Organization for Scientific Research (NWO) through the National Roadmap Initiative NL-BioImaging-AM

## Methods

Images in fig. 2 were acquired on a CellDiscoverer 7/7 Rev. 2 microscope. The cell line was pig kidney LLCPK1 expressing H2B-mCherry and *α*Tubulin-GFP (only H2B shown). Cells were maintained in MEM alpha supplemented with 10% FCS, 100 U/mL streptomycin, and 100 µg/mL penicillin under standard culture conditions (37 ^*°*^C, 5% CO_2_, 95% humidity). Overview scans were acquired in widefield mode with a 20x air objective and a 0.5x tube lens; detail scans were also acquired in widefield mode with a 50x water-immersion objective and 2x magnification.

## rtm-pymmcore: Automation of optogenetic targeting

**Lucien Hinderling, Benjamin Grädel, Alex Landolt, Maciej Dobrzinsky, Olivier Pertz**, University of Bern, Institute of Cell Biology

### Specific Focus and scientific questions asked

The cytoskeleton drives key cellular functions like morphology, polarity and migration, and is regulated by Rho GTPases that act as molecular switches. Traditionally, the Rho GTPases Rac1, Cdc42, and RhoA were thought to independently control actin polymerization, filopodia formation, and contractility in a top-down manner. Recent biosensor studies have revealed that these GTPases are however dynamically coordinated at the subcellular scale, interacting in complex signaling networks at the timescale of seconds. Feedback from cytoskeletal structures directly influences Rho GTPase activity, creating an intricate, reciprocal relationship between signaling and structure.

Optogenetics allows precise spatio-temporal control of Rho GTPase signaling, enabling direct causal testing of this feedback regulation, by shining light on specific cellular regions and measuring the signaling response. Applying optogenetic stimulation in live cells requires constant manual adjustment of the illuminated regions to follow cellular movements. This approach is thus time-intensive and limited in throughput, posing a challenge for studying cytoskeletal signaling in dynamic contexts.

Smart microscopy addresses these limitations by integrating optogenetics with automated imaging systems that use digital mirror devices (DMDs) or galvo mirrors to dynamically update illumination patterns. Using computer vision, these microscopes autonomously detect cytoskeletal structures and respond to cellular behavior. Adapting light stimulation patterns in real time, this approach allows to systematically probe local heterogeneity in cellular signal processing (Fig. 1A).

**Fig. 1:**
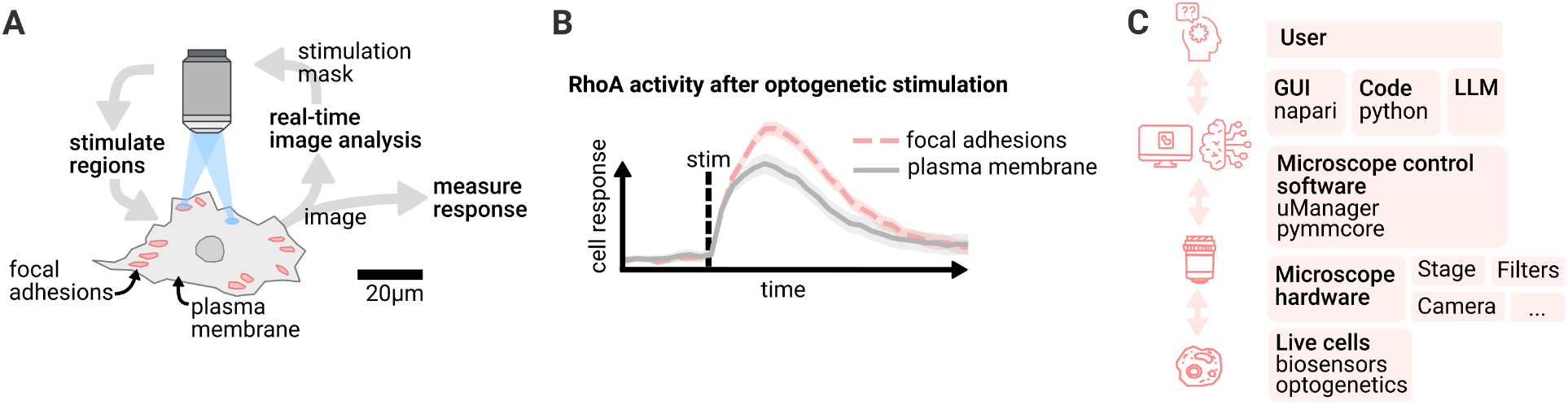
Smart microscopy for automated subcellular targeting: **A)** Cytoskeletal structures are detected with computer vision and targeted with optogenetic stimulation. Local response of the cells is measured. **B)** Cell response depends on stimulation location, highlighting subcellular heterogeneity in cellular signal processing. **C)** Overview of architecture. The user interfaces with the microscope control software via GUI, code or by using an LLM. The microscope controls live cells using optogenetics and reads their response using fluorescent biosensors.

#### Key findings and innovations

In a recent study exploring feedback regulation between Rho and a cytoskeletal structure called focal adhesions, we demonstrated the capability of this approach [1]. Using an optogenetic actuator-biosensor circuit to activate Rho and measure Rho activity, we investigated DLC1, a regulator of Rho that localizes to focal adhesions.

By automatically detecting focal adhesions, we systematically analyzed cellular responses to Rho activation, probing thousands of subcellular regions across hundreds of cells in a fully automated manner (Fig. 1B). With this high-throughput approach, we found that DLC1 maintains force homeostasis at focal adhesions by locally increasing Rho activity in response to mechanical stress. With the manual low-throughput methods we used initially, the regulatory dynamics were masked by cell-to-cell variability.

Together, optogenetics and smart microscopy offer a scalable, high-precision method to dissect complex feedback-driven signaling networks, providing novel insights into cytoskeletal regulation. By swapping out the optogenetic actuator and biosensor circuit, this approach can be adapted to study spatio-temporal signaling dynamics across various domains of cell biology.

#### Methodology and implementation details

To control and automate the microscope we built an open-source software stack (Fig. 1C). Our primary challenge was integrating control of all hardware devices (camera, stage, light source, DMD) within a single software environment. We build upon Micro-Manager, which provides drivers for over 250 devices. For feedback logic and data analysis, we chose Python due to its rich libraries for computer vision and signal processing. Python’s accessibility and feature-rich ecosystem make it an ideal choice. We bridge Micro-Manager’s C++ device drivers to Python using PyMMCore.

Our smart microscopy experiments rely on complex image processing pipelines that generate diverse outputs (segmentation masks, point detections, tracks, biosensor quantification). These pipelines have to be adapted to cellular variability: cell appearance, biosensors, and morphology differ from experiment to experiment based on factors like drug treatment. Since we need real-time image processing during experiments, there is no time for extensive model training or post-acquisition parameter tuning. To address this, we built Convpaint, an interactive pixel classifier that can be trained on a single sample image from that experiment. We use Napari, a multimodal image viewer for Python, to preview the image processing results, facilitating on-the-fly parameter tuning before starting the experiment.

Napari also allows us to create custom microscope control widgets (e.g., for previewing image analysis, calibrating the DMD, setting experimental parameters). This setup enables even complex smart microscopy experiments to be run without coding. While commercial microscope control software provides extensive GUIs for standardized experiments, custom experiments usually require scripting. With Napari, we can build GUIs to streamline accessibility even for highly customized experiments and technically complex setups. Recently a plugin has been developed (napari-micromanager) that provides a GUI for all (non-smart) microscope operations.

Each experiment is documented in a Jupyter notebook, detailing algorithms and experiment variables like the stimulation logic. This practically provides automated documentation. Re-running an experiment is as simple as re-running the notebook.

To maximize experimental throughput, we use a main thread for scheduling hardware control events (move stage, acquire image, stimulate region), while image processing runs on separate threads for different fields of view.

All image data and processing results are continuously stored, allowing inspection at any time during the experiment and faster iteration time when troubleshooting. From the extracted data, we also automatically generate quality control plots (segmentation overlays, tracking statistics). At the end of each experiment, plots can be created from the microscope output data with no further processing. All code is available on github, more details and application examples can be found in the paper [2].

#### Contributions to Interoperability

Micro-Manager not only offers extensive device driver support (see list of supported devices), making our workflow compatible with a wide range of custom microscope setups globally, but it also provides standardized interfaces for the core device types in modern microscopy. This framework ensures that our code remains hardware-agnostic; for instance, any DMD can be seamlessly integrated as long as it is implemented as a genericSLM device in Micro-Manager (we currently use the Andor Mosaic 3 and Mightex Polygon 1000). We define acquisition events in an abstract format using useq-schema, which could theoretically be adapted for use with other microscope control software.

#### Limitations

These abstractions are feasible due to a growing community around Micro-Manager. Sustained development and critical mass forming around its ecosystem encouraged many hardware vendors to make their devices compatible with the software. However, the current abstractions are designed for classical microscope automation without feedback components. All smart features in our pipeline remain outside any established standards. Currently, we rely on custom data structures to store results. We plan to implement OME-Zarr to standardize metadata storage and processing. Although much of our pipeline can be tested in silico, we lack a simulator for photomanipulation. In silico testing is essential for troubleshooting and the implementation of automated testing across our pipelines.

## Outcome-driven microscopy to guide cell biology

**Alfredo Rates, Josiah B Passmore, Lukas C Kapiten**, Utrecht University

### Outcome-driven microscopy

Smart microscopes take microscopy automation to the next level by adapting and optimizing the acquisition **based on the experiment in progress**. Thanks to information about the sample, either by user input, previous experiments, or defined strategies, with smart microscopy it is possible to get as much information as possible, while reducing sample damage. But smart microscopy is not limited to observation, it can also be expanded to have **control over the biology** thanks to **outcome-driven microscopy**. Outcome-driven microscopy is a smart microscopy strategy where the automation of the microscope is not only to change the acquisition parameters, but also to interact with the sample using the microscope hardware. As it is known, parameters such as temperature, flow, illumination power or color are inputs that affect biological samples. If this effect is controlled and known, it can be used to **drive the biological processes to desired states**. Indeed, this is possible with optogenetics. In optogenetics, photo-controllable proteins are used to activate processes at the subcellular level when illuminated with particular wavelengths. These proteins are not only sensitive to wavelength but also to light intensity and spatial patterning. Thus, by controlling these parameters, **optogenetics has the power to precisely control the biology of cells**.

#### Modular Python platform for smart microscopy

To achieve outcome-driven microscopy, we combine smart microscopy with optogenetics, changing the optogenetic signal on the fly, and having a feedback loop between the (cell) biology and the (microscope) hardware. For this, we developed a modular software architecture in Python.

**Fig. 1.**
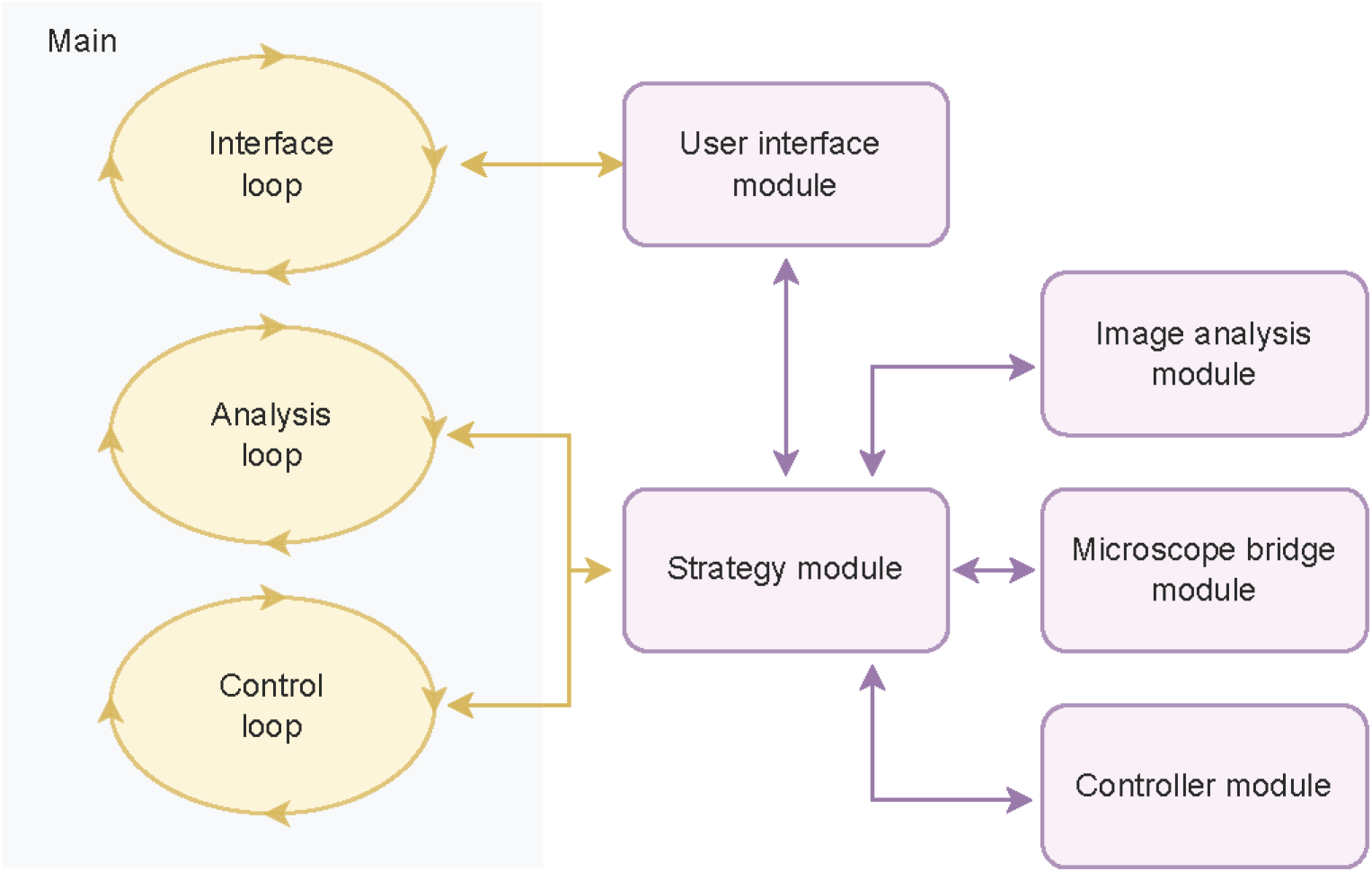
**Modules workflow**, including the main.py script and its three internal parallel loops. The external modules in purple are imported as external libraries. Arrows represent communication lines between modules. Figure taken from [4] with permission of the authors.

We developed a Python-based platform to perform automated, outcome-driven experiments. Our platform is organized into multiple modules, as described in Fig. 1. The platform has specific modules for user interface, image analysis (e.g., cell segmentation [1]), control strategy (e.g., PID controller [3]), microscope communication (for specific hardware, e.g., Micromanager or Zen Blue Zen Blue), and outcome-driven strategy, where the algorithm for the specific experiment is defined (e.g., control cell expression to certain levels [2]). The user interacts with the platform through a `main.py` script and a graphical user interface (GUI). The main connects to the rest of the modules, including the GUI (see Fig. 2), and runs three parallel, asynchronous loops. The three loops consist of the interface loop, the analysis loop, and the *control loop*.

**Fig. 2.**
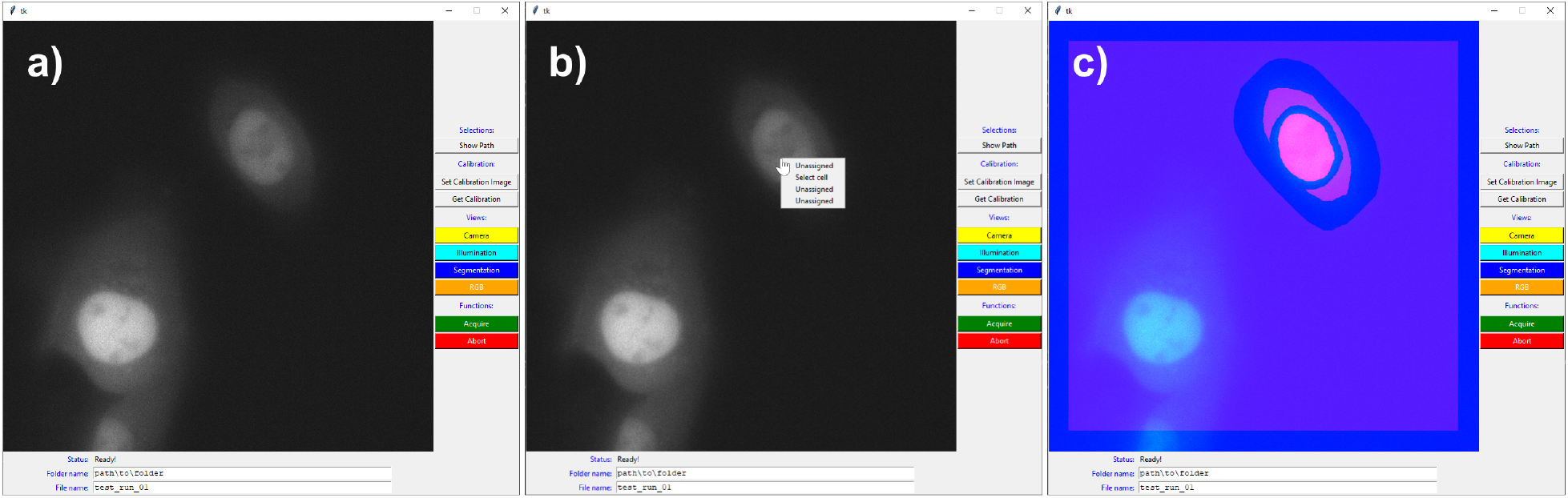
**Graphical user interface** in three different states. **A)** when the software is just initialized, **B)** when right-clicking the cell to segment, including the drop menu, and **C)** the RGB visualization showing camera image, cell segmentation, and illumination pattern. In this case, the segmentation separated the cytosol from the nucleus, adding a margin in between. Figure taken from [4] with permission of the authors.

The modules are imported to the main as external libraries based on the user input. Each module is structured as a class with an abstract class as a parent, meaning that they must have a list of functions and variables in order to communicate correctly with the rest of the platform. In case the user needs to develop a custom module, they can easily do it as long as the custom module includes these functions and variables. When running an experiment, the platform exports 4 main results. The first result is the time series acquisition, including channels and z-stacks, if previously defined. In addition, the platform exports the cell segmentation for each time step, the controller data (e.g., error, setpoint, laser power) as a .csv file, and the metadata of the experiment as a .json file.

#### Demonstration of outcome-driven microscopy using optogenetics

To demonstrate the power of outcome-driven microscopy, we applied our platform to directed single-cell migration, using an optogenetic construct to recruit the RAC1 effector TIAM1 to the plasma membrane. With outcome-driven microscopy, we adapt on the fly the area of the illumination within the cell, changing where the TIAM1 is recruited and thus drive migration in a specific direction. We change the illumination area of the optogenetic light using a digital micromirror device (DMD). For this implementation, the control module is based on a trajectory tracking controller based on the cell centroid relative to a path made up of setpoints, and the image analysis module uses the pre-trained AI segmentation method Segment Anything Model (SAM) [1] with a custom extension of its interface to track the cell. Thanks to our stable cell line and the outcome-driven platform, we could reliably guide cells around a specific path for many hours, maintaining a relatively consistent speed. Fig. 3 shows the cell migration path over the defined circle. This experiment was carried out in a Nikon Ti dual-turret inverted microscope and a Mightex Polygon 400 DMD, all controlled via Micromanager.

**Fig. 3.**
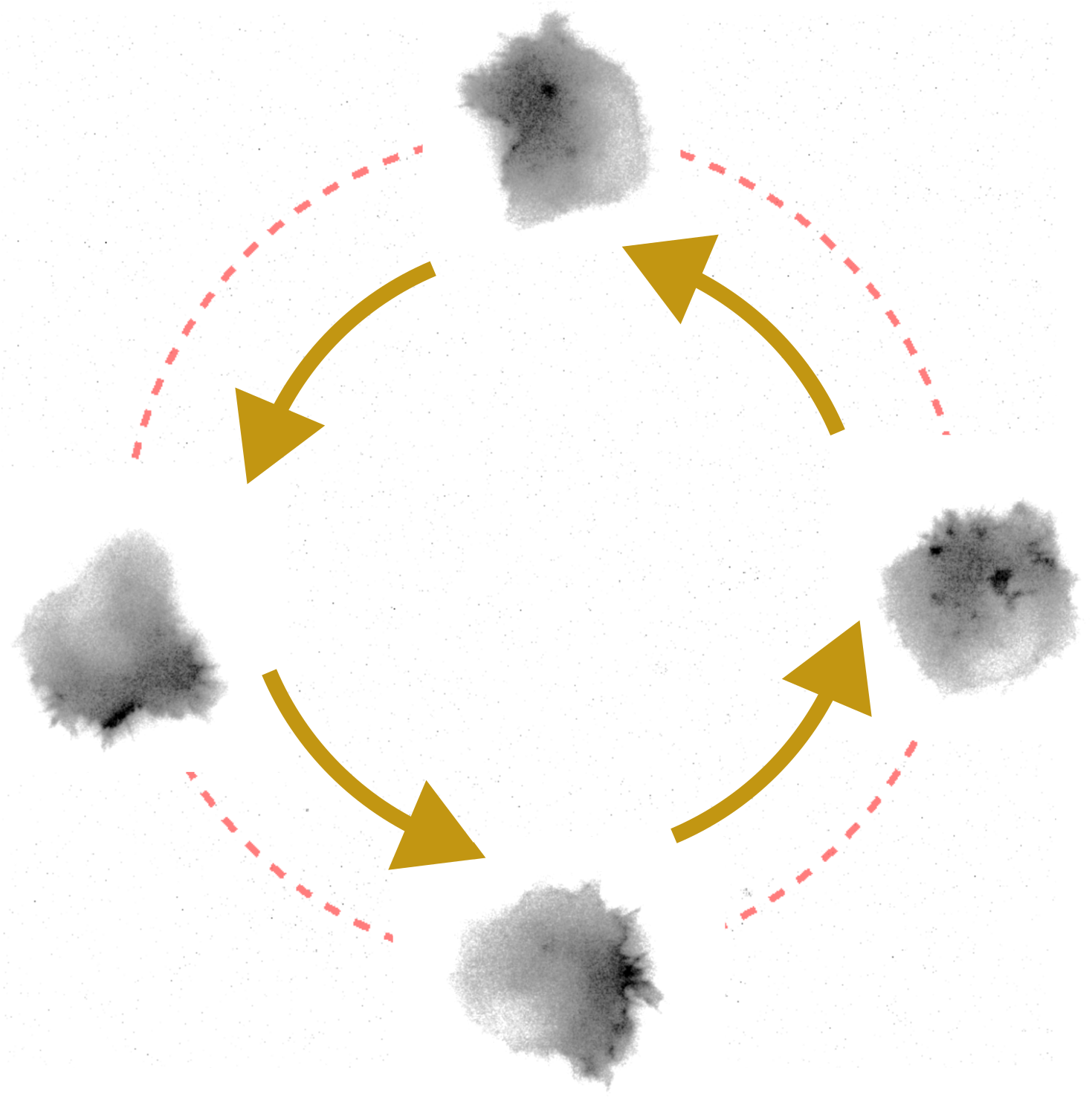
HT1080 cell migrating in a circular path due to blue light stimulus. The area of illumination was constantly adapted using our outcome-driven microscopy platform. This measurement was performed in a Nikon Ti using a Mightex Polygon 400 for activation.

Thanks to our modular approach for the microscope bridge, we were able to adapt our communication scheme from a Micromanager-controlled microscope to a Zeiss microscope controlled by ZenBlue. We used an LSM-980 with AiryScan2 module, and instead of modulating the optogenetic light with a DMD, we adapted the region of interest (ROIs) of the confocal scanning. The results of the confocal-based cell migration is shown in

**Fig. 4.**
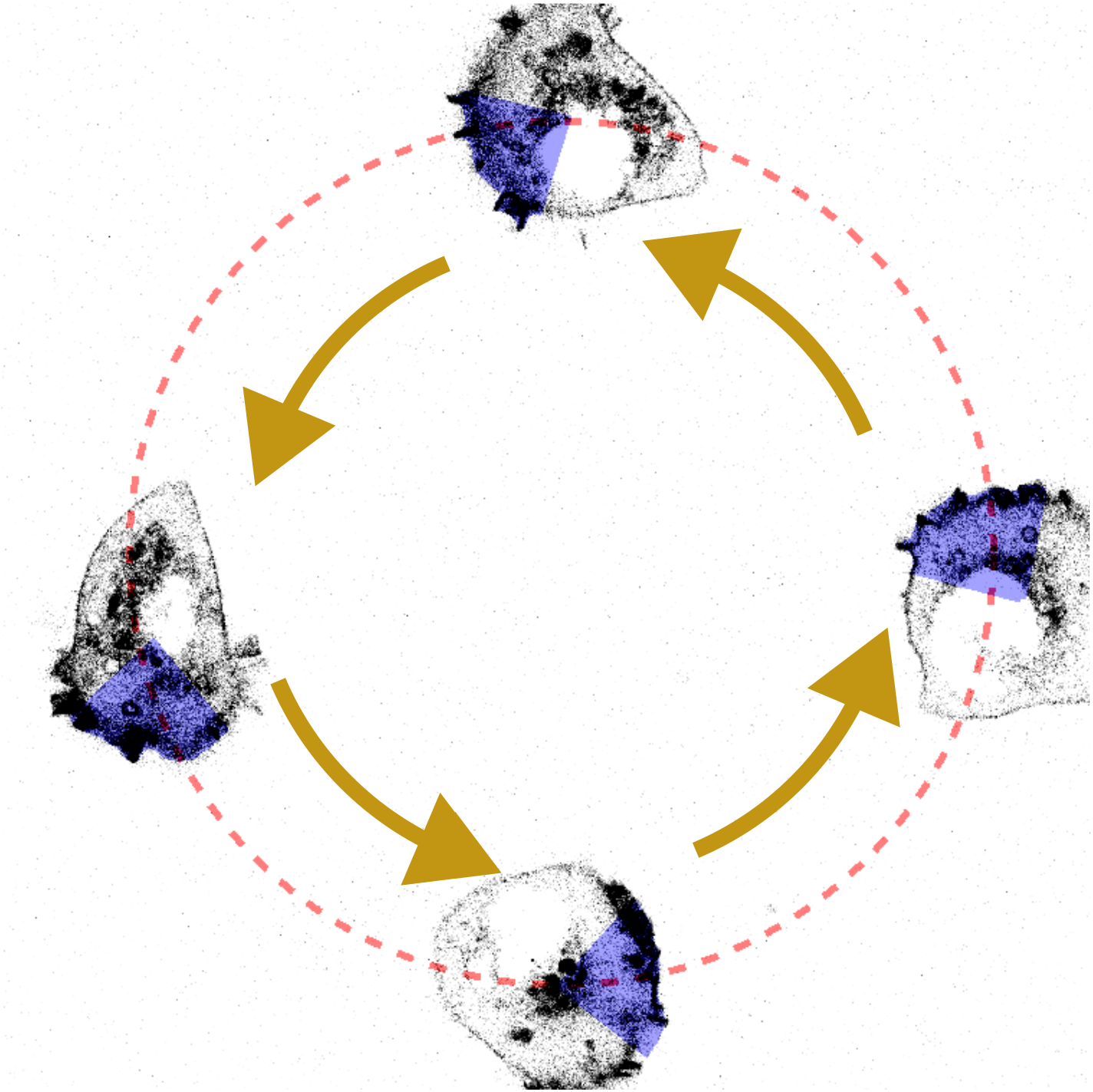
HT1080 cell migrating in a circular path due to blue light stimulus, same as Fig. 3. This time, the measurement was performed in a Zeiss LSM980 and the activation was done scanning the area with the built-in blue light laser.

### Limitations and outlook

The advantage of modularity is not only that modules are exchangeable, but also that the platform is easily upgraded and improved. Among the improvements planned to the platform, we first intend to **separate further the bridge module with the strategy module**. In particular, the platform is structured around the multi-dimensional acquisition process of Micromanager. This structure is ideal for our outcome-driven experiments and can be translated to other microscopes easily, but it limits experiments to having a constant acquisition configuration, such as time step and number of Z-stack layers. To make our platform flexible, we plan to follow the structure offered by the *pymmcore-plus* library. *pymmcore-plus* is an alternative to Pycromanager to control Micromanager-compatible hardware, and is based around a **queue of events** instead of the multi-dimensional acquisition, i.e., a list of orders to give to the microscope. Furthermore, we plan to **make a new interface based** on the *Napari* [5] library. Napari is an extensive visualization library, widely used for microscope data. This new interface offers more functionalities and interaction with the user, and it is also compatible with *pymmcore-plus*.

## Modular adaptive feedback microscopy workflows with AutoMicTools

**Aliaksandr Halavatyi, Manuel Gunkel, Rainer Pepperkok**, Advanced Light Microscopy Facility (ALMF), European Molecular Biology Laboratory (EMBL), Heidelberg Germany

### Specific Focus and scientific questions asked

Adaptive Feedback Microscopy is used to automate and run different kinds of microscopy experiments with integrated image analysis in a high throughput manner for example detection and high-resolution imaging of certain phenotypes or different kinds of photomanipulation experiments. Applying this methodology for various kinds of experiments often requires dedicated hardware. Many systems, for example confocal laser scanning microscopes (cLSMs), are controlled by vendor proprietary software. Therefore, microscope companies need to design dedicated modules for their softwares for running smart microscopy workflows. If different image analysis and workflow management tools are used on different microscope types to automate them, microscopy facilities and other research infrastructures operating microscopes of different brands need to support many additional software tools to benefit from available hardware. Migration of workflows implemented on different microscopes becomes hardly possible, even if needed imaging modalities are available. To overcome these limitations, we established a toolbox called *AutoMicTools* which enables setting up, testing and running adaptive feedback microscopy workflows on several advanced systems. The implemented workflows consist of multiple interacting modules to separate image analysis routines and logics for communicating to the microscope.

**Fig. 1:**
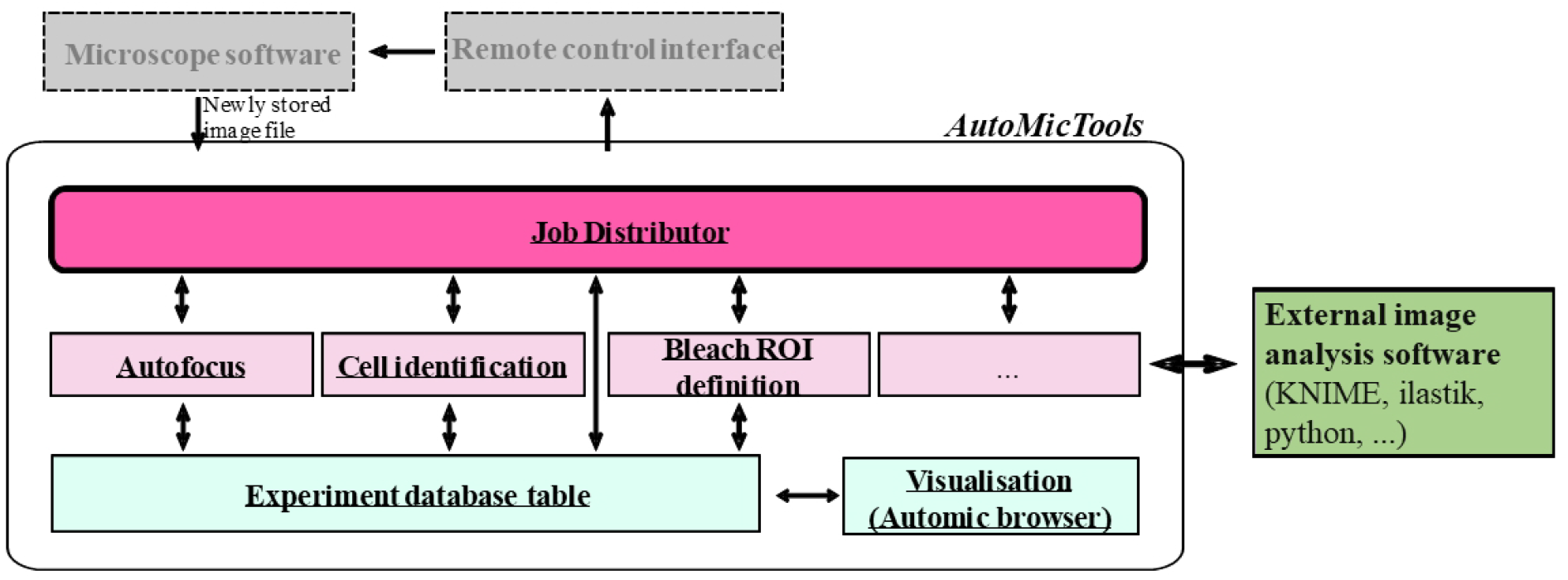
Structure of *AutoMicTools* workflow and its interaction with external software tools.

#### Key findings and innovations

Several workflows have been designed with *AutoMicTools* toolbox and run on different confocal microscopes in our facility:

##### High resolution multicolour volumetric imaging of classified phenotypes and rare events

Several versions of this workflow have been established in the context of cell biology and developmental biology studies. For example, live cell cultures [5], [1], have been imaged to identify and image cells in a particular stage of a cell cycle for a time period up to 72 hours. A screen of drosophila stained lines has been performed to collect high resolution data of gene expression patterns in those samples [4][3]. All projects have benefitted from the possibility to extend the workflows by integrating multiple image acquisition and analysis steps on demand. For example, an additional round of focusing has been implemented to estimate axial positions of individual cells or embryos after identifying them in 2D and automatically adjust high resolution imaging volume.

##### Automated photomanipulation

1. Fluorescence Recovery After Photobleaching (FRAP) experiments have been run automatically to measure turnover of COPII proteins at ER-exit sites (ERES). More than 500 recovery curves have been collected in each imaging session (∼16 hours long), allowing to study distributions of recovery rates and correlate them with other measured parameters of target cells.
2. Automated photo-activation has been set to enable researchers to automatically identify, fluorescently mark, sort and sequence single cells or cell populations carrying target phenotypes. The workflow has been initially established in the framework of a project aiming to study networks regulating Golgi structure [7]. Cells under well-defined gene knock-down conditions show heterogeneous Golgi morphology phenotypes suggesting that the different phenotypes are due to different molecular stages the cells are in. In order to address this question, it has been necessary to directly correlate the Golgi morphology phenotype with the molecular stage of the cells. The originally established protocol has been used for studying other biological phenomena by plugging in project-specific image analysis modules identifying target cells [2].

#### Methodology and implementation details

A core microscope-agnostic part of the workflow is implemented as the *AutoMicTools* library for ImageJ/Fiji containing multiple plugins. Several imaging and photomanipulation workflows are available in *AutoMicTools* as ready-to-use plugins, new workflows can be created by subclassing provided Job and JobDistributor classes using Java programming or Fiji scripting tools (Fig. 1):

##### Job

An *AutoMicTools* Job needs to be defined for each particular image analysis task. It combines functions for image analysis itself, pre- and post-processing, on-the-fly visualisation of analysis results and creating feedback commands in a microscope-agnostic manner. Project-specific analysis Jobs can be created by subclassing the Job_default base class using Java programming or Jython Scripting using provided examples as templates. Alternatively, predefined jobs accepting user-defined image analysis macros or scripts as parameters can be used. Jobs can be modified to to bridge to external applications for running image analysis (e.g. Python, Ilastik or KNIME).

##### JobDistributor

The JobDistributor module is responsible for monitoring a defined folder for the arrival of new files and - based on their name tags - executing predefined image processing tasks defined by the Job classes. An individual JobDistributor with an arbitrary number of Jobs has to be configured for each workflow by mapping the names of the image file tags to Analysis Jobs to be executed.

#### Contributions to Interoperability

*AutoMicTools* Fiji plugins allow to set up and debug microscope-agnostic parts of the workflow. The workflow can be simulated without access to the microscope when example images for different imaging jobs are available. The same core workflow can be run on different microscopes by using brand-specific communication tools, implemented using corresponding API layers:

##### Zeiss ZenBlue

We set up multiple *AutoMicTools*-based imaging and photomanipulation feedback microscopy workflows using Zeiss confocal microscopes, in particular LSM900 and LSM980, controlled by ZenBlue. For that we make use of the ZenBlue macro programming environment which is part of the Open Application Development (OAD) package. Established macros control acquisition loop, reuse and trigger a priori defined imaging settings based on the feedback received from image analysis [code repo].

##### Zeiss ZenBlack

Multiple of the above mentioned workflows we also run on the previous generation of Zeiss confocals (LSM780/880). Macro programming in this software is only possible with Visual Basic (VB), which is not commonly used these days. To link *AutoMicTools* to ZenBlack, we make use of the MyPic macro developed by Antonio Politi in the Ellenberg lab current repository [6]. MyPic has a friendly GUI to keep track of multiple acquisition/photomanipulation settings, and to define default positions; it is capable to receive a feedback from an external application (e.g. *AutoMicTools*) with computed imaging positions of photomanipulation ROIS via the Windows registry.

##### Evident(Olympus) FluoView

Software driving FluoView3000 and FluoView 4000 point scanning confocal microscopes, is equipped with an RDK module, which allows to change acquisition parameters and control some of the hardware components (e.g. stage, focus drive and objective turret) by sending XML-formatted commands from different programing languages. We have developed a Fiji plugin which makes use of this functionality and controls Acquisition settings and sample positions in *AutoMicTools* workflows. Settings for each imaging job on the microscope are provided from a configuration file.

##### Nikon NIS Elements

The Nikon software NIS-Elements provides also a macro environment. We have written a set of macros which, when executed, wait for a specific text file containing information which experiment to run. ROIs are transferred as binary tif files via the NikonCommander in automic tools. The workflow is controlled by an additional jython script. Leica LMD (Leica Laser Microdissection 7) An image produced by the LMD system (in *.bmp format) is analysed and stored in a new path. The resulting ROIs are stored in an xml file that can be parsed by the microscope software. For synchronization a small executable is needed which is called by the LMD software and closed by automic tools. The close of the exe triggers the LMD to proceed with the next position [code repo].

#### Limitations

Although the implemented schema has a lot of flexibility for customising and extending the workflows for project-specific needs, it has several restrictions linked to the limitations of APIs that are provided by vendors for their proprietary software. Different APIs support different functionalities, which makes generalization challenging. For example, the transfer of image data from acquisition software to our Fiji plugins currently requires saving image files, as in majority of use cases API to transfer image data in memory is currently not available. This aspect can be important for time critical applications, such as tracking of fast live samples by moving the acquisition field-of-view. Likewise, certain essential features, such as specifying arbitrary shapes of photomanipulaiton ROIs, rotation of the field of view, changing scanning area for LSM systems and changing the z-stack height, are not available for many vendor APIs: We collaborate with our industry partners in several formats, including Smart Microscopy Working Group, to enhance the power of provided APIs, that will contribute to flexibility of *AutoMicTools* and other packages making use of them for microscope automation.

## Smart Microscopy as a Service

**Rafael Camacho**, Centre for Cellular Imaging, University of Gothenburg

### Approach to smart microscopy

At the Centre for Cellular Imaging (CCI), Smart Microscopy is developed within a fee-for-service model, where users are trained to operate commercially available microscopy equipment from vendors such as ZEISS and Thermo Fisher. This approach prioritizes accessibility and reproducibility, ensuring that automation strategies remain compatible with vendor-provided microscope control software. Smart Microscopy at CCI is implemented with a holistic perspective, addressing not only adaptive feedback microscopy—where acquisition parameters are adjusted dynamically based on real-time sample analysis—but also downstream processes such as image processing, quality control monitoring, and data management. By considering the entire imaging pipeline from acquisition to analysis and integration, we aim to enhance efficiency, reproducibility, and scalability within a research infrastructure setting.

#### Methodology, Implementation details

At CCI, we implement Smart Microscopy through automated feedback loops that connect image analysis with hardware control. Our workflows leverage machine learning-based object detection and segmentation to enable real-time adaptive imaging, adjusting acquisition parameters based on sample characteristics.

1. Light Microscopy, vendor ZEISS We utilize Open Application Development (OAD), a framework within ZEISS ZEN Blue that integrates an Iron Python script editor to automate microscopy workflows. OAD enables responsive imaging strategies, processing real-time image analysis cues from both internal algorithms (running within ZEN) and external tools such as Python scripts and ImageJ/FIJI plugins. Our primary efforts have focused on:
  - **Targeted Imaging**: Adaptive imaging routines that prioritize specific sample features.
  - **Object Tracking**: Automated tracking of dynamic structures in live-cell imaging applications.
2. Electron Microscopy (ZEISS SEM & Thermo Fisher TEM) ZEISS SEM systems lack an efficient API for direct hardware communication (an issue currently being addressed). As a result, we have concentrated on:
  - **On-the-Fly Image Analysis and Quality Control**: Implementing real-time image evaluation workflows to detect acquisition anomalies and improve data reliability.
  - **Data Standardization**: Developing pipelines for converting complex vendor-specific image data structures (e.g., multi-image projects with proprietary metadata) into OME-Zarr, an open, cloud-compatible image format. For TEM workflows, we emphasize data format standardization to ensure interoperability with downstream analysis tools, particularly through the adoption of OME-Zarr.
3. Data Management Infrastructure As of early 2025, CCI has deployed an **OMERO-based image data management system** that streamlines data flow across all facility microscopes. This infrastructure facilitates seamless data transfer from acquisition systems to computational analysis servers, and supports integration with users’ own storage solutions, working towards ensuring FAIR (Findable, Accessible, Interoperable, Reusable) data principles.

#### Key features and innovations

The Smart Microscopy implementation at CCI integrates advanced automation strategies across diverse imaging modalities, ensuring seamless operation within a fee-for-service core facility model. Our key innovations focus on adaptive imaging, interoperability, and data standardization, enhancing both image acquisition and post-processing workflows to improve efficiency and reproducibility. Adaptive imaging and automated feedback loops enhance real-time decision-making, enabling more precise and efficient imaging workflows. Targeted imaging on light microscopes leverages machine learning-based object detection within ZEISS ZEN Blue’s Open Application Development (OAD) framework, dynamically selecting regions of interest (ROI) based on sample characteristics. This also facilitates object tracking for live-cell studies, though current research is focused on improving feedback speed for faster adaptive responses. In electron microscopy, on-the-fly image analysis for SEM workflows enables real-time quality assessment and anomaly detection, optimizing acquisition efficiency in high-throughput environments. These innovations create a more responsive, automated, and reproducible microscopy workflow, reducing manual intervention while enhancing data quality and throughput. CCI’s Smart Microscopy implementation also enhances interoperability and scalability, ensuring seamless integration across imaging modalities and research infrastructures. Standardized data conversion to OME-Zarr facilitates compatibility across light, SEM, and TEM workflows, ensuring structured data handling and longterm accessibility. Multi-software integration maintains flexibility, enabling workflows that connect ZEISS ZEN, FIJI/ImageJ, and Python-based analysis tools while preserving vendor compatibility. The deployment of an OMERO-based data management system further supports automated image transfer, metadata structuring, and remote access, streamlining data handling from acquisition to analysis. Designed for **open-access imaging facilities and research environments**, CCI’s Smart Microscopy solutions lower the technical barrier for advanced imaging automation, making cutting-edge workflows more accessible to a broader user base. The infrastructure is optimized for scalability and interoperability, allowing workflows to be shared, adapted, and deployed across multiple research institutions.

#### Contributions that could contribute to interoperability

All Smart Microscopy developments at CCI are designed with open software and interoperability as core principles, ensuring seamless integration across imaging platforms and analysis tools. By adopting standardized data formats such as OME-Zarr and implementing OMERO-based data management, we facilitate structured, vendor-independent data handling. Our multi-software integration strategy ensures compatibility with ZEISS ZEN, FIJI/ImageJ, and Python-based analysis workflows, enabling flexible automation while maintaining vendor compatibility. These efforts lay the foundation for scalable, shareable, and adaptable imaging workflows, with interoperability remaining a key focus for future developments (see roadmap for details).

#### Current bottlenecks, Roadmap

While CCI’s Smart Microscopy implementation has made significant progress in automation, interoperability, and data standardization, several challenges remain in fully realizing the potential of adaptive microscopy workflows. One of the main bottlenecks is limited access to vendor APIs, particularly for electron microscopy systems. While ZEISS light microscopes support automation through the Open Application Development (OAD) framework, EM platforms lack efficient software interfaces for direct hardware communication. This limitation prevents real-time adaptive feedback loops, requiring workarounds such as post-acquisition quality control rather than live adjustments. We are actively collaborating with industry partners to explore solutions that enable deeper automation and integration within these systems. Another challenge is the speed and efficiency of real-time feedback microscopy, particularly in live-cell imaging. While targeted imaging and object tracking are already implemented, response times remain constrained by latency in processing real-time image analysis outputs and sending commands back to the microscope. Further optimization of computational workflows and hardware-software communication is needed to improve the responsiveness of adaptive imaging, ensuring that real-time decision-making can keep pace with dynamic biological processes. In terms of data management and interoperability, we have successfully deployed an OMERO-based infrastructure, but ensuring seamless integration across different imaging modalities and user storage solutions remains an ongoing effort. The conversion of complex, vendor-specific data structures into standardized formats such as OME-Zarr is progressing, but further development is needed to refine metadata extraction, enhance compatibility with emerging image analysis pipelines, streamline large-scale data transfers, and integrate computational resources such as national CPU and GPU clusters for scalable analysis. Looking ahead, our roadmap prioritizes the development of a Python-based Smart Microscopy Workflow Manager, designed to act as a user-level interface that allows researchers to define imaging workflows independent of the underlying microscope hardware. This system will abstract communication with different microscopes—even from different vendors—ensuring a unified approach to controlling diverse imaging platforms. Additionally, it will provide flexibility in image analysis execution, seamlessly integrating processing across Java, Python, or proprietary software environments, enabling users to focus on scientific insights rather than software compatibility. By addressing these challenges, we aim to make Smart Microscopy at CCI more robust, accessible, and scalable, ensuring that our workflows continue to support cutting-edge research in imaging sciences while paving the way for broader adoption across research infrastructures.

## CelFDrive: Multimodal AI-Assisted Microscopy for Detection of Rare Events

**Scott Brooks [1,2], Sara Toral-Pérez [1,2], David S. Corcoran [1,2], Nina Pučeková [1,2], Karl Kilborn [3], Brian Bodensteiner [3], Hella Baumann [3], Nigel J. Burroughs [5,2], Andrew D. McAinsh [1,2], Till Bretschneider [6,2]**

[1]: Warwick Biomedical Sciences, Warwick Medical School, University of Warwick, Coventry, CV47AL [2]: Centre for Mechanochemical Cell Biology, University of Warwick, Coventry, United Kingdom

[3]: Intelligent Imaging Innovations, Denver, Colorado, United States of America [4]: Intelligent Imaging Innovations, London, United Kingdom

[5]: Zeeman Institute (SBIDER), University of Warwick, Coventry, United Kingdom [6]: Department of Computer Science, University of Warwick, Coventry, CV47AL

### Specific Focus and scientific questions asked

The crucial physical process during mitosis is the segregation of the duplicated chromosomes into daughter cells. This depends on the kinetochore, a dynamic, multi-protein complex that assembles on the centromere of each chromosome and forms attachments to the mitotic spindle. Advances in live cell imaging and computational analysis now permit the quantitative study of kinetochore dynamics during mitosis. This is a powerful approach to understanding how and when kinetochores move chromosomes, and what determines whether they are moved to the correct daughter cell [3]. However, a barrier to investigating kinetochore dynamics throughout mitosis is identifying cells that have not yet broken down the nuclear envelope and initiated key events i.e. spindle assembly. Finding such prophase cells is hard as they are 1) rare in a normally cycling population, and 2) can often be difficult to distinguish from interphase cells.

Lattice Light Sheet Microscopy (LLSM) is a state-of-the-art technique for acquiring 3D volumes of delicate biological processes over extended periods at high temporal resolution in living cells. LLSM makes it possible to capture kinetochore dynamics from prophase to anaphase with low phototoxicity and minimal photobleaching. However, LLSM is limited to a small field of view, which can make it difficult to find rare cells of interest. Moreover, even when a cell of interest is identified, determining its exact phase can be challenging, potentially leading to wasted time and missed fleeting events. Often, this leads to large storage of redundant data and inefficient use of researcher time, preventing the generation of the large datasets we require to produce reliable quantitative insights.

Smart microscopy has the power to address this “needle in a haystack” problem by rapidly detecting rare cells, minimising redundant data and accelerating high resolution imaging workflows.

#### Key findings and innovations

CelFDrive is an integrated artificial intelligence-driven platform for automated high-resolution 3D imaging across various fluorescence microscopes. It overcomes the limitations of manual imaging by applying deep learning-based cell classification to low-magnification images, converting image-based detections into precise stage coordinates. This facilitates automated repositioning for subsequent high-resolution 3D imaging modalities. CelFDrive accelerates the detection of rare events in large cell populations, such as the onset of cell division. In our example we achieve an acceleration of 34 times compared to a human expert.

**Fig. 1:**
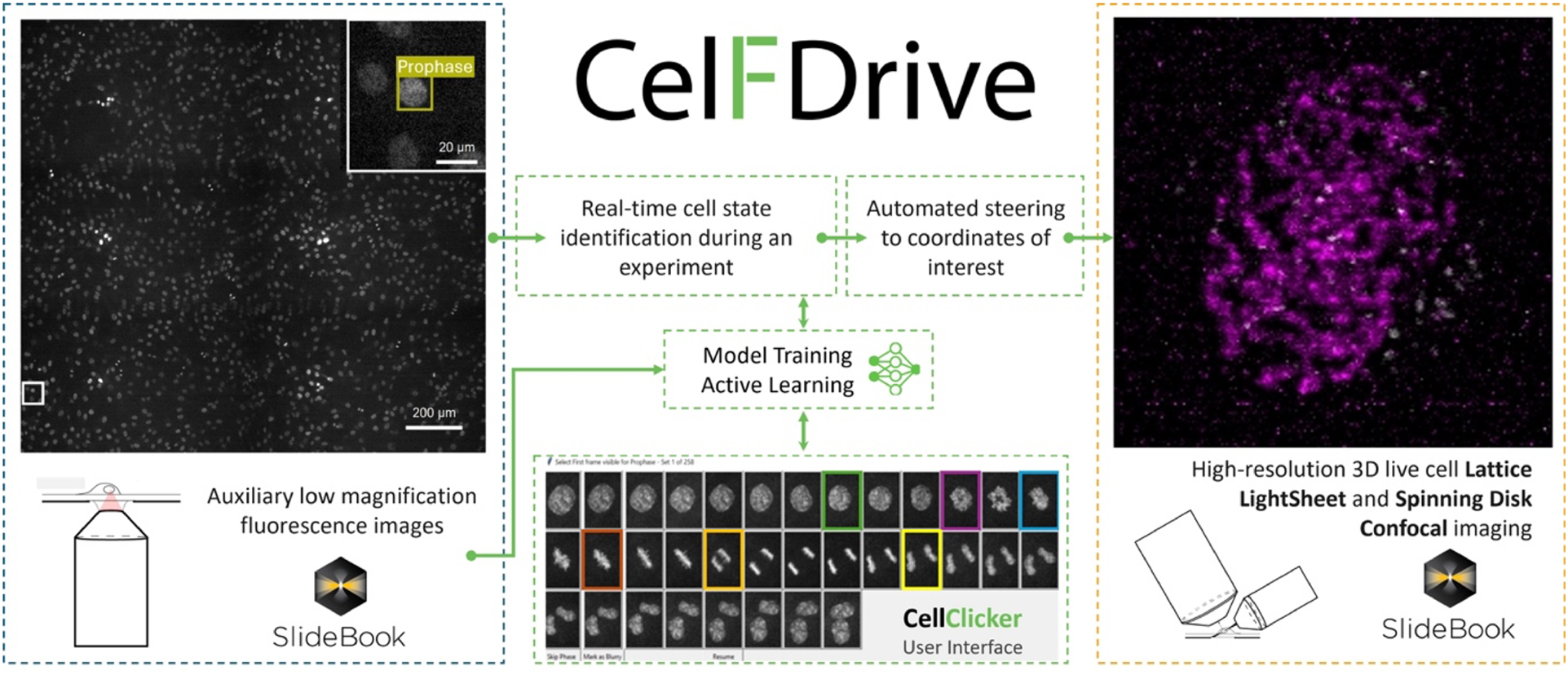
CelFDrive overview: The flow of data through various components of the CelFDrive. Initial search region is captured via a low magnification montage. Cells of interest are detected by a deep learning model which has been trained on time-lapse movies from the low magnification objective. Data annotation is performed using the CellClicker and CellSelector interfaces. Image coordinates are converted to physical stage locations and high-resolution 4D time-lapses are taken at the region(s) detected.

Using CelFDrive we have been able to capture vast 4D timelapse datasets with LLSM that depicts the loading of endogenous NDC80-EGFP (a kinetochore marker) onto the centromeres alongside secondary markers for the chromosomes, mitotic spindle and/or nuclear envelope. From this we use our Kinetochore Tracking software (KiT) [1] to gain insight into kinetochore dynamics at the earliest moments of mitosis at higher temporal resolution than previously possible. These image sequences are providing new insights: 1) We revealed the relocation of TMR:HALO-NUP107 (a nucleoporin) from the nuclear envelope onto the kinetochores. This is the first reported live imaging data that captures this event directly. 2) We found that the kinetochores’ outer plate (NDC80-EGFP) is assembled before the NUP107 translocation. Thus, efficient identification of late prophase cells with CelFDrive allowed capture of the data necessary to order molecular events during early mitosis (Outer plate assembly -> completion of nuclear breakdown -> translocation of Nucleoporins to kinetochores). 3) We successfully used a multimodal model which supports spinning disk and widefield systems as a high throughput solution to quantify levels of expression of secondary markers within dividing cells across a whole coverslip.

The broader implications of this work extend beyond the study of mitosis to diverse dynamic processes and rare events at both cell and tissue scales for potential drug discovery applications and clinical diagnostics. Our method highlights the potential of integrating artificial intelligence with modern microscopy techniques

#### Methodology and implementation details

CelFDrive interacts with our industry partner’s (3i) software, SlideBook. SlideBook’s Conditional Capture module automates imaging workflows by triggering macro scripts directly after image capture. In the current workflow, a large field of view is imaged with an auxiliary objective and coordinates are transferred from SlideBook to Python. From here an object detection model is applied in parallel to process patches of images and determine classes and coordinates. Images are translated into physical locations using pixel spacing and an object’s image position, and if specified, a coverslip tilt is incorporated to correct the Z position. SlideBook capture scripts enable automated multi-point acquisition of cells detected in a specific phase of interest, allow the user to select cells from a list of top hits, or a user can specify that they wish to just image the top confidence cell with the most desirable class (e.g. a prophase cell).

We developed a multimodal workflow that can be easily installed on a variety of 3i microscopes (Lattice Lightsheet, Marianas spinning disk confocal, widefield systems) for the detection of cells in different stages of mitosis. By training a model on different modalities, stains and magnifications we can deploy the same model to different microscopes with little to no retraining. In addition, innovative python-based user interfaces, including CellClicker and CellSelector, were developed to support efficient multi-user annotation of time series data, thereby improving the robustness of the training sets and reducing subjectivity. These interfaces allow users to apply CelFDrive to their unique imaging tasks, providing an adaptable and scalable solution. We provide an anaconda environment file to assist users with installation of the required dependencies.

To detect cells of interest, we fine-tuned state-of-the-art real-time object detection models on diverse live image datasets of DNA-stained mitotic cells acquired using various magnifications of widefield microscopy and spinning disk confocal microscopy. The model architectures used are from the YOLO family and it is natively compatible from versions 5 to 12 [2] as well as real-time detection transformer (RT-DETR) [4], however, it is easy to adapt the software to different object detections models. The resulting generalisable model facilitates high-precision smart microscopy which is cross-platform and cross-magnification, one of the first solutions of its kind presented for live fluorescence microscopy.

#### Contributions that could contribute to interoperability

This work is free to use for academic research. The pythonic nature of CelFDrive means that it can be integrated with any microscope that can trigger python commands. In addition, for the case of cell division, the model trained can be applied to most DNA-stained images of cells at different magnifications. This model can be applied to other smart microscopy methods or used as a post-acquisition analysis tool.

#### Bottlenecks / limitations of approach, specific requests / suggestions to community and industry

The primary limitation of this approach is that a supervised model has been used, this means that to train a model on a new biological use case, a large quantity of data is required to produce a multimodal model. Although for a specific use case on a specific microscope, this data requirement is much lower depending on the process to be detected.

In addition, CelFDrive has been optimised to work with 3i-systems Synergy module (python-based) approach, where image data and coordinate data are defined in a specific convention which may not be applicable to other systems. This may require an intermediary script to, for example, re-order image multidimensional data, convert coordinates and monitor watch file systems. This poses a question about standardisation of industry and academic microscope control software, where defining a common data structure format for smart microscopy API integration would allow for ease of implementation of different techniques across platforms and increased uptake of smart microscopy.

## The EnderScope as an educational platform for Smart Microscopy

**Erwan Grandgirard** (IGBMC: Illkirch-Graffenstaden, France), **Jerome Mutterer** (CNRS, Strasbourg University, Strasbourg, France)

### Specific Focus and scientific questions asked

The smart microscopy field is rapidly expanding and users are expected to appropriate its base concepts to take full advantage of the available new imaging approaches. Commercial often exemplify classical smart workflows which can limit the potential applications that users can envision. Moreover, practical learning about smart microscopy can be hindered by the current low number of available systems, or by costly access fees when highend devices actually are available. As discussed in the associated article, a further barrier is that developing smart microscopy workflows often requires learning at least some notions of programming or algorithmics, device control, image acquisition and image processing principles. The EnderScope with its matching control library aims to address these issues by providing a low cost, easy to learn ecosystem to teach, learn and develop smart microscopy [1].

#### Methodology, Implementation details Hardware setup

The hardware setup of the EnderScope is built around the frame of an cartesian 3D printer, which provides a reliable and precise 3-axis motorized stage for positioning. A Raspberry Pi serves as the central computational unit, coordinating the system and hosting the control software. Imaging is performed using interchangeable Raspberry Pi camera modules, ranging from the standard Camera Module v2 to the High Quality Camera and even third-party options such as the Arducam 64 MP autofocus camera, allowing flexibility in resolution, optics, and field of view. Other cameras also are an option, provided they have a device driver for RaspberryPi. Illumination is delivered by an array of RGB LEDs controlled by an Arduino microcontroller, which replaces the earlier manual switch with programmable lighting modes and adjustable intensity. All components communicate via serial over USB or the Pi’s camera connector, creating a compact, low-cost, and customizable imaging platform that can be assembled from widely available, low cost, parts.

#### Software library

The `enderscope.py` software library provides the core interface that makes the EnderScope both accessible and versatile. Written in Python, it abstracts low-level device control into intuitive, high-level functions for the stage, illumination, and camera. The library includes dedicated classes such as Stage for precise axis movements using G-code, Enderlights for programmable multi-color LED illumination, and helper tools like SerialDevice for seamless USB communication. Synchronization mechanisms ensure that commands—such as moving the stage, setting illumination, and capturing images—execute in the proper sequence, preventing motion blur or misaligned acquisitions. Beyond code, the library also integrates simple graphical interfaces via iPywidgets, giving users manual control for exploration and setup. By combining structured programmatic access with hands-on examples and Jupyter notebooks, `enderscope.py` enables both beginners and advanced users to design, test, and expand smart microscopy workflows with ease.

#### Contributions to Interoperability

Using Python for the EnderScope library matches the dominant language in smart microscopy. Most imaging tools such as scikit-image, Napari, and Micro-Manager bindings, are Python-based, ensuring easy integration and interoperability. Its simplicity lowers the entry barrier for beginners while offering advanced capabilities for researchers, making the system both accessible and powerful. To facilitate user onboarding, examples are provided for tasks of basic or intermediate complexity to programmatically control all system elements, or perform classical computational imaging tasks. This includes: Stage movements along 3 axis; Illumination settings in various modes; Image capture using the Raspberry camera; Autofocus (using Laplacian variance on a Z-stack); Shadow/reflection removal with multi-angle illumination; SmartScan: overview scan + optimized region-of-interest imaging using TSP optimization. Scripts developed using the `enderscope.py` library can easily be translated to other smart microscopy APIs that typically provide identical or similar objects and associated functions.

#### Limitations

- Mechanical precision: The used printer stage, while low-cost, lacks the nanometer accuracy and stability of professional microscope stages, limiting resolution and repeatability.
- Optical constraints: Standard Raspberry Pi or Arducam cameras and simple optics cannot match the sensitivity, dynamic range, or optical flexibility of scientific-grade cameras.
- Illumination: Although the LED array enables multi-angle and color variation, it cannot provide the coherence, polarization, or spectral specificity required for advanced fluorescence techniques.
- Speed and synchronization: Stepper-driven stage movements are relatively slow, and command execution through serial communication can introduce latency compared to dedicated hardware controllers.
- Software complexity: While Python lowers barriers, meaningful customization still requires coding skills, which may discourage users without programming backgrounds.
- Scalability: The open-source and DIY nature makes it excellent for prototyping and education, but less suited for high-throughput or research grade smart microscopy applications without further redesign and ruggedisation.

## PyCLM: programming-free, closed-loop microscopy for optogenetic stimulation

**Harrison Oatman [1], Jared Toettcher [2,3]**

[1] Lewis Sigler Institute, Princeton University

[2] Department of Molecular Biology, Princeton University

[3] Omenn-Darling Bioengineering Institute, Princeton University

### Specific Focus and scientific questions asked

How is cell signaling coordinated across entire macroscopic tissues comprising thousands of cells or more? This question cannot be answered without considering biology occurring at vastly different scales, from the minutes-timescale signaling states of individual cells to collective properties such as local cell density, tissue jamming, and diffusion-mediated ligand gradients across entire tissue regions. Receptor tyrosine kinases (RTKs) are at the center of many tissue-scale signaling processes in developing embryos and regenerating tissues. For example, waves of extracellular signal-related kinase (ERK) activity propagate for millimeters to centimeters across vertebrate tissues in a wide range of contexts, where they are thought to orchestrate collective properties of the tissue such as its density and directional migration. The study of tissue-scale signaling is enabled by two optical technologies: **live-cell biosensors** of cell signaling to report on the current activity state of all cells, and **optogenetic tools** to perturb signaling dynamics to better understand their specific functions. Many biosensors and optogenetic tools have been developed for studying receptor tyrosine kinase signaling at different nodes [6, 4, 1, 3, 7]. We may now ask a set of questions to (1) better understand the function of endogenous RTK dynamics and (2) deliver appropriate light inputs to control RTK dynamics and alter functional outcomes at the tissue scale (e.g. migration, cell density, regenerative phenotypes). Some of these questions include the following:

- How do traveling waves of RTK activity affect tissue-scale movement?
- How do specific dynamic properties (e.g. wave speed, spatial frequency) contribute to the outcome?
- Can we combine real-time measurement of cellular position with dynamic control of RTK stimulation to guide tissue flows?

Answering any of these questions requires the ability to deliver dynamic light inputs that vary both over time and in space to large numbers of cells. For the final question, one further capability is required: the ability to automatically update light inputs to hundreds or thousands of cells based on real-time measurements of cells’ current positions, a procedure that might be termed ‘closed-loop microscopy’ and which would constitute a subset of smart microscopy applications. Our PyCLM software package addresses all of the aforementioned challenges by providing a framework for imaging cells, segmenting them in real-time, and delivering optogenetic stimuli [5]. We were motivated by an important design criterion: it should be as easy as possible to set up experiments that combine any set of previously-defined imaging, segmentation, and stimulation procedures. Ideally, the end user (a new student in the lab, or a collaborator) should be able to combine any set of previous workflows without writing any new code at all.

#### Key findings and innovations

We used PyCLM to stimulate tissues of MCF10A breast epithelial cells with traveling waves of RTK activity. Indeed, we found that waves are sufficient to induce tissue translation: collective movement of cells in the direction of the wave. Wave speed played an important role in the migration response, with slow waves (∼1-10 μm/hr) driving slow migration, intermediate wave speeds (∼10-25 μm/hr) achieving optimal migration, and fast waves (>25 μm/hr) failing to induce tissue translation at all, instead producing small oscillations in cell position. We also explored other use cases of closed-loop microscopy, including using real-time cell segmentation to identify single cells in an MCF10A epithelial tissue, and then stimulating each cell to produce counterclockwise rotational tissue flows. Finally, we implemented an experimental workflow for single-cell feedback control of fluorescent intensity. In this workflow, cells are automatically segmented and their fluorescent intensity measured, and a bang-bang controller is used to apply light inputs to drive each cell to a desired fluorescent set point.

#### Methodology and implementation details

PyCLM integrates excellent tools that are already available for subsets of the closed-loop microscopy workflow. Microscope hardware control is provided by pymmcore-plus v0.13.7 and Micromanager [2] v2.03, whereas automated cell segmentation is implemented using Cellpose13 v4.0.6. All code and software requirements are available on the project’s Github page. Pymmcore-plus is available on Github as well. PyCLM is organized as a set of modules that pass information in a loop (Fig. 1A). A master scheduler ensures that each position is imaged and stimulated at the desired frequency, a microscope manager controls hardware through Micromanager, a segmentation module performs image segmentation using Cellpose, and a pattern module defines the next optogenetic stimulation pattern, which may vary depending on the outcomes of imaging and segmentation. These modules run as separate threads to maximize the number of experiments which can be run simultaneously. PyCLM can run multiple experiments simultaneously, with a simple schema for specifying the details of these experiments. The user chooses positions, sets an imaging schedule, and determines which experiments will be run by adding configuration files to a directory (Fig. 1B). The first step is to define a set of multipoint XYZ stage positions at which to carry out the imaging and stimulation experiments. These can be chosen interactively in MicroManager or proprietary imaging programs (e.g. NIS Elements) and conveniently exported. Assign a name to each position to identify the experiment associated with it. For example, a set of positions named “feedback.1”, “circle125.1”, and “circle125.2” will perform a feedback control experiment on one position (Fig. 1C) and stimulate with a 125 μm circular light pattern on the other two positions. Next, experiment configuration files in a Toms Obvious Minimal Language (.TOML) format specify the imaging conditions and stimulation parameters for an experiment. These files can be reused and modified or built from scratch using human-readable language. Finally, a master schedule is supplied in scheduler.toml to determine the timing of the experiments. Configuration files for currently implemented experiments (e.g. feedback.toml; circle125.toml) are provided alongside the software. Once directory setup is complete, PyCLM performs the experiment and continuously saves the results to the same location.

#### Contributions to Interoperability

PyCLM was written with interoperability in front of mind. The package is written to be agnostic of underlying microscope hardware provided that a stage (for multipoint imaging), camera (for image acquisition), and digital micromirror device (for optogenetic stimulation) are present. PyCLM is compatible with any MicroManager-defined channels and configuration groups. The output of PyCLM experiments is packaged as Hierarchical Data Format 5 (HDF5) files, an open-source file format that can be processed using a variety of downstream software tools, and allows for detailed export of experiment metadata.

#### Limitations

Our lab is actively using PyCLM for an increasing number of experiments, and new workflows are under construction. Nevertheless, it is still the case that a relatively small set of experimental workflows have been implemented to date, mostly focused on constant and wave-based illumination and intensity-based feedback control. Also, while our software should be interoperable with other Micromanager-controlled microscopes, it has so far been used only on a single hardware system. Finally, while we have implemented cell segmentation, the current version of PyCLM does not support single-cell tracking over time. This restricts PyCLM to using feedback controllers that only require current, instantaneous information about cell states (e.g. current intensity). Adding support for single-cell tracking and memory of prior states is a top priority for future versions.

**Figure 1.**
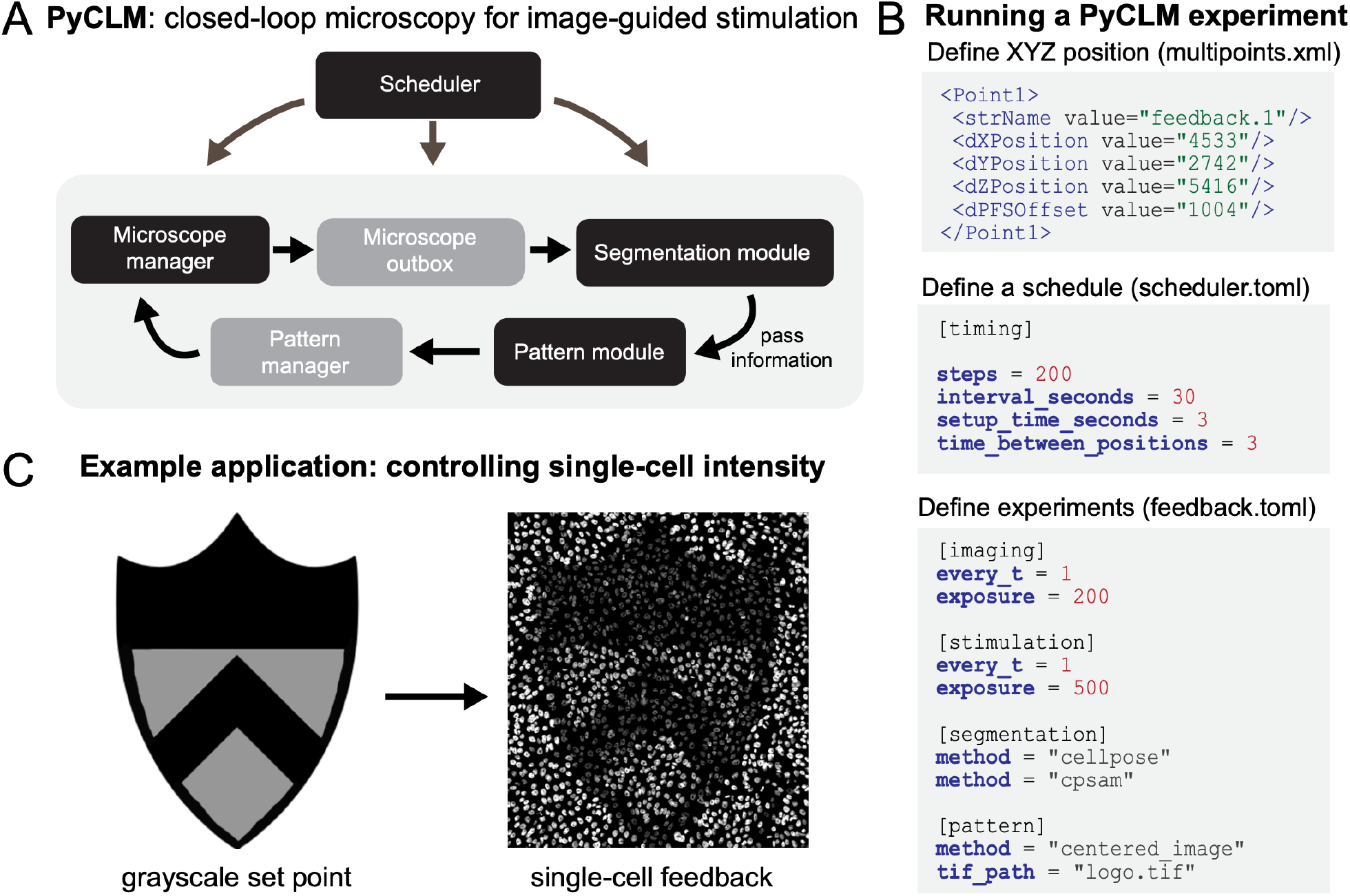
Closed-loop microscopy using PyCLM. **(A)** Schematic of the code organization. PyCLM is written as a series of modules that pass information in a loop. A microscope manager controls the underlying imaging and stimulation hardware, a microscope outbox collects the resulting imaging data, a segmentation module performs image segmentation, a pattern module uses the segmentation and imaging results to define a new optogenetic stimulation pattern, and a pattern manager stores the patterns and sends them at the appropriate time to the microscope manager. **(B)** Designing a PyCLM experiment is performed using a series of configuration files. A multipoints.xml file stores information about the experiment to run at each stage position, and a scheduler.toml file specifies the overall parameters of the imaging/stimulation sequence. Each individual experimental workflow is specified in a corresponding .toml file, which defines the imaging configurations, segmentation methods, and stimulation pattern module to be run. **(C)** PyCLM enables many experimental workflows including constant and dynamic light patterns, real-time optogenetic stimulation of migrating cells, and feedback control of fluorescence intensity, shown here for MCF10A cells expressing the TagRFP photochromic protein.

## Inscoper

**Claire Demeautis, Célia Martin, Rémy Torro, Thomas Guilbert, Otmane Bouchareb** INSCOPER SAS, Cesson-Sévigné, France

### Approaches to smart microscopy

Smart microscopy stands at the forefront of our R&D initiatives, with innovative applications continually emerging. In this context, we introduce two of our advanced solutions, each designed to address specific application requirements: the **Roboscope**, optimized for the detection of rare cellular events, leveraging sequence interruption, and the **Python script call module**, seamlessly incorporated within our image acquisition software.

#### 1. Roboscope: methodology, implementation details

Our approach focuses on developing an innovative software solution to make microscopes smarter and more autonomous. The software combines light microscopy experiment acquisitions, advanced analysis algorithms, sequence interruption and deep learning integration to enable custom event-driven acquisition. The system is adaptable to various experimental workflows, enabling the capture of transient events while minimizing data consumption.

We designed an automation technique capable of interrupting a main sequence to start a new one, based on the outcome of an image analysis workflow over the image stream. The new sequence may also be interrupted, repeating this pattern as needed. The Roboscope is able to screen a sample at low magnification looking for an *Event Of Interest* (EOI, *i*.*e*. mitosis here). Upon detection of such an EOI by the image analysis workflow, the main screening sequence is interrupted and a new sequence at higher magnification takes over, using parameters returned by the image analysis workflow. This solution requires an Inscoper device controller, the Inscoper Imaging Software and a CPU. A graphical processing unit (GPU) is recommended to increase the computation speed in deep learning applications but not required. While the device controller runs the sequence, the images are analysed through the CPU and GPU. As soon as an EOI is detected, the running sequence is interrupted, new acquisition parameters are calculated and sent to the device controller.

Users can design workflows for their specific applications, defining the targeted event and the consequential actions. The workflow is set up graphically via an ergonomic user interface (Figure 1). Theoretically, new model training takes less than 30 minutes on a Nvidia GeForce RTX 2080Ti GPU.

**Fig. 1:**
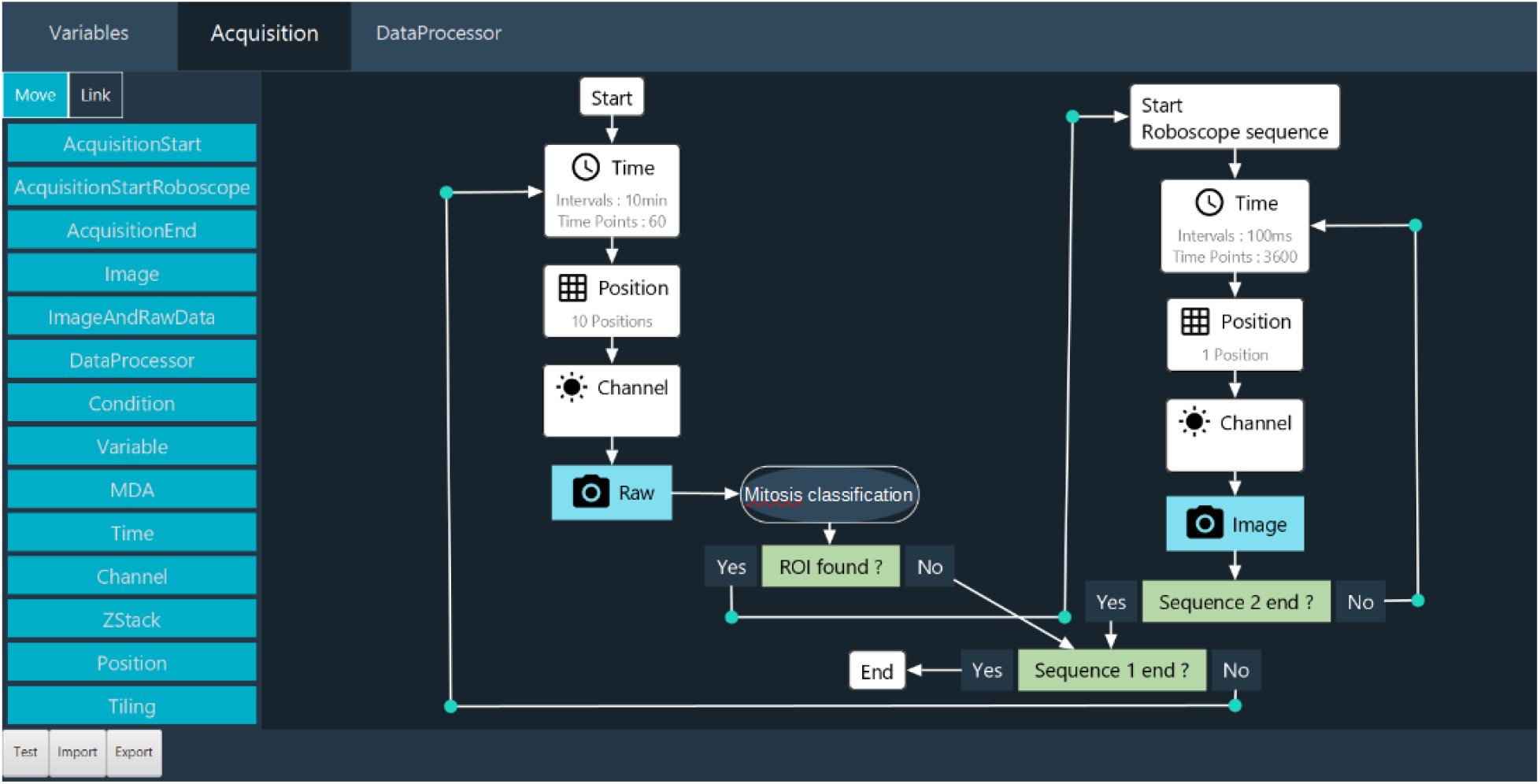
Roboscope GUI.

The key innovation of our development is the interruption sequence during a sequence. With this method, the user doesn’t miss an interesting event. The main features of our approach are :

- **Adaptable**, with a modular software architecture that allows the EOI detection to be modified according to the user’s needs. A novel network embedding, sGAN, can be fine-tuned to a wide range of biological applications. Users can also trigger their own Python scripts, custom-made for their applications of interest.
- **Easy to use**, without coding skills on a graphical interface, to retrain the models.
- **Fast**, with optimised microscope control and sequence interruption, dedicated modules running in parallel.
- **Parsimonious**, by limiting photobleaching and phototoxicity and only sending qualified events to the computer. This considerably reduces the amount of data generated and saved.

#### 2. Python script call module: methodology, implementation details

Within the Inscoper Imaging Software (IIS), we have developed a module that allows users to integrate custom Python scripts to perform smart microscopy tasks. This module, incorporated into our Multidimensional Acquisition (MDA) sequence builder, is triggered upon image reception. It enables adaptive-feedback microscopy: real-time adjustments to device parameters based on outcomes from image analysis. Additionally, it enables users to write metadata about the image content and dynamically display informative visualizations during acquisitions. Code-wise, users define two functions:

**Fig. 2:**
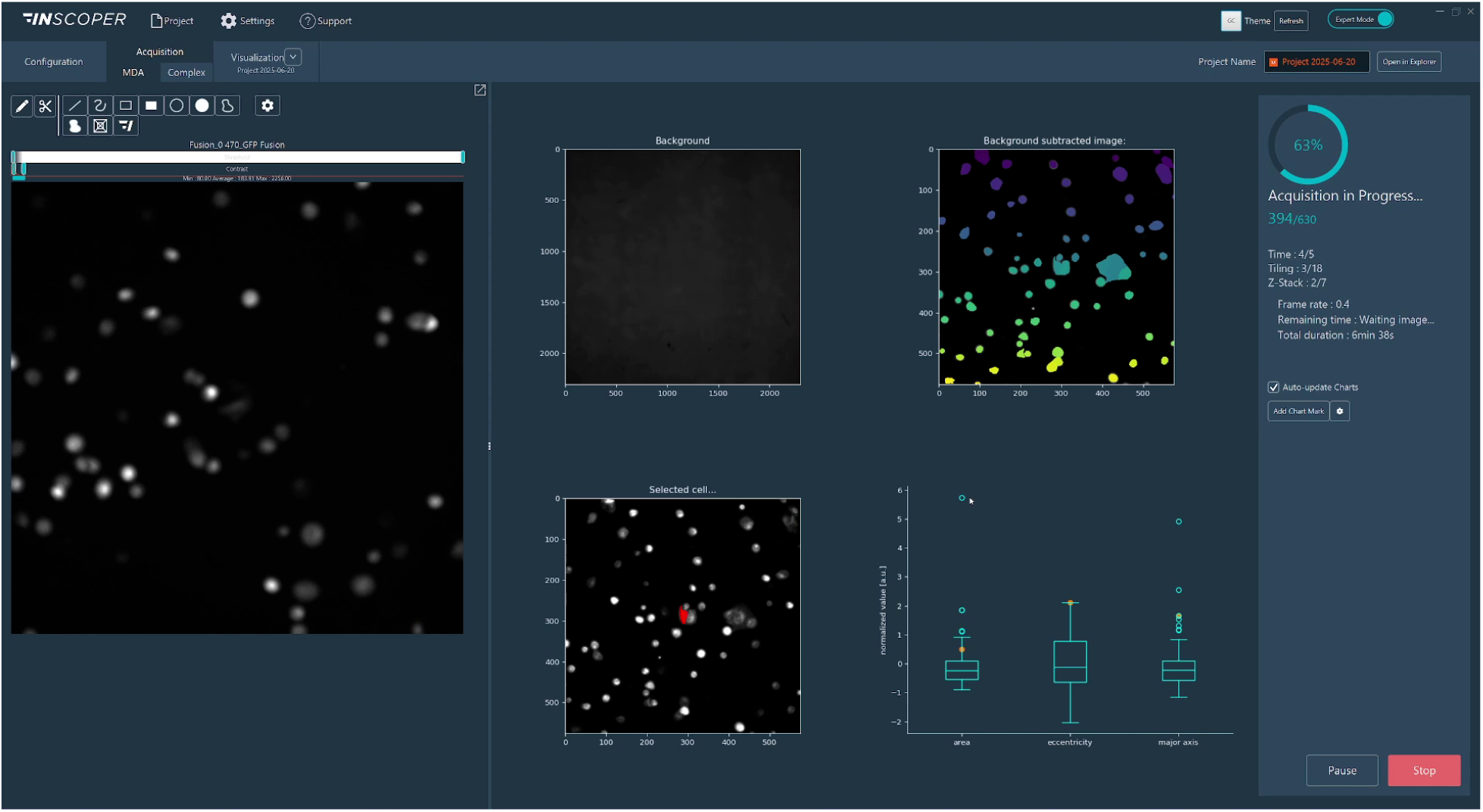
Real-time visualizations from a Python script in IIS.

As part of the IIS framework, we bundle a Python distribution pre-installed with commonly used libraries for image analysis. We enable users to switch to their system python installation or one of their custom environments so that they can use the libraries they are familiar with and benefit from any hardware-specific installation, such as GPU-optimized libraries. To streamline the scripting process, we provide a utility library that simplifies conversions between image space and hardware space (for instance, converting pixel displacement into lateral X and Y stage displacements), writing custom image metadata, and transmitting figures (generated using `matplotlib, seaborn`, or `plotly`) directly into the IIS interface.

- Image-provider function: Uses metadata from the most recently received image to check the current acquisition context and select images requiring processing. Typically, this selection may be empty in scenarios such as mid-stack Z-plane captures or initial channel acquisitions within a multi-channel sequence, effectively preventing unnecessary computations.
- Processing function: Accepts the image selection identified by the image-provider function and performs analytical operations. The outputs from this function can include:
  – Updated device parameters for adaptive feedback
  – Quantitative image metadata entries
  – Saved files (e.g., cropped images, segmentation masks, image reconstructions)
  – Real-time visualizations displayed within IIS (dashboard-like)
  – Any combination of the above

Graphically, users import their custom Python script (.py file) into the Python script module and associate the defined image-provider and processing functions. Initially, all function arguments and outputs are explicitly matched by type and value. Users can subsequently save presets, retaining only adjustable arguments that are not tied to specific hardware parameters.

#### Contributions that could contribute to interoperability

Our acquisition solution includes a device controller and software that is compatible with almost all camera-based microscope setups and all their associated devices on the market (supported device list). Also, using the agnostic useq-schema to describe the main sequence and its various dependencies is easy to implement in our Inscoper’s Roboscope. Finally, Inscoper is thinking about its acquisition workflows and machine learning training in terms of open source availability, with the aim of forming a community ready to exchange, share and populate a database enabling each user to lift their technological barriers.

#### Current bottlenecks, Roadmap

Real-time device updates currently affect acquisition performance, creating limitations for certain optimizations with our Inscoper hardware. In general, fully unrestricted adaptive microscopy—where every device parameter can be adjusted at any time—may lead to suboptimal system efficiency. To mitigate this, pre-defining which devices and acquisition structures are modifiable helps maintain performance and stability. The Roboscope addresses this challenge by pre-configuring successive acquisition sequences tailored to specific applications.

We aim to enhance virtual device simulations to better replicate real-world acquisition and smart microscopy scenarios. For example, a virtual camera could simulate image acquisition by cropping from a larger mosaic image, where adjustments to X and Y parameters effectively shift the crop region, emulating believable microscopy interactions.

We want to apply our approaches to more biological applications and systems. We will soon integrate a new microscopy modality (photomanipulation) into our smart microscopy products.

## Open Application Development (OAD) in Zeiss ZEN

**Philipp Seidel**, Carl Zeiss Microscopy GmbH

### Approaches to smart microscopy

The user base of ZEN is diverse both in terms of research / application focus and technical expertise. Hence, we aim at offering Smart Imaging interfaces at different levels of integration with ZEN’s main user interface. In the following sections I aim at illustrating these different tools and, their key features, their key applications, limitations and if available, references to further material.

#### Guided Acquisition / Automated Photomanipulation

**Fig. 1:**
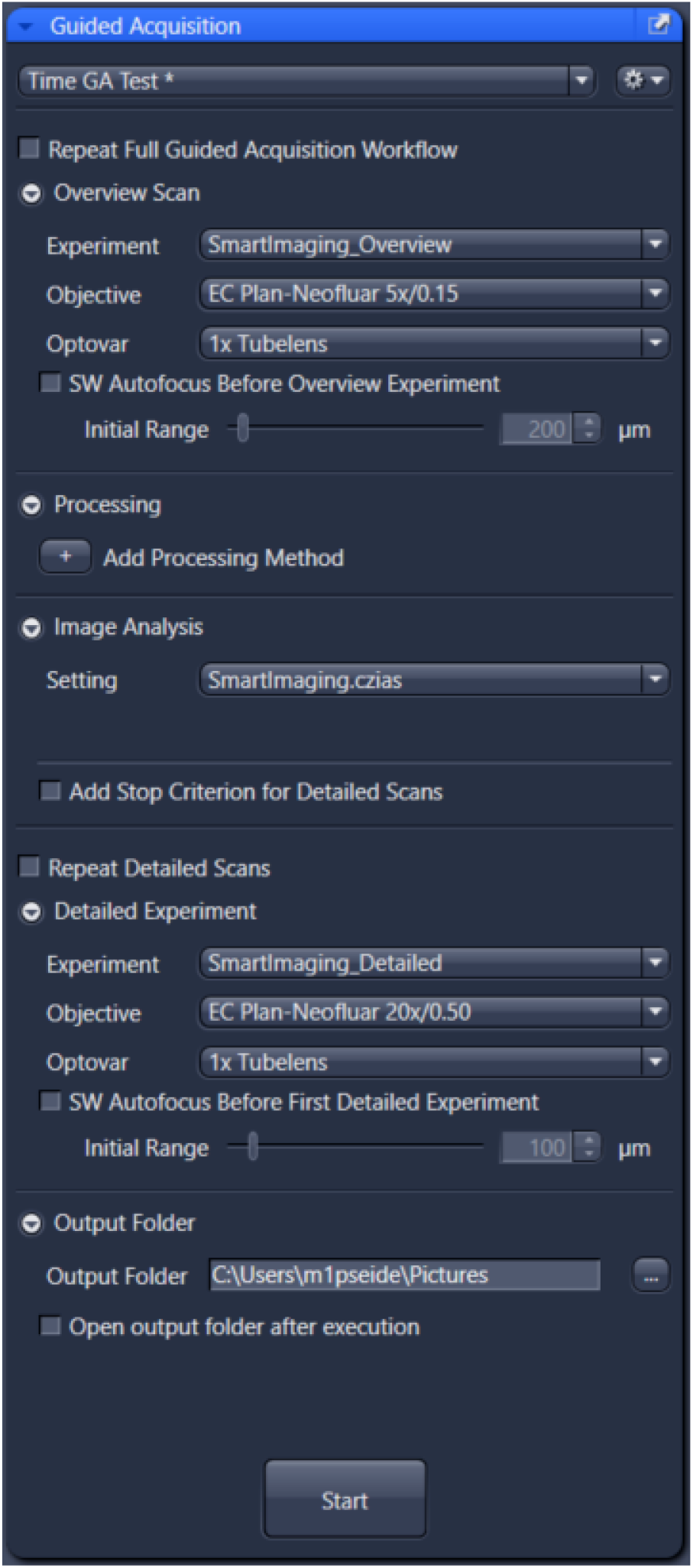
Guided Acquisition / Automated Photomanipulation.

These GUI-based tools offer a simplified entry into typical Smart Imaging workflows addressing imaging novices and scientists without access to coding expertise.

The **Guided Acquisition (GA)** user interface is shown in Fig. 1. It follows a fixed workflow, where an overview scan is followed by an image analysis block and several detailed scans. Positions (FOVs) of the detailed scans are derived from the image analysis. To keep the user interface simple, the user plugs in pre-configure experimental and image analysis settings for the three main steps. The user can furthermore add certain optional components, e.g. additional processing steps for the raw overview image, or manually adjust the number of detailed scans. Time-lapse acquisitions can be defined to allow for detection of sparse events.

GA builds modular on top of generic ZEN functionality like Acquisition Tab and Image Analysis Wizard. Besides simplifying usability, this comes with the benefit that other built-in features (like Deep Learning segmentations) also seamlessly work with GA. GA also is compatible with image data correlation via ZEN Connect, or high throughput well plate experiments.

Typical applications are high-resolution 3D imaging of sparse samples like embryos or organoids. Here, via Smart Imaging, the main workflow improvements are reduced data sizes and time-to-result. Follow the links at the end of this section for more details on such use cases.

Automated Photomanipulation follows a very similar approach. In a dedicated UI, the user assigns a pre-configured acquisition experiment and image analysis scheme. The notable difference is that instead of defining FOVs for subsequent high-resolution scans, the workflow defines image sub-regions that are subsequently bleached and tracked (by time-lapse acquisitions) to determine fluorescence recovery (FRAP) or similar read-outs. Besides, this tool could also serve simple opto-genetics experiments.

Links:

- Guided Acquisition OAD documentation
- Guided Acquisition Webinar
- Smart Workflows with Celldiscoverer 7

#### Experiment Feedback

**Fig. 2:**
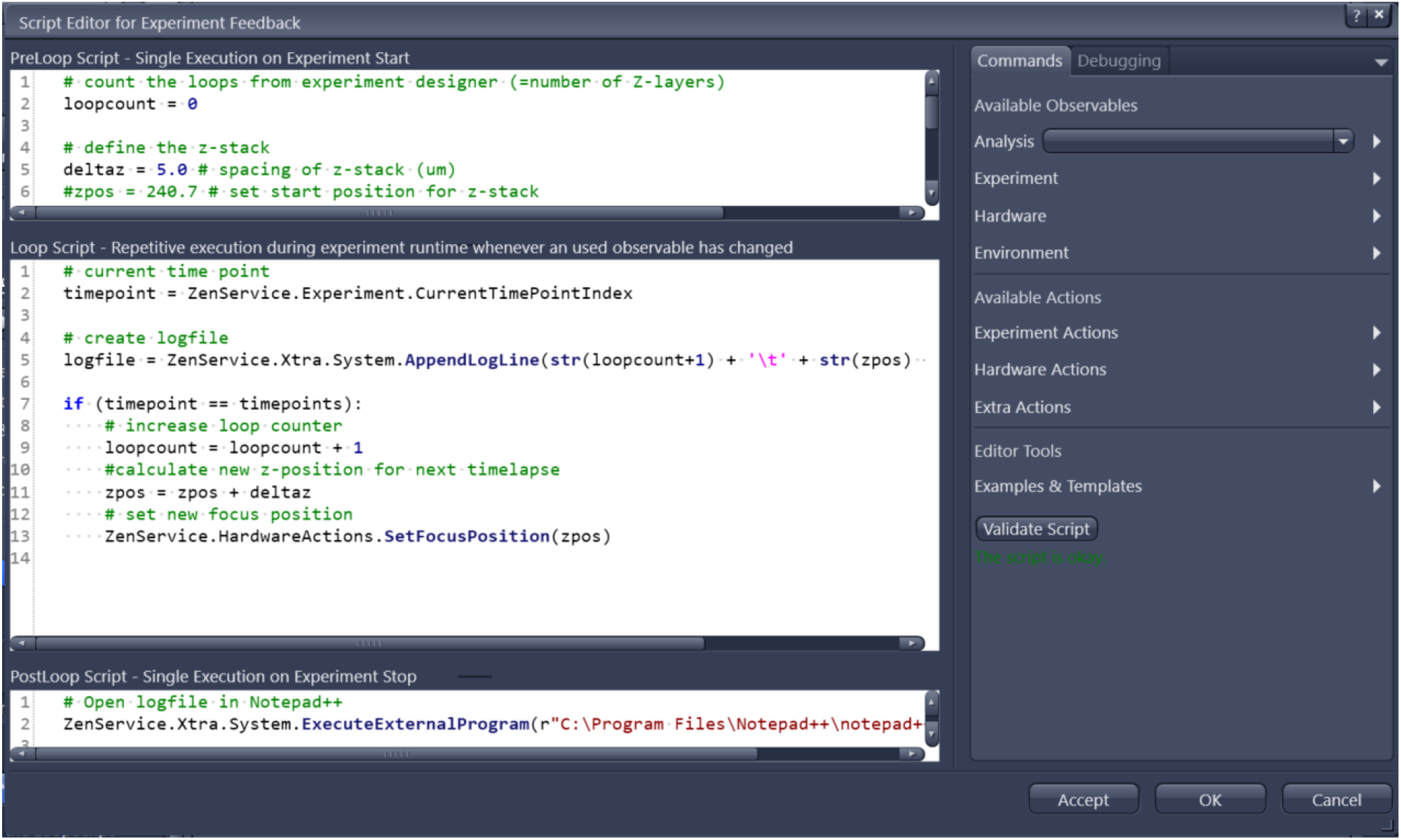
Experiment Feedback.

This is basically an advanced scripting tool directly integrated into a ZEN acquisition setup. It does require coding but is simpler to set up and in certain situations more powerful than OAD scripts.

The crucial feature of Experiment Feedback is the possibility to tune many experimental parameters “on-the-fly” during an ongoing acquisition. This includes parameters like exposure time, lamp intensity, time series intervals, objectives, stage positions and many more. It features (like GA) a pre-configured image analysis to measure sample parameters and automatically respond to them.

Some use cases become very easy to implement with Experiment Feedback, e.g. tracking objects during acquisition. Please find more examples in the linked Github page.

Links:

- https://github.com/zeiss-microscopy/OAD/tree/master/Experiment_Feedback
- ZEN Developer Toolkit

#### Macro Editor / Simple API / Object Model

**Fig. 3:**
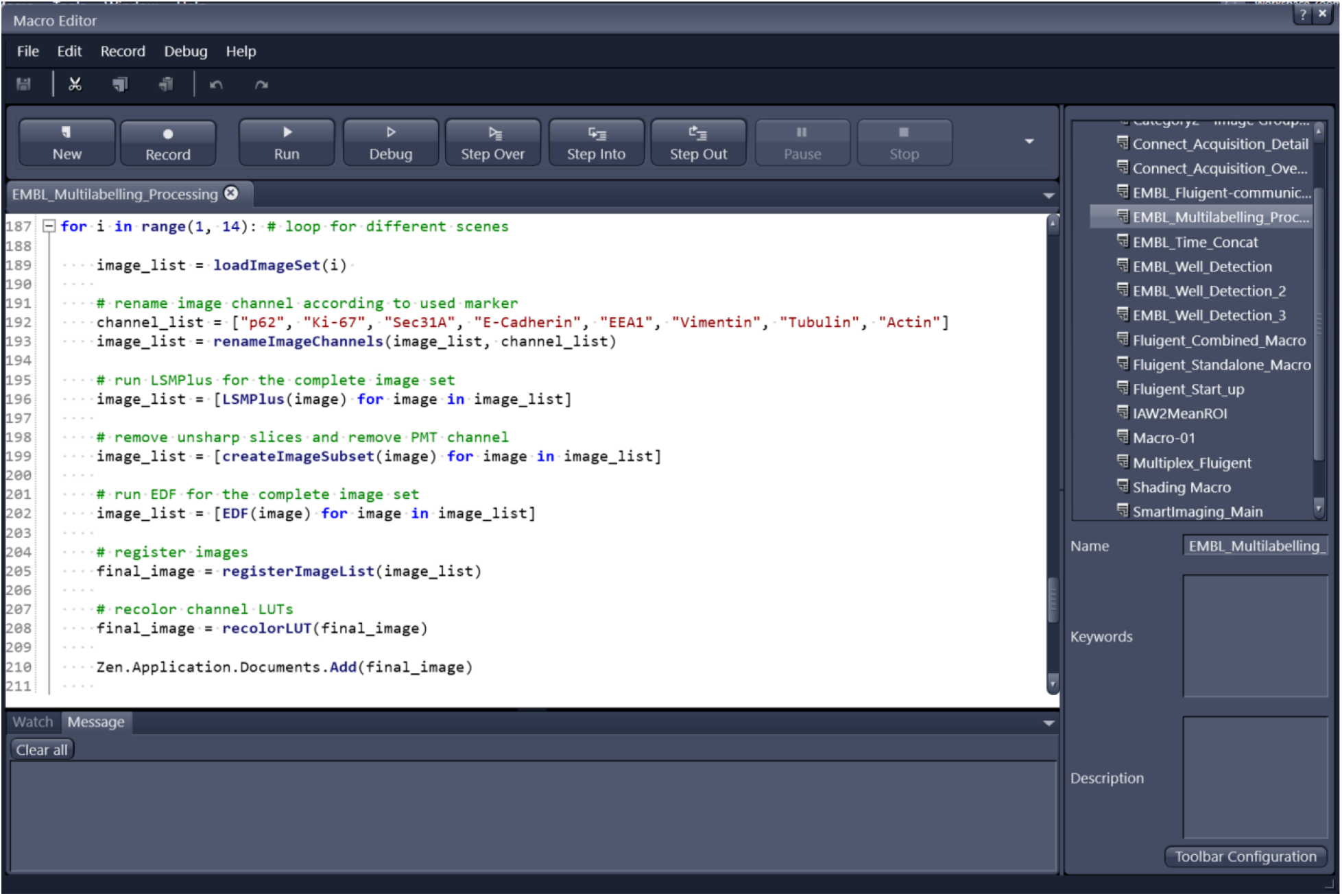
Macro Editor.

The main feature of OAD, as of today, is the scripting interface called “Simple API”. Most importantly, it comes with a small macro editor integrated in ZEN, and it offers interfaces for external applications written e.g. in Python (see links below for more details).

Internally, the Simple API uses IronPython, which is a C# implementation of Python with the purpose to serve .NET applications. Most noteworthy, this allows for implicit use of ZEN’s object model, hence, direct access to experimental setups, processing functions or czi image, and their respective methods. The macro editor is equipped with debugging, code-completion, recording (of a user’s interaction with ZEN) and documentation, simplifying onboarding and troubleshooting.

For external applications, COM and TCP-IP interfaces can be used. This allows tackling more complex use cases that requires a more refined code structure or additional libraries.

Simple API is most useful for integrating standard ZEN functionality into custom workflows for the purpose of process automation. Standard smart microscopy is covered by accessibility of basic HW functions like stage movements and objectives. However, it is not designed to change deeper acquisition parameters dynamically (like Experiment Feedback), and therefore has limitations for advanced smart imaging approaches.

Links:

- Various scripts
- Interfaces
- Application
- Developer Toolkit

## ImSwitch for openUC2

**Benedict Diederich**, openUC2 GmbH

### Specific Focus and Scientific Questions Asked

openUC2 integrates modular microscopy with advanced automation to explore dynamic cellular behaviors and visualize individual fluorescent objects or scan large microscopic objects, ideally making these capabilities universally accessible. Our primary goal is to support researchers who lack access to cutting-edge microscopy technology, enabling them to conduct sophisticated experiments across diverse fields, including virology and all areas of education. Leveraging Arkitekt’s integration, we aim to standardize microscopy control workflows, simplifying the setup and execution of complex imaging experiments that may also include auxiliary hardware such as robot arms or pipetting robots.

We are developing a versatile platform that encompasses a broad array of tools suitable for smart experimental designs, particularly those that require more than a single image capture and benefit from combining various modules to conduct intricate experimental protocols. Our main focus lies on live-cell imaging, such as characterizing bacterial responses to bacteriophage exposure under different environmental conditions. Through this approach, we aim to systematically analyze effects related to temporal dynamics, spatial positioning, and morphological changes. Ultimately, our platform seeks to significantly lower the entry barrier for advanced microscopy experiments, democratizing access and fostering broader participation within the scientific community. This we do by providing parts that feature a lower price tag and increased simplicity by making different parts compatible with each other using the openUC2 cube-like toolbox. With this openUC2 aim to **become the Raspberry Pi for optics**.

**Fig. 1:**
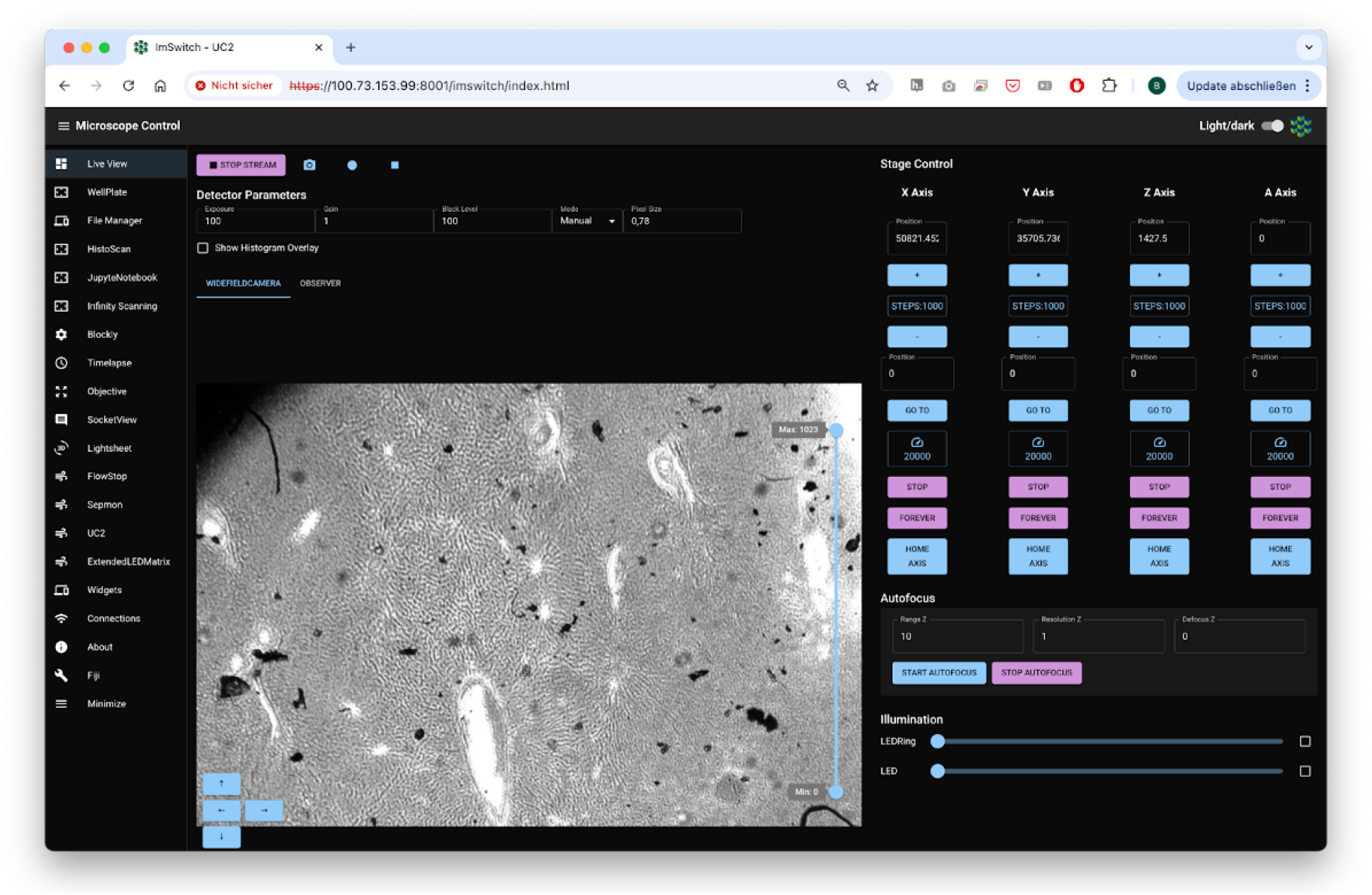
Standard GUI from openUC2’s ImSwitch in the browser.

#### Key Findings and Innovations

openUC2 cubes controlled through ImSwitch combined with Arkitekt provide modular, scalable microscopy setups that dynamically respond to smart events in multiscale microscopy. We took great care to make these experiments as reproducible as possible and ensure a platform independent operation. For this, we created Docker containers for the different apps such as microsocopy control, image processing and hardware interaction, while drivers for hardware control are embedded. This ensures reproducibility and standardization of microscopy environments across diverse research setups, ensuring consistent and reliable outcomes - independent from the underlying Python environment. Integration with Opentrons robots and robotic arms further expands automation beyond imaging, facilitating comprehensive experimental workflows including sample preparation, reagent addition, and plate handling. The modular nature of openUC2 significantly simplifies setup complexity, enhancing throughput and reproducibility in smart microscopy experiments.

**Fig. 2:**
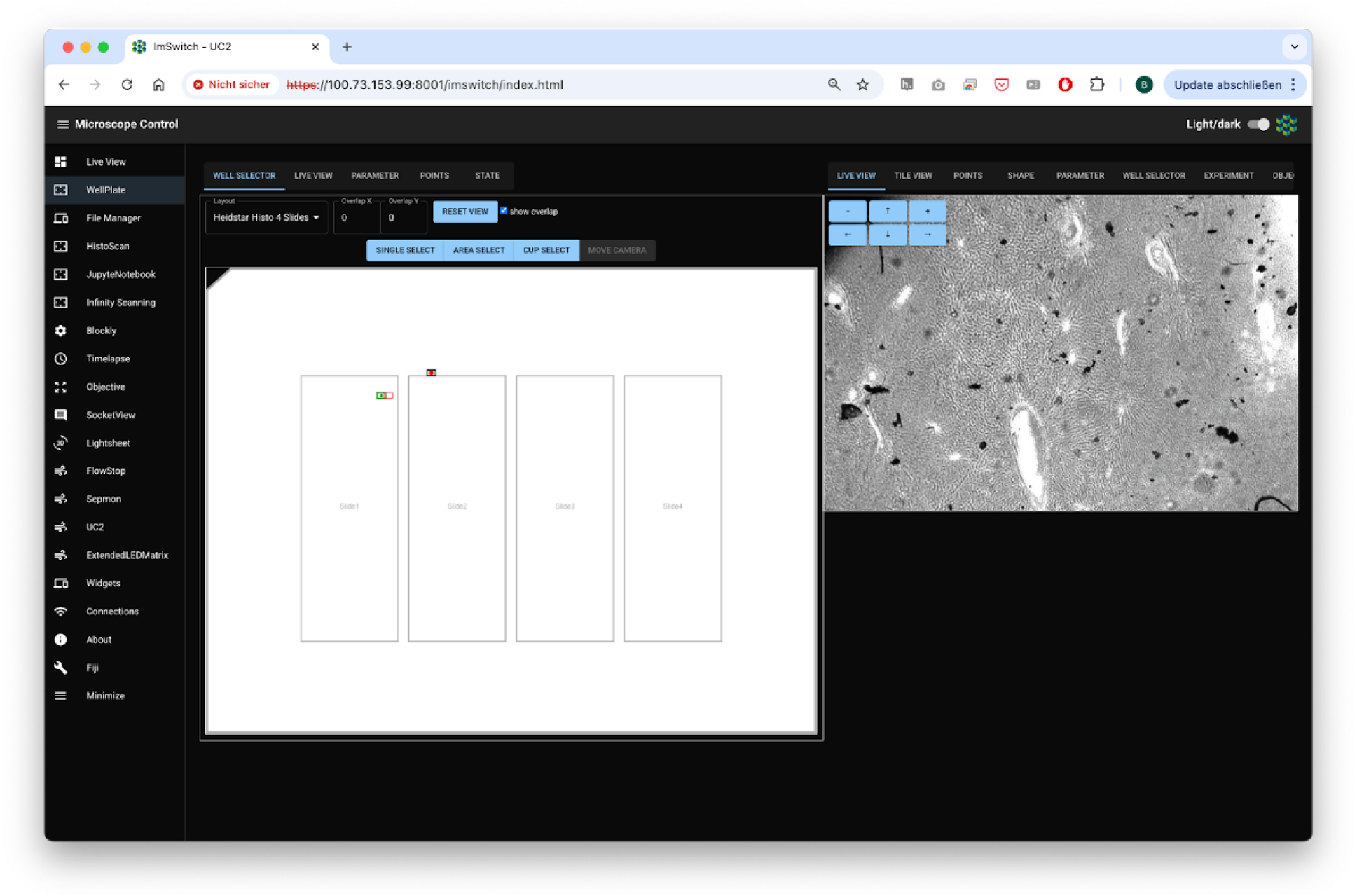
Different Apps in form of controllers enable more complex experiments including the visual graphical interface. The operational model splits into educational and professional aspects. Educational initiatives include introductory Jupyter Notebooks and workshops designed to teach students how to build smart microscopes, capturing their first images and developing fundamental algorithms such as autofocus. Knowledge gained in educational settings is directly transferable to professional equipment, ensuring consistency as the software and hardware platforms remain uniform across both educational and professional contexts. Jupyter notebooks have full access to the hardware and can include any external python library. It can run on the same hardware as the microscopy server (e.g. Raspberry Pi) that also drives the hardware or on a remote cluster HPC if more computational power is desired.

**Fig. 3:**
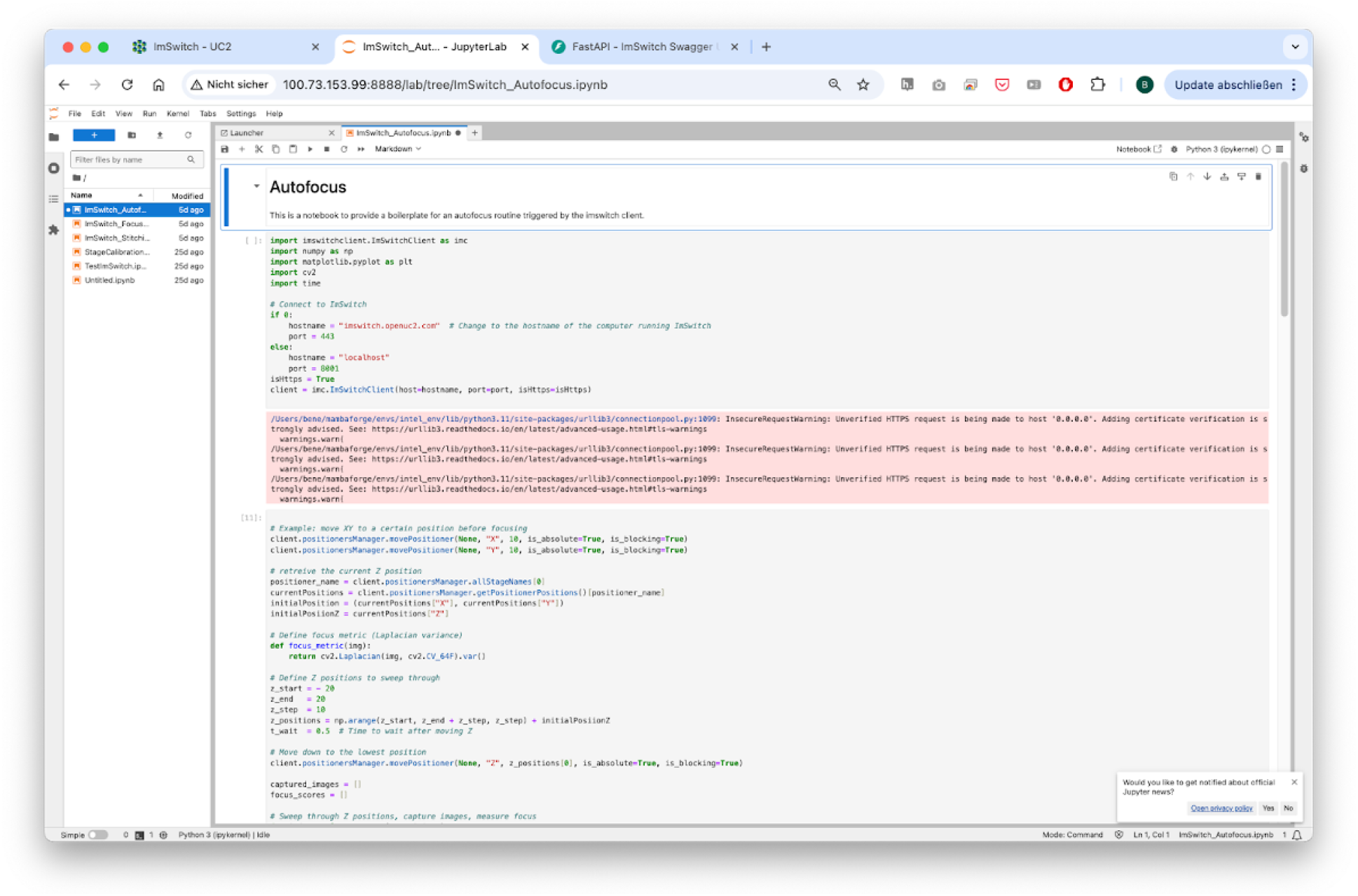
A Jupyter Notebook Server can offer the creation of simple smart microscopy workflows by using common image processing pipelines and direct in-browser visualisation.

#### Methodology and Implementation Details

openUC2 microscopes rely on the already well established ImSwitch microscopy control software that was implemented in Python. We have ensured that the Model-Viewer-Presenter scheme of the underlying architecture fully avoids the use of QT component to make it more lightweight and operatable through the web. This also makes the use inside docker container a lot easier since information can be transported through TCP/UDP packages. In collaboration with developers from the Forklift framework, we have developed a dedicated operating system based on Raspberry Pi OS, embedding all necessary drivers and software components directly onto a flashable SD card. The compilation process is fully automated through GitHub Actions, enhancing maintainability and portability.

A dedicated workflowmanager inside ImSwitch can formulated complex workflows as a list of tasks, while they are first formulated and compiled into hardware instructions before they are carried out on the hardware itself. The tasks can be provided by a python dictionary of hardware (e.g. capture image, move stage) or software events (e.g. triggers, waits, image processing results) or by a list that is uploaded via the REST API.

**Fig. 4:**
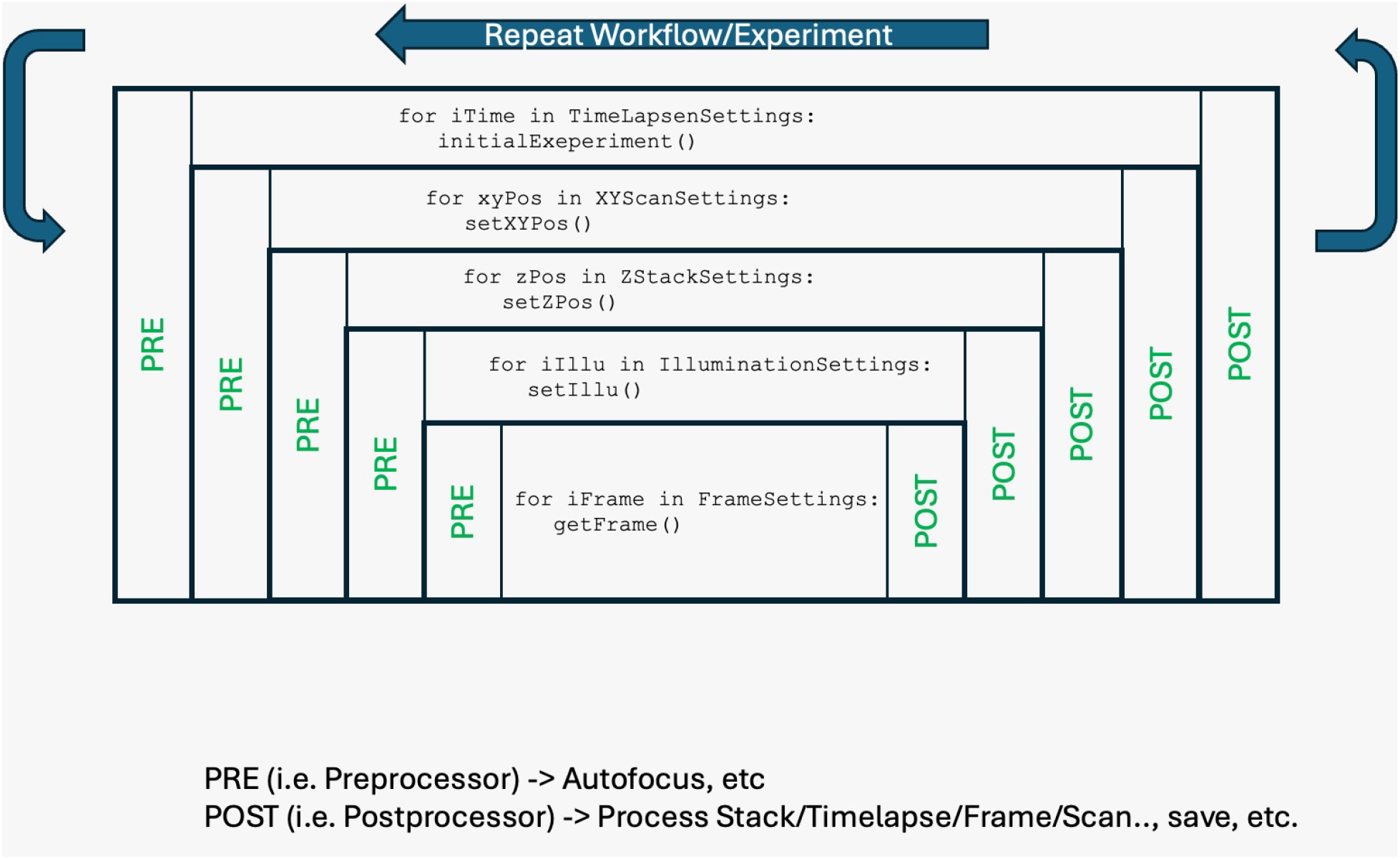
An example how a workflow in ImSwitch can be formulated. Every stp has a pre and post function that is exucted. The graph is build up first, before the instructions are executed in a timely manner. One example to formulate this list visually is blockly. It creates a JSON object with all the necessary steps that will be interpreted by the ImSwitch backend and executed on the hardware.

**Fig. 5:**
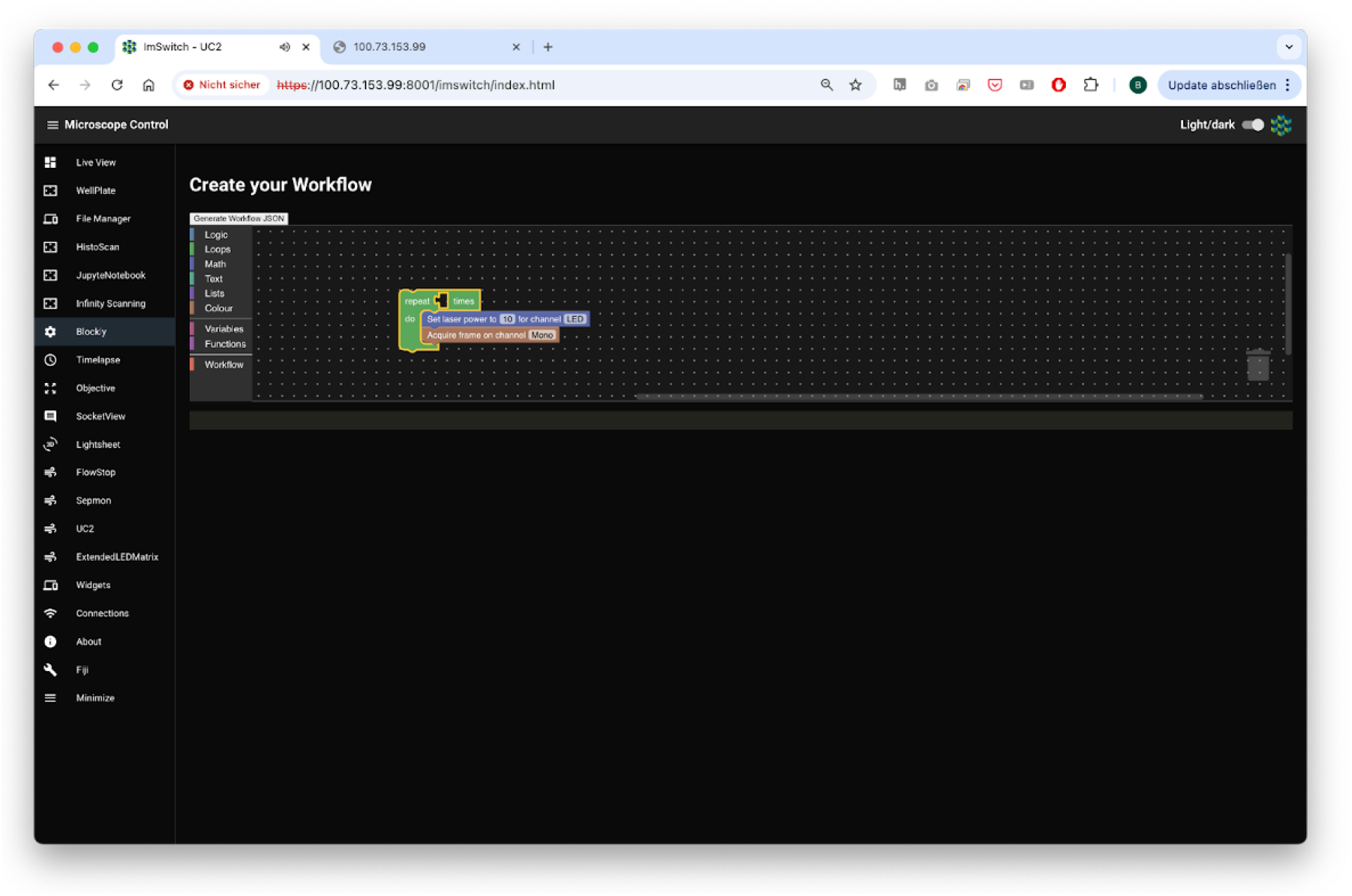
Blokly converts the different steps into a json object that is then later interpreted by the Workflowmanager. Our approach utilizes Docker containers for virtualizing software components, such as the imSwitch system, device drivers, and image processing tools. This significantly simplifies software management and ensures reproducibility across platforms. All device control commands are issued via REST APIs, signal updates are offered through a websocket connection. Instead of relying on the QT-based gui int he former ImSwitch variants, we have constructed a React-based web application that integrates seamlessly into the Arkitekt network, providing intuitive access to most available parameters and functions as defined services. Here, the microscope becomes a dedicated micro-service, that can for example offer the functionality of scanning a 2D area or vary the optical resolution by changing the objective lens. The dataflow inside Arkitekt enables image processing on dedicated hardware before a control command is back-channeled to the microscope to interact with the sample in a smart way. This not only includes microsocpes, but can be widened to include robots and alike.

**Fig. 6:**
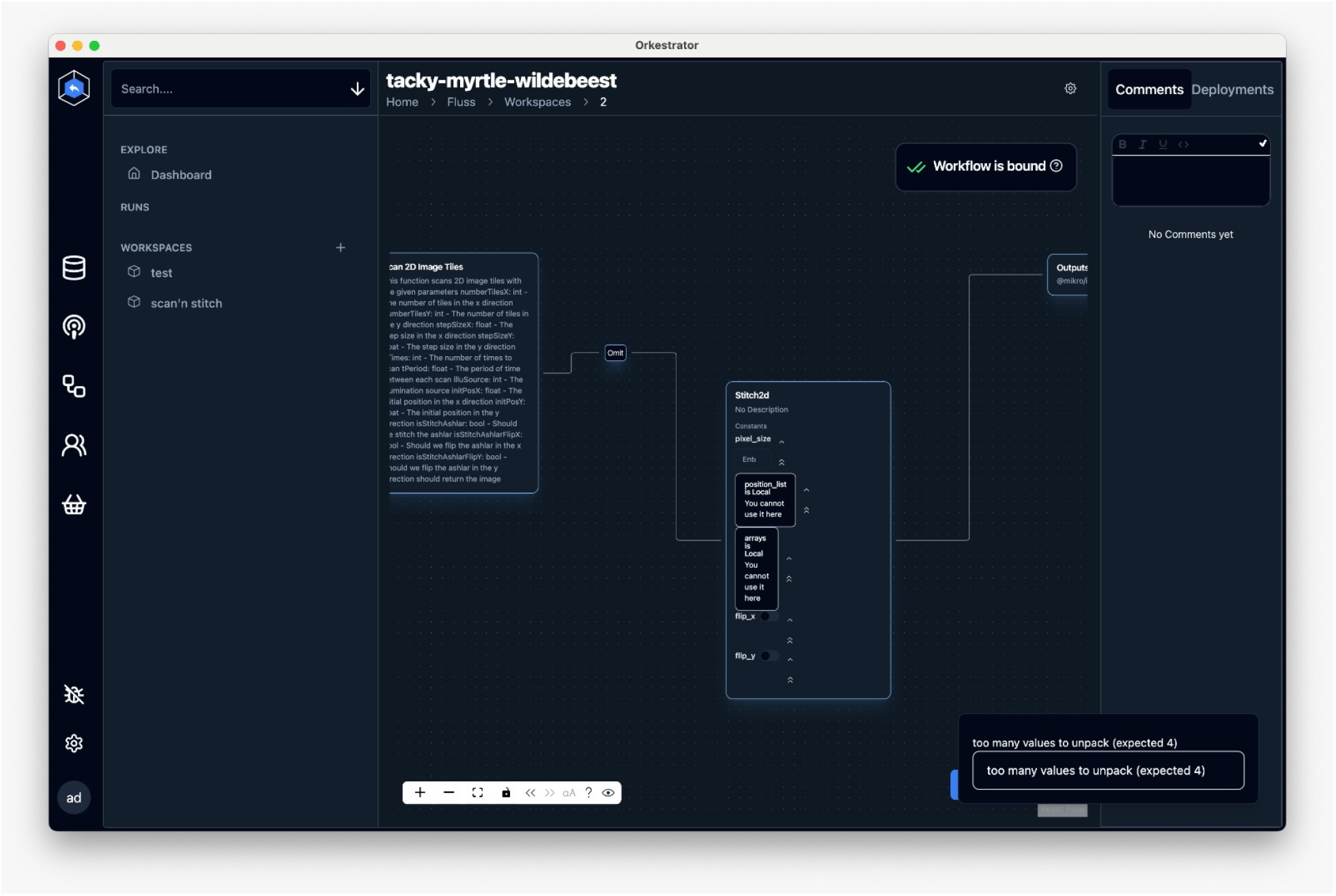
The Arkitekt framework enables a simple way of creating complex workflows where multiple hard- and software components can be joined in a highly decentralized manner. Data flow management relies on a microservice architecture, where data accompanied by metadata is streamed from individual nodes to a central Arkitekt server node and subsequently distributed through lazy loading to processing or storage nodes. This abstraction layer ensures compatibility with common formats such as OME-Zarr or OME-TIFF, facilitating efficient data handling and interoperability.

### Contributions to Interoperability

openUC2 is entirely open source, committed to creating a highly interoperable ecosystem where diverse components seamlessly interact through standardized interfaces. In addition to that, we have created a firmware specifically designed for the ESP32 microocontroller. This firmware is designed to control various hardware components such as lasers, LEDs, motors, and galvo scanners using a unified codebase adaptable across multiple hardware configurations. This flexibility is further enhanced through CAN and I2C interfaces, facilitating the integration and control of external components. Low-level hardware control emphasizes human-readable interoperability through JSON-based dictionaries, which simplifies adding new parameters and functionalities. To facilitate rapid testing and interaction, we developed a web application leveraging WebSerial API, enabling immediate hardware access without additional installation. This setup supports straightforward experiments, such as browser-based time-lapse imaging using standard USB and UVC webcams, further emphasizing ease-of-use and accessibility.

**Fig. 7:**
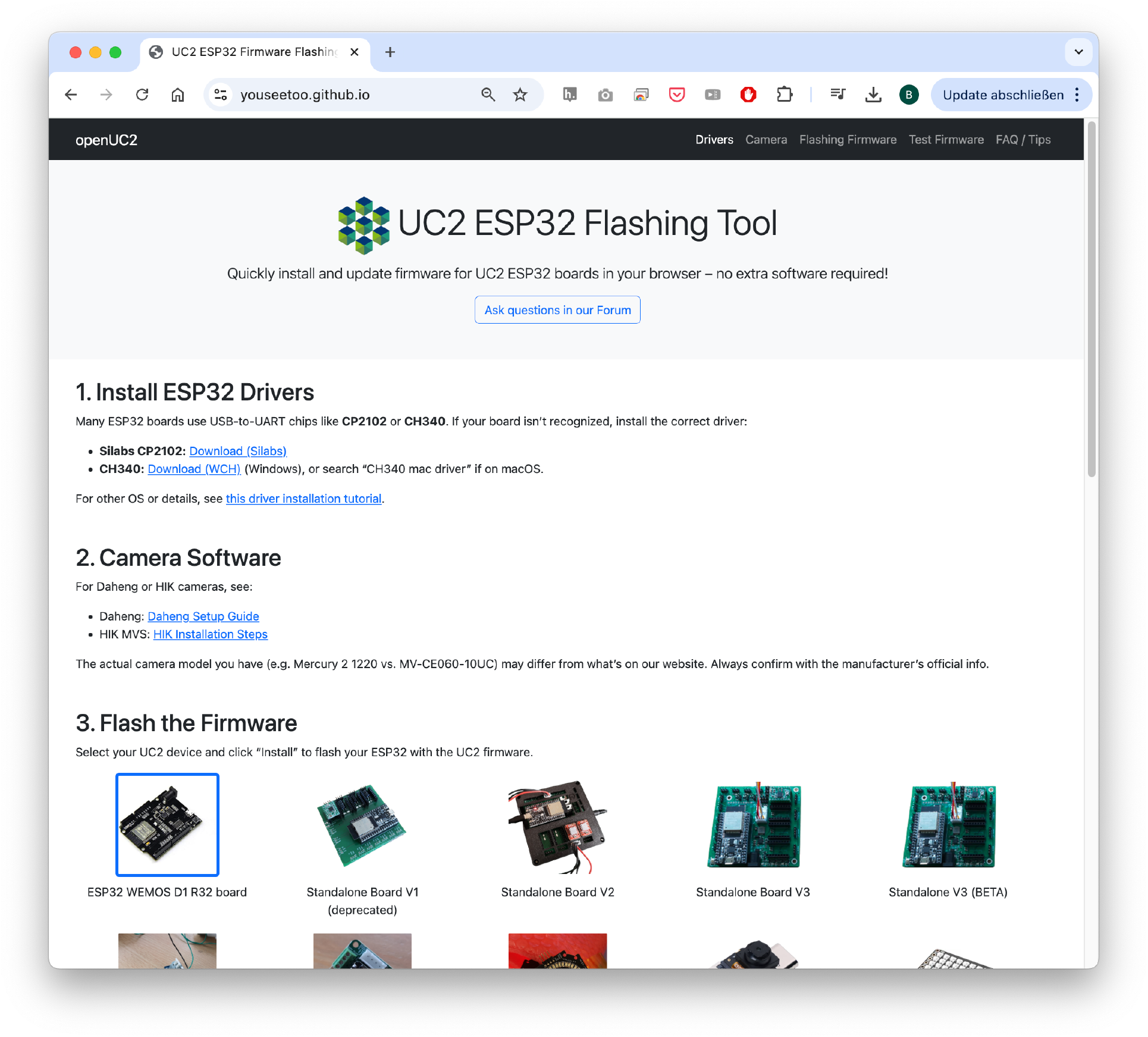
To further lower the entry barrier to get starting with smart microscopy experiments, openUC2 offers the abbility to flash the firmware to the ESP32s using the openUC2 Flashing tool in the web.

**Fig. 8:**
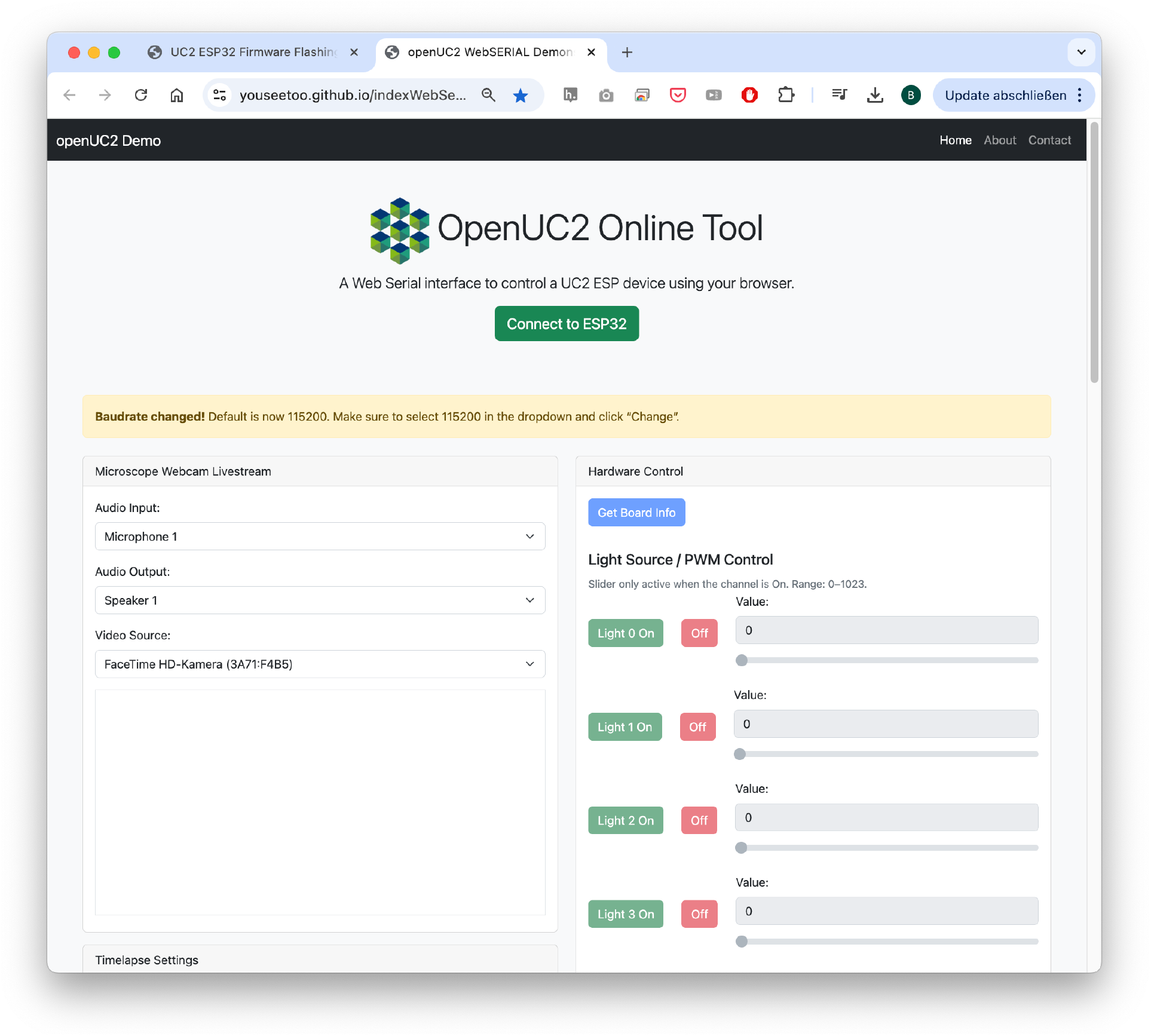
The Web-serial Standard enables the use of usb serial devices without the installation of additional software. A notable collaboration with FLIMLabs showcases this interoperability, where their open REST API was successfully integrated into the ImSwitch environment, enabling direct control and analysis of FLIM hardware. The combination of openUC2 and ImSwitch aims to foster an open ecosystem that allows extensive external interaction without restrictive agreements like NDAs.

**Fig. 9:**
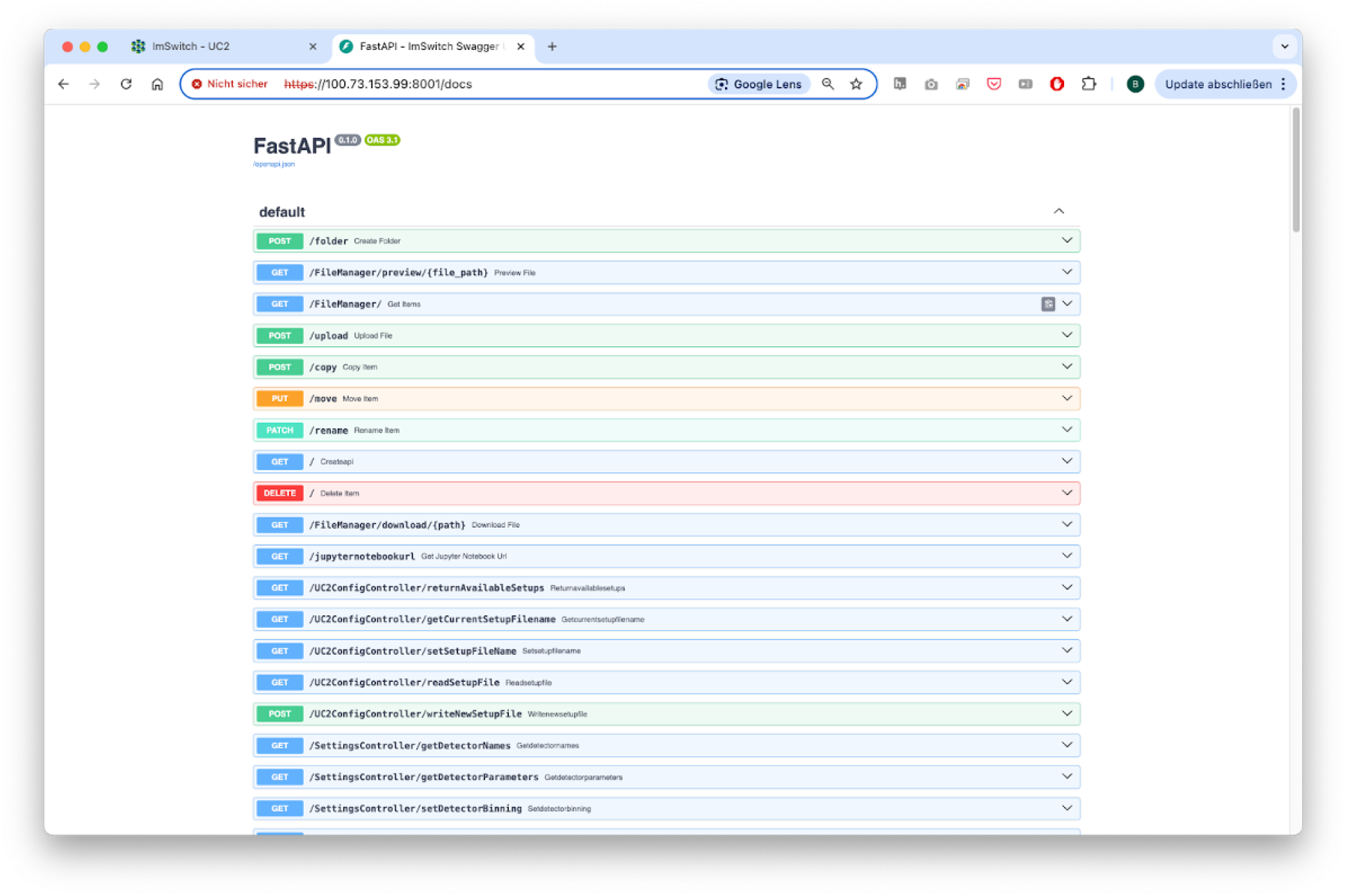
An open REST API documented through the openapi swagger schema enables a fast interaction with the ImSwitch backend through the browser.

#### Limitations

The development of openUC2 in the professional software domain is still relatively early-stage, making the system potentially prone to errors. However, by developing a dedicated operating system for platforms such as Raspberry Pi or Jetson Nano, we significantly reduce such vulnerabilities, as all drivers and software components are thoroughly tested and continuously integrated through GitHub Actions. In case of issues, software versions can easily be reverted to stable builds, minimizing downtime. Documentation remains incomplete, presenting challenges for new users integrating their interfaces with our software. We actively encourage community feedback and participation, providing resources through openUC2’s GitHub repository and forums to foster a collaborative environment for addressing questions and continuously improving our documentation.

Here’s a structured list of relevant links you should consider including to make the post complete and easily navigable for readers:

#### Core Software Repositories and Documentation

- **openUC2 GitHub organization** – all related hardware and software repos Github
- **openUC2 ImSwitch fork** – if customized from the original repo
- **Firmware GitHub repository** – ESP32-based controller code Github
- **openUC2 Docs** – general documentation including assembly and setup GH Pages
- **ImSwitch documentation** – architecture, modules, API references Read-The-Docs or openUC2 GH Pages

#### Docker and OS Images

- **GitHub Container Registry** – Docker images for ImSwitch and drivers ARM64 Version, AMD64
- **Raspberry Pi OS (ImSwitch version)** – prebuilt image with instructions Forklift Repo, [Prebuild OS on Zenodo]
- **Documentation for Forklift Integration** – GH Pages

#### Web-Based Tools

- **Firmware Flash Tool (WebSerial)** – browser-based firmware upload Firmware Flasher
- **Firmware Test Tool** – live hardware interaction in the browser Firmware Testing
- **Blockly Workflow Editor** – visual workflow editor generating JSON Static ImSwitch Frontend
- **ImSwitch Web GUI** – hosted or local version (if available for testing/demo) Live ImSwitch Demo hosted on AWS - it may be down

#### Integration and APIs

- **Arkitekt project** – network-based microservice control environment https://github.com/arkitektio
- **Arkitekt documentation** – node types, setup, integration with ImSwitch https://arkitekt.live/
- **openUC2 Arkitekt Plugin** – Sample implementation to expose services: https://github.com/openUC2/imswitch-arkitekt-next

#### Interfacing and Extensions

- **Jupyter Notebook Examples** – smart microscopy, autofocus, time-lapse sample implementatoins
- **Hardware Integration Guides** – Opentrons, robotic arms, etc. (comming soon)
- **ESP32 WebSerial interface** – for external components (lasers, motors, galvos)
- **ESP32 CAN/I2C Extensions** – for adding peripherals CAN Interface

#### Community and Support

- **openUC2 Forum/Discussions** – GitHub or external forum Discourse Forum

## Automation of spatial single-cell analysis with Cytely

**Karl Johansson, Johannes Kumra Ahnlinde, Oscar André, Philip Nordenfelt, Pontus Nordenfelt** Cytely AB, Medicon Village, Lund, Sweden

### Specific Focus and scientific questions asked

Traditional microscopy excels at capturing subcellular structures but typically falls short in providing comprehensive population-level context, which is crucial to understanding cellular heterogeneity and rare cellular events. Conventional high-resolution imaging approaches tend to focus on a limited number of cells, thereby introducing potential biases and limiting statistical power. Conversely, high-content screening often compromises spatial resolution, making it reducing the ability to discern subcellular phenotypes. The central challenge addressed by Cytely is how to efficiently automate microscopy workflows to integrate high-resolution, single-cell spatial context with broad, population-level insights, enabling unbiased and reproducible analysis of diverse cellular phenotypes, particularly rare and dynamic events.

#### Key findings and innovations

Cytely is based on the scientific foundation coming from the Nordenfelt Lab [1], leveraging principles from datadriven microscopy (DDM) to significantly enhance accuracy and throughput in spatial single-cell analysis. The platform integrates whole populations overview and data at a single-cell level, allowing for rapid, unbiased identification of cellular phenotypes (Fig. 1). Cytely effectively characterizes thousands of cells based on features such as cell morphology, fluorescence intensity, and subcellular organization. This approach facilitates the precise selection and revisiting of cells of interest, allowing in a second step for high-resolution spatial imaging of rare or phenotypically distinct cellular subsets (Fig. 2).

**Fig. 1:**
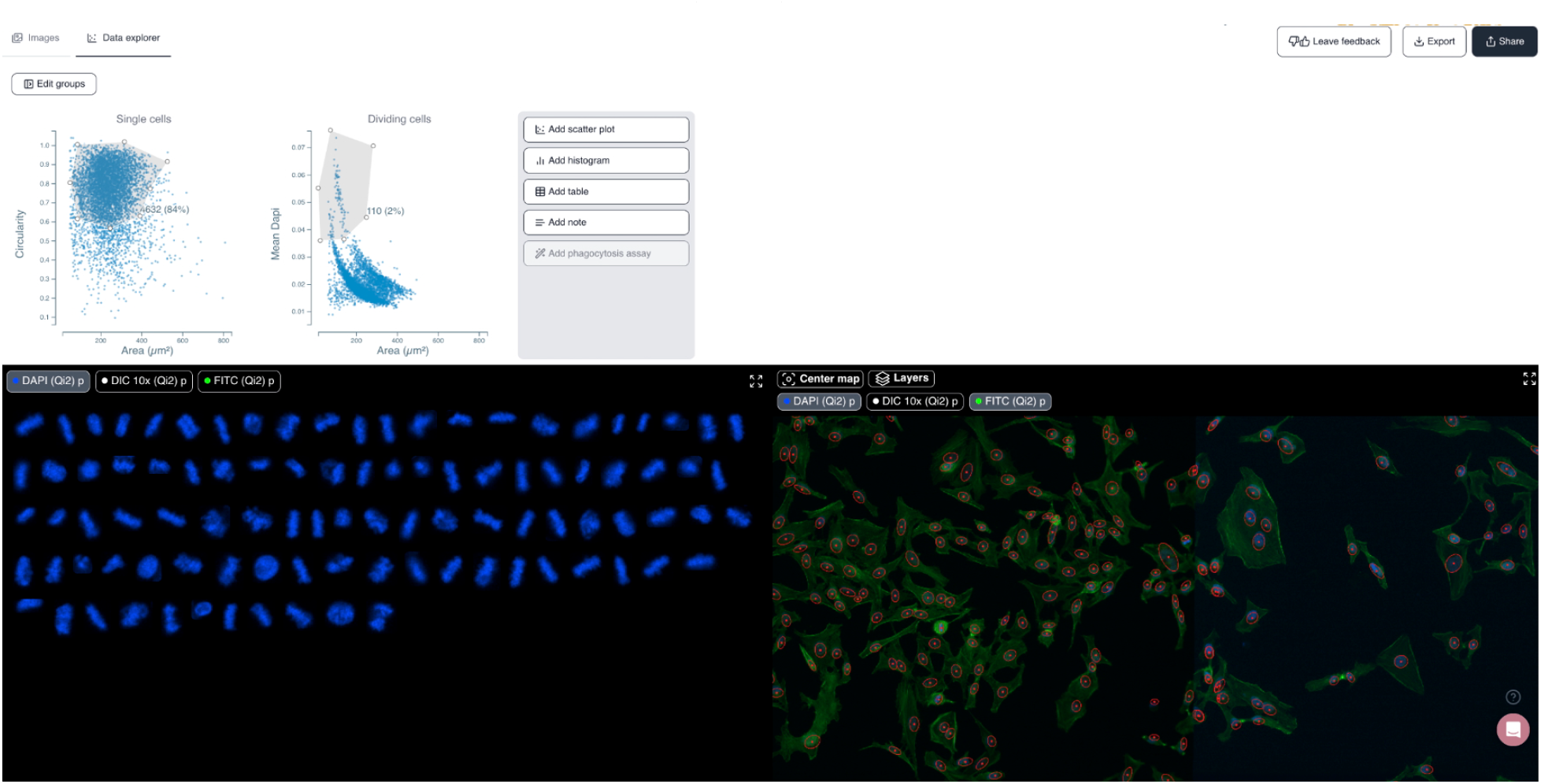
Cytely’s standard browser GUI. The mitotic cell subpopulation of a HeLa cell sample is displayed (link to dataset).

**Fig. 2:**
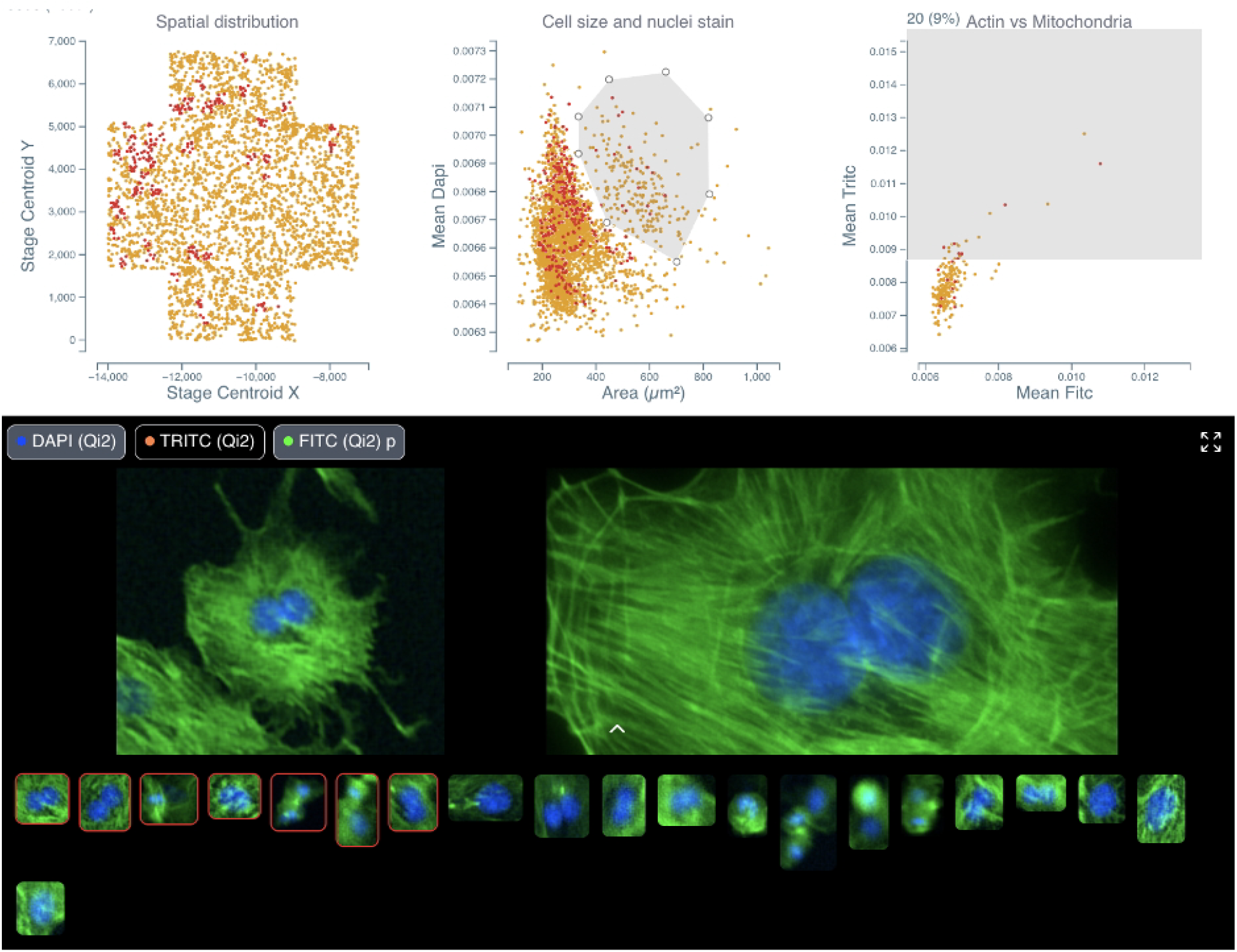
Side by side comparison of initial low-resolution image and curated high-resolution reimaging of the same cell. The dataset consists of bovine pulmonary artery endothelial cells (link to dataset).

In experiments investigating cellular phagocytosis, Cytely demonstrated its unique ability to visualize the spatial localization of engulfed particles within individual cells, identifying heterogeneity in uptake rates that traditional bulk analyses missed. Similarly, Cytely’s approach enabled automated detection and characterization of cellular events, such as cell division or apoptosis, and allowed for curation of such data through high-resolution re-imaging at a significantly higher rate and accuracy compared to conventional microscopy.

#### Methodology and implementation details

Cytely is implemented as a microscopy-based, hardware-agnostic platform compatible with various imaging systems equipped with motorized stages. The workflow begins with a low-resolution, population-level image acquisition phase that rapidly segments and extracts morphological and intensity-based cellular features. The software leverages classical segmentation approaches combined with advanced AI algorithms to robustly characterize cellular phenotypes in real-time, enabling immediate identification of target cell populations. Once specific cellular subsets are identified, Cytely triggers high-resolution spatial imaging, automatically directing microscopy resources to acquire detailed, single-cell data. The entire workflow, from image acquisition to analysis and data documentation, is managed within an intuitive, integrated software interface, eliminating the need for complex scripting or manual intervention. Image analysis is performed using defined schemas, allowing for easy replication and all datasets are stored centrally and can be shared both internally and externally, facilitating data transparency (example of shared dataset).

#### Contributions to interoperability

Cytely’s platform is completely hardware agnostic and is designed to integrate seamlessly with widely used microscopy systems. The modular and flexible nature of Cytely’s analysis pipeline enables straightforward adaptation to diverse experimental setups and research needs. Its compatibility with open standards and extensive device support promotes broad community adoption and facilitates integration into existing laboratory workflows.

#### Limitations

While Cytely robustly handles standard microscopy scenarios, particularly challenging tissues with complex segmentation requirements may still present difficulties. Additionally, the system currently relies on user-driven parameter tuning prior to experiments to optimize analysis outcomes, particularly in contexts with significant biological variability. Future iterations aim to implement adaptive parameter tuning algorithms to minimize manual intervention further and enhance the usability and reproducibility of experiments.

## Visual programming for Smart Microscopy with NIS-Elements (Nikon)

**Nelda Antonovaite, Andrii Rogov**, Nikon Europe B.V.

### Approach to smart microscopy

NIS-Elements is a comprehensive software platform designed for all Nikon microscopes, supporting a wide array of imaging modalities such as upright and inverted widefield, confocal (AX and NSPARC), multiphoton, STORM, and SIM. Beyond Nikon hardware, NIS-Elements allows seamless integration with numerous third-party devices, including spinning-disks, deepSIM modules, cameras, light sources, stages, and modular illumination devices such as photostimulation, TIRF, and FRAP. Additionally, the software natively supports NI DAQ boards, enabling low-latency control of almost any external device. With the release of NIS-Elements version 6.10, a Python package was introduced, further expanding device control capabilities and workflow customization.

This extensive compatibility empowers users to perform sophisticated experiments, including advanced feedback microscopy. In addition to robust device control, developers of feedback microscopy require software that is flexible enough to enable customized and automated acquisition workflows, as well as the integration of real-time, on-the-fly analysis. NIS-Elements addresses these requirements by allowing full automation of decision-making in complex experiments, reducing human error and bias, increasing throughput, and freeing microscopists from repetitive tasks.

Within NIS-Elements, the JOBS module enables users to create custom acquisition workflows that incorporate on-the-fly analysis through the General Analysis module (GA3). JOBS utilizes a visual programming interface, where users can drag and drop various tasks into a sequence.These tasks come with prebuilt parameter options and support advanced features like loops, phases, and wizards. This approach ensures that every step of the acquisition process can be fully customized and automated, with analysis results dynamically guiding subsequent acquisition steps.

The advanced GA3 image analysis module also employs visual programming, with a key distinction: its interconnected, branched architecture is designed to handle multiple channels, binary masks, measurement results, and graphs in parallel, rather than in a simple sequential order. GA3 leverages the full suite of image processing tools in NIS-Elements, including AI-based denoising, deblurring, deconvolution, and user-trained deep learning models for segmentation and signal-to-noise ratio enhancement (NIS.ai module), all of which can improve the speed and quality of analysis.

Both JOBS and GA3 are designed to be intuitive, allowing users to create sophisticated acquisition and analysis workflows with minimal training. They offer substantial flexibility, supporting tasks such as wellplate and slide scanning, multi-pass acquisitions (e.g., low magnification followed by high magnification, or combinations of widefield and confocal imaging), conditional imaging (e.g., stopping when a certain number of cells are imaged or when rare events are detected), and tracking.

For advanced applications, Python scripting is available both within JOBS (for external device control) and GA3 (for open-source analysis tools, e.g., CellPose-SAM). Together, NIS-Elements with JOBS and GA3 bridges the gap between limited, easy-to-use preconfigured software and complex, highly flexible scripting-based solutions.

#### Implementation and key features

##### JOBS module

The JOBS editor features an intuitive graphical interface that is easy to learn and requires no programming experience. Users can drag and drop over 180 available tasks to create custom image acquisition workflows (see Figure 1 and Video 1). These tasks include:

- Defining and manipulating wellplates and scanning areas for various types of holders
- Creating imaging loops, phases, and conditional branches (if/else)
- Setting acquisition parameters such as objective, channels, timelapses, Z-stacks, triggering, image capture, and saving
- Defining focusing procedures
- Performing on-the-fly image analysis and interactive functions, such as prompting the user with questions
- Device control, Python scripting, macros, and more

Each JOBS function also has a panel where optional parameters can be selected (see Figure 2 and Video 1). Furthermore, software commands are recorded in Command History and can be used as macro commands. You can edit or extend them using macro command library. If any specific software function for device control or handling of images is not available within JOBS editor, it can be included as a macro command. Similarly, Python editor allows to control devices that are not integrated within the NIS-Elements software or NI DAQ board. Once a JOBS workflow is created, a wizard interface can be configured by simply marking JOBS tasks that should be added to the wizard. Additional descriptions, remarks, recommendations or even images can be added to explain to the user the purpose of the settings and guide through the selection. When launched, the user only needs to select variables that fit their experiment (see Figure 3 and Video 2). This way, even less experienced users can run complicated imaging experiments.

Finally, JOBS execution can be monitored by showing captured images as well as plotting analysis results (see Video 2). Additional interactions can be set up to ask the user to make the decision on the go or make the decision automatically based on conditions or analysis results.

#### General Analysis module (GA3)

Similar to JOBS, GA3 uses a visual programming approach, where prebuilt image analysis functions—covering processing, segmentation, measurement, data handling, and more—can be dragged and dropped into a dynamically connected analysis sequence (see Figure 4). GA3 offers over 400 functions, organized into twelve categories, with tools available for both 2D and 3D data:

- Image Processing
- Image Operations
- Multi-Dimensional Processing & Conversions
- Segmentation
- Binary Processing
- Binary Operations
- Measurement
- Data Management
- Results & Graphs
- Sources & Reference
- NIS.ai

If a specific function is missing or if an open-source tool is required, the GA3 Python editor provides further extensibility.

Analyses can be previewed live on selected regions of interest, allowing users to inspect and quickly adjust settings while building workflows. The interactive graphs and tables make it easy to explore analysis data in relation to the image and generated binaries.

Any GA3 workflow can be incorporated into a JOBS workflow, using the captured image as input and producing outputs such as tables (with points, regions, wells, etc.) or binary masks. These outputs can then be used within JOBS to drive decisions based on analysis results (see examples in Figure 5 and Video 3).

#### Current bottlenecks and roadmap

A primary limitation is the speed of image analysis for very large datasets. This can be improved through the use of more powerful computers, development of GPU implementations for computationally intensive tasks and ensuring that only relevant data - excluding empty frames - is being acquired. Additionally, parallel computing solutions are being explored. While GA3 and JOBS are designed for usability, there remains a learning curve, so additional improvements are planned to streamline workflow creation. Importantly, the integration of Python in both GA3 and JOBS has opened the door to utilizing open-source Python packages and community-developed resources—a direction that continues to be actively developed.

#### Figures and video screencasts

**Figure 1.**
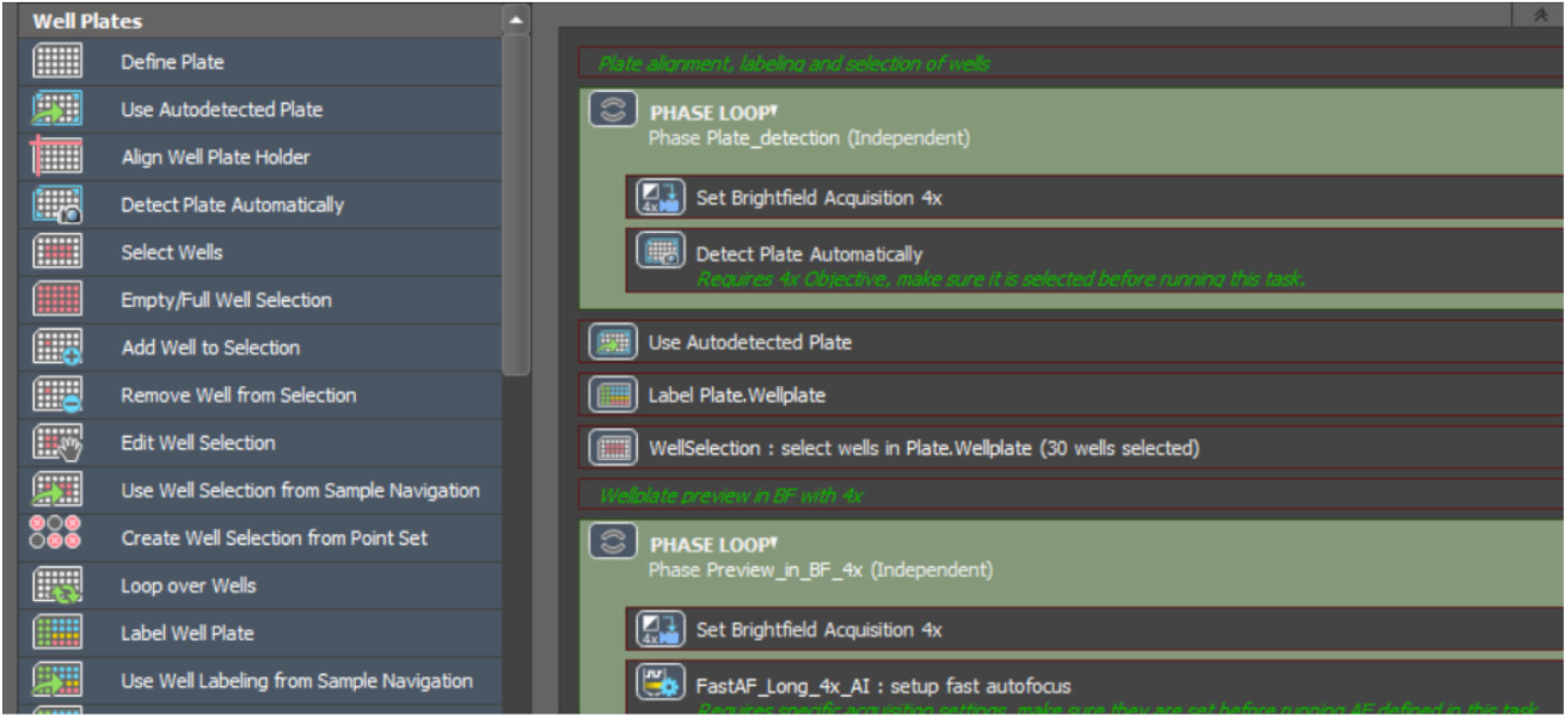
JOBS editor. Acquisition and analysis functions can be dragged into the sequence to create a custom acquisition workflow for feedback microscopy.

**Figure 2.**
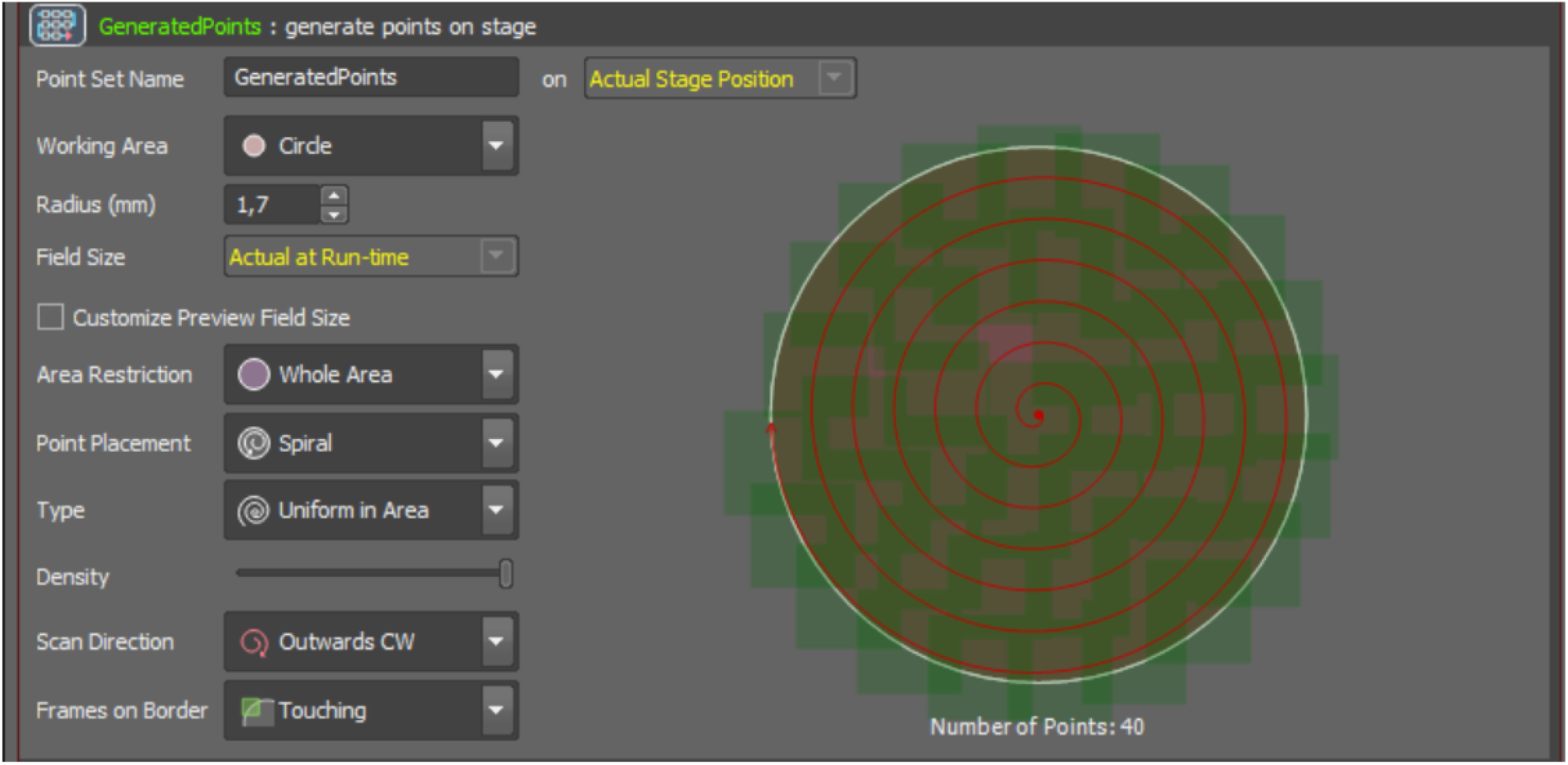
Example of optional parameters for „generate points” function within JOBS editor.

**Figure 3.**
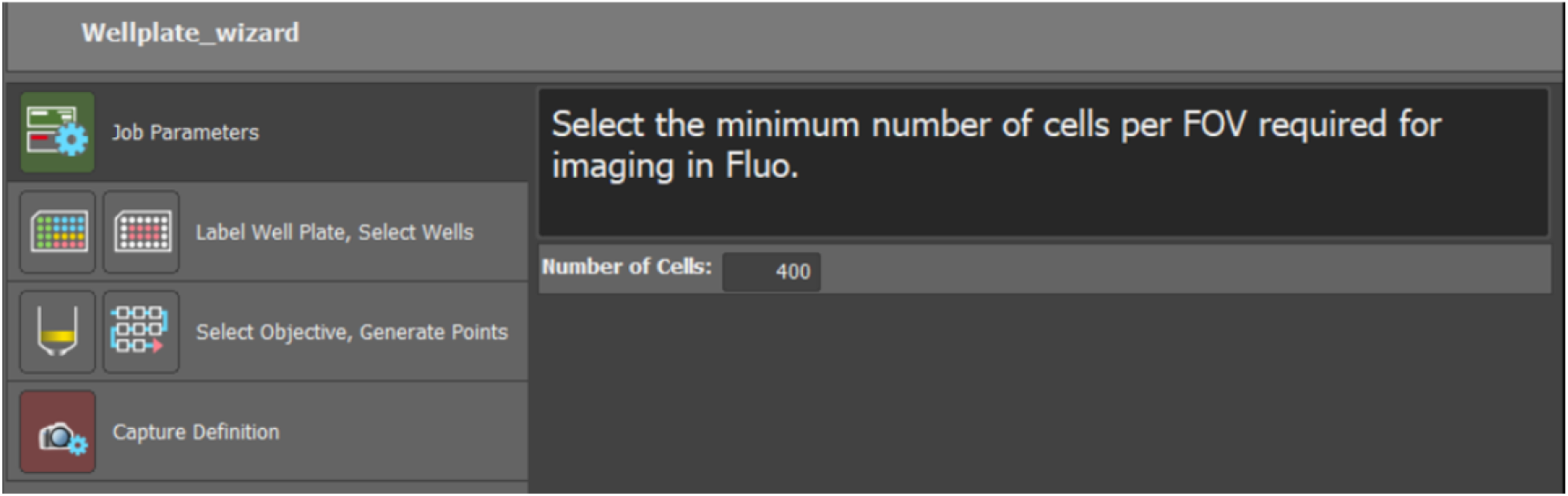
JOBS workflow can be converted into a simple wizard enabling users that are not experienced with microscopy, device control and image analysis, to run feedback microscopy experiments.

**Figure 4.**
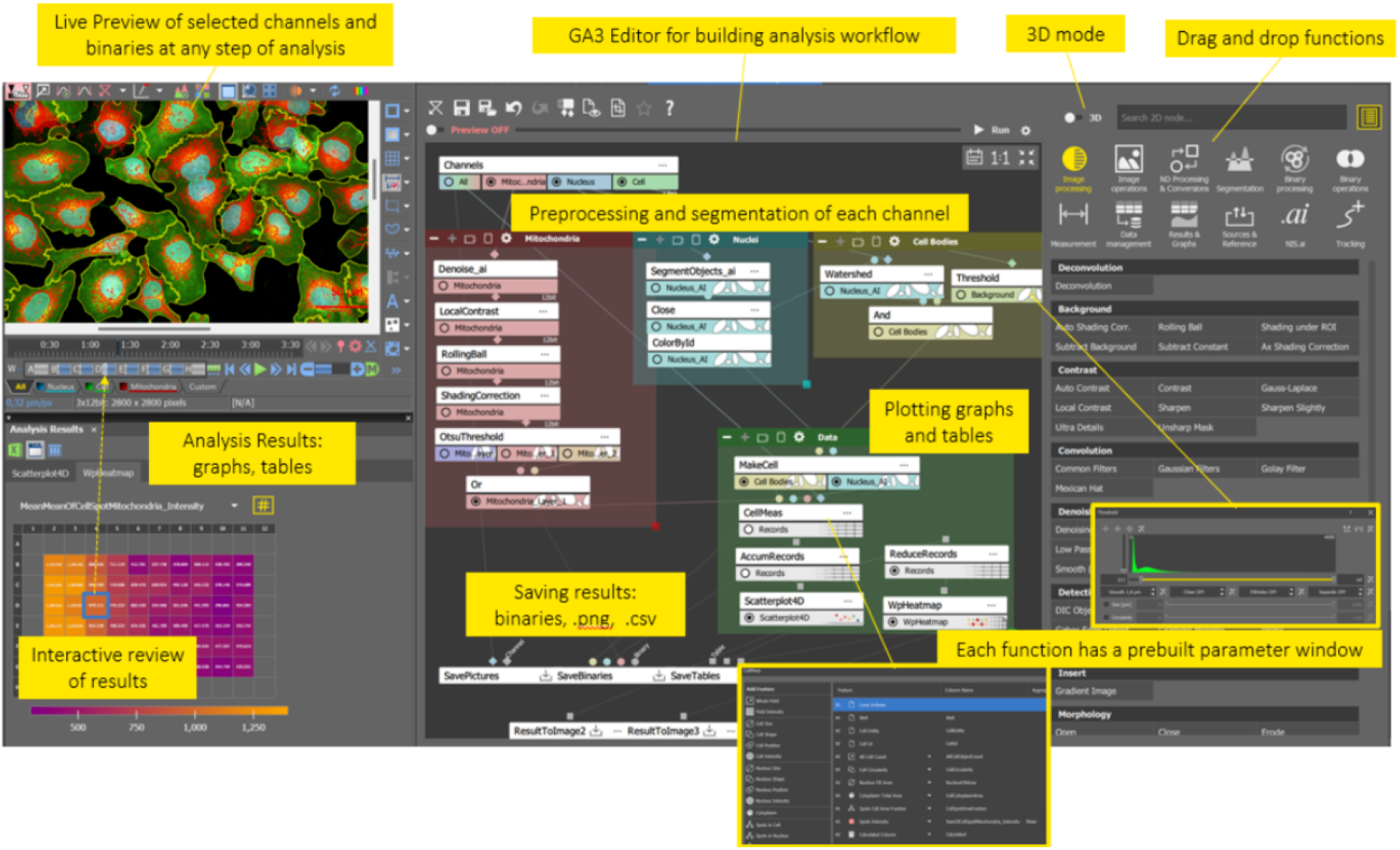
GA3 editor is a visual programming-based interface where completely customized image analysis workflows can be created and then integrated into JOBS editor enabling feedback microscopy experiments within NIS-Elements.

**Figure 5.**
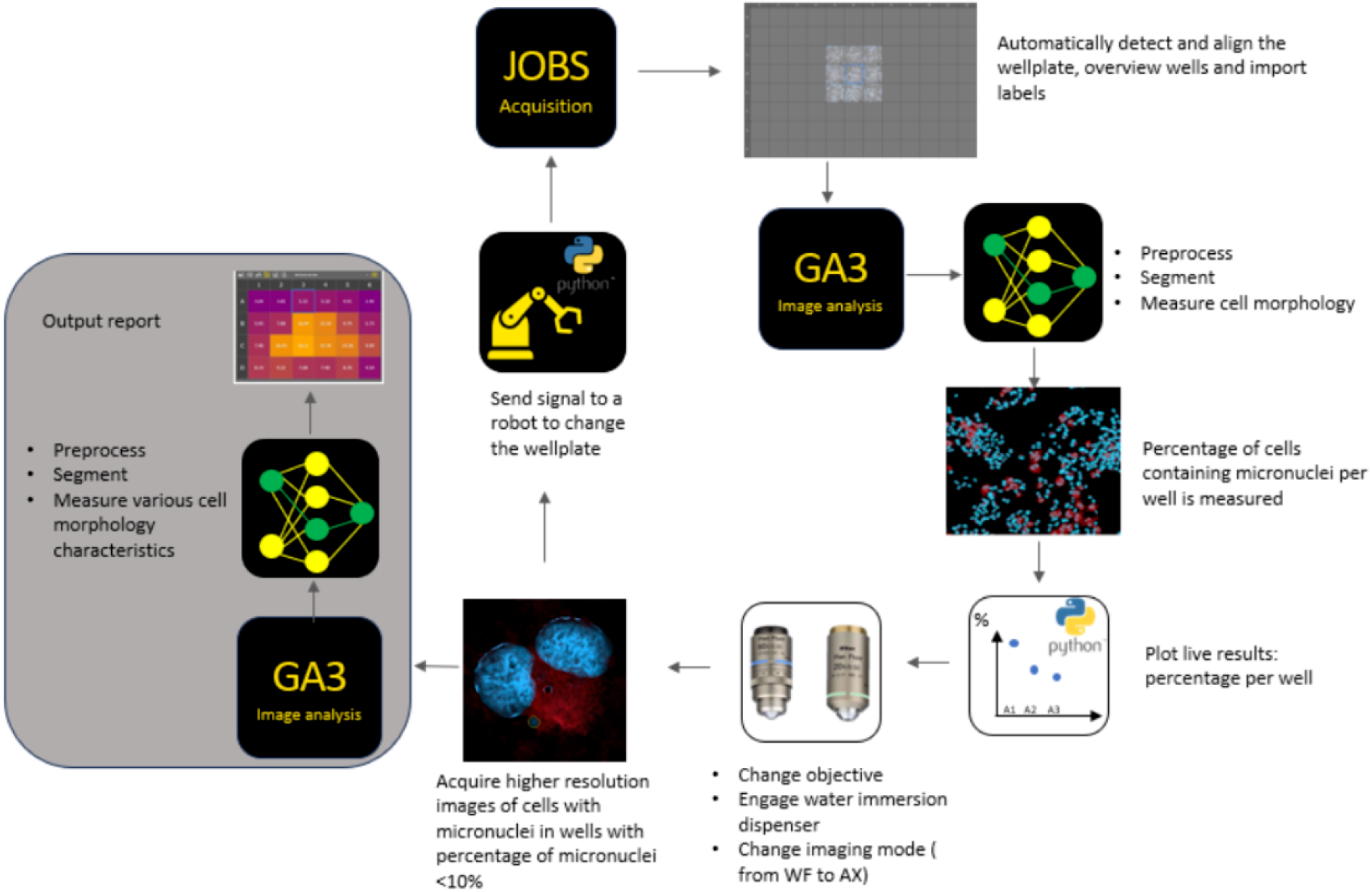
Example of the feedback microscopy experiment workflow within NIS-Elements software using JOBS and GA3 modules. Video 1 : Simple Wellplate acquisition Video 2 : Target Cell Count Python Plot Video 3: Automated Feedback Microscopy

#### Documentation and references

A lot of information about the use of GA3, JOBS and other preprocessing tools (deconvolution, NIS.ai) are included in the NIS-Elements help files. Some examples are also included.
A GitHub page is available that include JOBS, GA3 and Python examples:
  - JOBs examples
  - GA3 and Python examples
NIS-Elements tutorials are continuously being updated on Nikon’s e-learning platform which is available to all Nikon customers by submitting a request at Nikon e-learning
Some examples of publications that have implemented feedback microscopy are listed below:
  - High-Throughput Microenvironment Microarray (MEMA) High-Resolution Imaging | SpringerLink
  - Data-driven microscopy allows for automated context-specific acquisition of high-fidelity image data: Cell Reports Methods
  - Robotic data acquisition with deep learning enables cell image–based prediction of transcriptomic phenotypes
Other Nikon resources can be found here: Resources | Nikon Europe B.V., including application notes featuring feedback microscopy:
  - High content screening lifetime data_ A feedback mic Exp WF
  - Feedback microscopy using Nikon AX confocal microscopy and JOBS software to bridge mm to µm structures

## Footnotes

1 smartmicroscopy.github.io

2 eurobioimaging.eu

3 smartmicroscopy.github.io/implementations

4 smartmicroscopy.github.io

5 smartmicroscopy.org

6 forum.image.sc/tag/smart-microscopy

7 image.sc

8 microlist.org

## Bibliography

[1] Leonor Morgado, Estibaliz Gómez-de-Mariscal, Hannah S. Heil, and Ricardo Henriques. The rise of data-driven microscopy powered by machine learning. Journal of Microscopy, 295(2):85–92, March 2024.ISSN 1365-2818. doi:10.1111/jmi.13282.

[2] Henry Pinkard and Laura Waller. Microscopes are coming for your job. Nature Methods, 19 (10):1175–1176, September 2022.ISSN 1548-7105. doi:10.1038/s41592-022-01566-4.

[3] Martin Schorb, Isabella Haberbosch, Wim J. H. Hagen, Yannick Schwab, and David N. Mastronarde. Software tools for automated transmission electron microscopy. Nature Methods, 16(6):471–477, May 2019.ISSN 1548-7105. doi:10.1038/s41592-019-0396-9.

[4] Oscar André, Johannes Kumra Ahnlide, Nils Norlin, Vinay Swaminathan, and Pontus Nordenfelt. Data-driven microscopy allows for automated context-specific acquisition of high-fidelity image data. Cell Reports Methods, 3(3):100419, March 2023.ISSN 2667-2375. doi:10.1016/j.crmeth.2023.100419.

[5] Catherine Bouchard, Theresa Wiesner, Andréanne Deschênes, Anthony Bilodeau, Benoît Turcotte, Christian Gagné, and Flavie Lavoie-Cardinal. Resolution enhancement with a task-assisted gan to guide optical nanoscopy image analysis and acquisition. Nature Machine Intelligence, 5(8):830–844, July 2023.ISSN 2522-5839. doi:10.1038/s42256-023-00689-3.

[6] Ewald R. Weibel. An automatic sampling stage microscope for stereology. Journal of Microscopy, 91(1):1–18, February 1970.ISSN 1365-2818. doi:10.1111/j.1365-2818.1970.tb02199.x.

[7] J F Brenner, B S Dew, J B Horton, T King, P W Neurath, and W D Selles. An automated microscope for cytologic research a preliminary evaluation. Journal of Histochemistry & Cytochemistry, 24(1):100–111, January 1976.ISSN 1551-5044. doi:10.1177/24.1.1254907.

[8] Howard C. Berg. How to track bacteria. Review of Scientific Instruments, 42(6):868–871, June 1971.ISSN 1089-7623. doi:10.1063/1.1685246.

[9] J.M.S. Prewitt. Intelligent microscopes: a scientific poem. in COMPSAC 79. Proceedings. Computer Software and The IEEE Computer Society’s Third International Applications Conference, 1979., pages 477–487, 1979. doi:10.1109/CMPSAC.1979.762545.

[10] Yimo Zhang. An automated microscope system for cell image processing. in International Conference on Optoelectronic Science and Engineering ‘90, page 211. SPIE, July 1990. doi:10.1117/12.2294849.

[11] E. Lægsgaard, F. Besenbacher, K. Mortensen, and I. Stensgaard. A fully automated, ‘thimble-size’ scanning tunnelling microscope. Journal of Microscopy, 152(3):663–669, December 1988.ISSN 1365-2818. doi:10.1111/j.1365-2818.1988.tb01435.x.

[12] F H Deindoerfer, W B Boris, J R Gangwer, C W Laird, and J W Tittsler. Automated intelligent microscopy (aim) and its potential application in the clinical laboratory. Clinical Chemistry, 28(9):1910–1916, September 1982.ISSN 1530-8561. doi:10.1093/clinchem/28.9.1910.

[13] J. M.S Prewitt, B. H. Mayall, and M. L. Mendelsohn. Pictorial Data Processing Methods In Microscopy. In Harold L Kasnitz, editor, Filmed Data and Computers, volume 0006, pages 138–151. International Society for Optics and Photonics, SPIE, 1966. doi:10.1117/12.971062.

[14] Christian Conrad, Annelie Wünsche, Tze Heng Tan, Jutta Bulkescher, Frank Sieckmann, Fatima Verissimo, Arthur Edelstein, Thomas Walter, Urban Liebel, Rainer Pepperkok, and Jan Ellenberg. Micropilot: automation of fluorescence microscopy–based imaging for systems biology. Nature Methods, 8(3):246–249, January 2011.ISSN 1548-7105. doi:10.1038/nmeth.1558.

[15] Yuki Tsukada and Koichi Hashimoto. Feedback regulation of microscopes by image processing. Development, Growth & Differentiation, 55(4):550–562, April 2013.ISSN 1440-169X. doi:10.1111/dgd.12056.

[16] Cyril Bourgenot, Christopher D. Saunter, Jonathan M. Taylor, John M. Girkin, and Gordon D. Love. 3d adaptive optics in a light sheet microscope. Optics Express, 20(12):13252, May 2012.ISSN 1094-4087. doi:10.1364/oe.20.013252.

[17] Aurore Masson, Paul Escande, Céline Frongia, Grégory Clouvel, Bernard Ducommun, and Corinne Lorenzo. High-resolution in-depth imaging of optically cleared thick samples using an adaptive spim. Scientific Reports, 5(1), November 2015.ISSN 2045-2322. doi:10.1038/srep16898.

[18] Loïc A Royer, William C Lemon, Raghav K Chhetri, Yinan Wan, Michael Coleman, Eugene W Myers, and Philipp J Keller. Adaptive light-sheet microscopy for long-term, high-resolution imaging in living organisms. Nature Biotechnology, 34(12):1267–1278, October 2016.ISSN 1546-1696. doi:10.1038/nbt.3708.

[19] Loïc A. Royer, William C. Lemon, Raghav K. Chhetri, and Philipp J. Keller. A practical guide to adaptive light-sheet microscopy. Nature Protocols, 13(11):2462–2500, October 2018.ISSN 1750-2799. doi:10.1038/s41596-018-0043-4.

[20] Lucas von Chamier, Romain F. Laine, and Ricardo Henriques. Artificial intelligence for microscopy: what you should know. Biochemical Society Transactions, 47(4):1029–1040, July 2019.ISSN 1470-8752. doi:10.1042/bst20180391.

[21] Sergei V. Kalinin, Evgheni Strelcov, Alex Belianinov, Suhas Somnath, Rama K. Vasudevan, Eric J. Lingerfelt, Richard K. Archibald, Chaomei Chen, Roger Proksch, Nouamane Laanait, and Stephen Jesse. Big, deep, and smart data in scanning probe microscopy. ACS Nano, 10(10):9068–9086, September 2016.ISSN 1936-086X. doi:10.1021/acsnano.6b04212.

[22] Joanna W. Pylvänäinen, Estibaliz Gómez-de Mariscal, Ricardo Henriques, and Guillaume Jacquemet. Live-cell imaging in the deep learning era. Current Opinion in Cell Biology, 85: 102271, December 2023.ISSN 0955-0674. doi:10.1016/j.ceb.2023.102271.

[23] Sergei V. Kalinin, Debangshu Mukherjee, Kevin Roccapriore, Benjamin J. Blaiszik, Ayana Ghosh, Maxim A. Ziatdinov, Anees Al-Najjar, Christina Doty, Sarah Akers, Nageswara S. Rao, Joshua C. Agar, and Steven R. Spurgeon. Machine learning for automated experimentation in scanning transmission electron microscopy. npj Computational Materials, 9(1), December 2023.ISSN 2057-3960. doi:10.1038/s41524-023-01142-0.

[24] Inês Cunha, Emma Latron, Sebastian Bauer, Daniel Sage, and Juliette Griffié. Machine learning in microscopy – insights, opportunities and challenges. Journal of Cell Science, 137(20), October 2024.ISSN 1477-9137. doi:10.1242/jcs.262095.

[25] Nico Scherf and Jan Huisken. The smart and gentle microscope. Nature Biotechnology, 33 (8):815–818, August 2015.ISSN 1546-1696. doi:10.1038/nbt.3310.

[26] P. S. Kesavan and Pontus Nordenfelt. Reconceptualizing smart microscopy: From data collection to knowledge creation by multi-agent integration, 2025. doi:10.48550/ARXIV.2505.20466.

[27] Willi L. Stepp, Emine Berna Durmus, Santiago N. Rodriguez Alvarez, Juan C. Landoni, Giorgio Tortarolo, Kyle M. Douglass, Martin Weigert, and Suliana Manley. Smart hybrid microscopy for cell-friendly detection of rare events. bioRxiv, April 2025. doi:10.1101/2025.04.04.647219.

[28] Scott Brooks, Sara Toral-Pérez, David S. Corcoran, Karl Kilborn, Brian Bodensteiner, Hella Baumann, Nigel J. Burroughs, Andrew D. McAinsh, and Till Bretschneider. Celfdrive: Artificial intelligence assisted microscopy for automated detection of rare events. bioRxiv, 2024. doi:10.1101/2024.10.17.618897.

[29] Dora Mahecic, Willi L. Stepp, Chen Zhang, Juliette Griffié, Martin Weigert, and Suliana Manley. Event-driven acquisition for content-enriched microscopy. Nature Methods, 19(10): 1262–1267, September 2022.ISSN 1548-7105. doi:10.1038/s41592-022-01589-x.

[30] Yu Shi, Jimmy S. Tabet, Daniel E. Milkie, Timothy A. Daugird, Chelsea Q. Yang, Alex T. Ritter, Andrea Giovannucci, and Wesley R. Legant. Smart lattice light-sheet microscopy for imaging rare and complex cellular events. Nature Methods, 21(2):301–310, January 2024.ISSN 1548-7105. doi:10.1038/s41592-023-02126-0.

[31] Jonatan Alvelid, Martina Damenti, Chiara Sgattoni, and Ilaria Testa. Event-triggered sted imaging. Nature Methods, 19(10):1268–1275, September 2022.ISSN 1548-7105. doi:10.1038/s41592-022-01588-y.

[32] Khalid A. Ibrahim, Camille Cathala, Carlo Bevilacqua, Lely Feletti, Robert Prevedel, Hilal A. Lashuel, and Aleksandra Radenovic. Self-driving microscopy detects the onset of protein aggregation and enables intelligent brillouin imaging. Nature Communications, 16:6699, 7 2025.ISSN 2041-1723. doi:10.1038/s41467-025-60912-0.

[33] Marco Raffaele Cosenza, Alice Gaiatto, Büsra Erarslan Uysal, Álvaro Andrades Delgado, Nina Luisa Sautter, Michael Adrian Jendrusch, Sonia Zumalave Duro, Tobias Rausch, Aliaksandr Halavatyi, Eva-Maria Geissen, Patrick Hasenfeld, Isidro Cortes-Ciriano, Andreas Kulozik, Rainer Pepperkok, and Jan O. Korbel. Origins ofde novochromosome rearrangements unveiled by coupled imaging and genomics. bioRxiv, August 2024. doi:10.1101/2024.08.15.607890.

[34] Christian Tischer, Volker Hilsenstein, Kirsten Hanson, and Rainer Pepperkok. Adaptive fluorescence microscopy by online feedback image analysis, page 489–503. Elsevier, 2014. doi:10.1016/b978-0-12-420138-5.00026-4.

[35] Josiah B. Passmore, Alfredo Rates, Jakob Schröder, Menno T. P. van Laarhoven, Vincent J. W. Hellebrekers, Henrik G. van Hoef, Antonius J. M. Geurts, Wendy van Straaten, Wilco Nijenhuis, Florian Berger, Carlas S. Smith, Ihor Smal, and Lukas C. Kapitein. Outcome-driven microscopy: Closed-loop optogenetic control of cell biology. BioRxiv, December 2024. doi:10.1101/2024.12.12.628240.

[36] Max Heydasch, Lucien Hinderling, Jakobus van Unen, Maciej Dobrzynski, and Olivier Pertz. Gtpase activating protein dlc1 spatio-temporally regulates rho signaling. eLife Sciences Publications, Ltd, December 2023. doi:10.7554/elife.90305.1.

[37] Lucien Hinderling, Alex E. Landolt, Benjamin Grädel, Laurent Dubied, Cédric Zahni, Moritz Kwasny, Agne Frismantiene, Talley Lambert, Maciej Dobrzynski, and Olivier Pertz. Realtime feedback control microscopy for automation of optogenetic targeting. bioRxiv, August 2025. doi:10.1101/2025.08.17.670729.

[38] Harrison R. Oatman, Beena C. Lad, and Jared Toettcher. Pyclm: programming-free, closed-loop microscopy for real-time measurement, segmentation, and optogenetic stimulation. bioRxiv, September 2025. doi:10.1101/2025.08.29.673155.

[39] Josiah B. Passmore, Wilco Nijenhuis, and Lukas C. Kapitein. From observing to controlling: Inducible control of organelle dynamics and interactions. Current Opinion in Cell Biology, 71:69–76, August 2021.ISSN 0955-0674. doi:10.1016/j.ceb.2021.02.002.

[40] Galit Lahav, Nitzan Rosenfeld, Alex Sigal, Naama Geva-Zatorsky, Arnold J Levine, Michael B Elowitz, and Uri Alon. Dynamics of the p53-mdm2 feedback loop in individual cells. Nature Genetics, 36(2):147–150, January 2004.ISSN 1546-1718. doi:10.1038/ng1293.

[41] Harish Shankaran, Danielle L Ippolito, William B Chrisler, Haluk Resat, Nikki Bollinger, Lee K Opresko, and H Steven Wiley. Rapid and sustained nuclear–cytoplasmic erk oscillations induced by epidermal growth factor. Molecular Systems Biology, 5(1), January 2009.ISSN 1744-4292. doi:10.1038/msb.2009.90.

[42] D. E. Nelson, A. E. C. Ihekwaba, M. Elliott, J. R. Johnson, C. A. Gibney, B. E. Foreman, G. Nelson, V. See, C. A. Horton, D. G. Spiller, S. W. Edwards, H. P. McDowell, J. F. Unitt, E. Sullivan, R. Grimley, N. Benson, D. Broomhead, D. B. Kell, and M. R. H. White. Oscillations in nf-κb signaling control the dynamics of gene expression. Science, 306(5696): 704–708, October 2004.ISSN 1095-9203. doi:10.1126/science.1099962.

[43] Jared E Toettcher, Delquin Gong, Wendell A Lim, and Orion D Weiner. Light-based feedback for controlling intracellular signaling dynamics. Nature Methods, 8(10):837–839, September 2011.ISSN 1548-7105. doi:10.1038/nmeth.1700.

[44] Armin Baumschlager, Stephanie K. Aoki, and Mustafa Khammash. Dynamic blue light-inducible t7 rna polymerases (opto-t7rnaps) for precise spatiotemporal gene expression control. ACS Synthetic Biology, 6(11):2157–2167, October 2017.ISSN 2161-5063. doi:10.1021/acssynbio.7b00169.

[45] Evan J Olson, Lucas A Hartsough, Brian P Landry, Raghav Shroff, and Jeffrey J Tabor. Characterizing bacterial gene circuit dynamics with optically programmed gene expression signals. Nature Methods, 11(4):449–455, March 2014.ISSN 1548-7105. doi:10.1038/nmeth.2884.

[46] Jeffrey J. Tabor, Anselm Levskaya, and Christopher A. Voigt. Multichromatic control of gene expression in escherichia coli. Journal of Molecular Biology, 405(2):315–324, January 2011.ISSN 0022-2836. doi:10.1016/j.jmb.2010.10.038.

[47] Andreas Milias-Argeitis, Marc Rullan, Stephanie K. Aoki, Peter Buchmann, and Mustafa Khammash. Automated optogenetic feedback control for precise and robust regulation of gene expression and cell growth. Nature Communications, 7(1), August 2016.ISSN 2041-1723. doi:10.1038/ncomms12546.

[48] Stephanie K. Aoki, Gabriele Lillacci, Ankit Gupta, Armin Baumschlager, David Schwein-gruber, and Mustafa Khammash. A universal biomolecular integral feedback controller for robust perfect adaptation. Nature, 570(7762):533–537, June 2019.ISSN 1476-4687. doi:10.1038/s41586-019-1321-1.

[49] Gabriele Lillacci, Yaakov Benenson, and Mustafa Khammash. Synthetic control systems for high performance gene expression in mammalian cells. Nucleic Acids Research, 46(18): 9855–9863, September 2018.ISSN 1362-4962. doi:10.1093/nar/gky795.

[50] Marc Rullan, Dirk Benzinger, Gregor W. Schmidt, Andreas Milias-Argeitis, and Mustafa Khammash. An optogenetic platform for real-time, single-cell interrogation of stochastic transcriptional regulation. Molecular Cell, 70(4):745–756.e6, May 2018.ISSN 1097-2765. doi:10.1016/j.molcel.2018.04.012.

[51] Jose L. Avalos, Jared E. Toettcher, and Evan M. Zhao. System and method of optogenetically controlling metabolic pathways for the production of chemicals, March 2024. Published 07 Mar 2024; application US18/502,266.

[52] Han Woong Yoo, Michel Verhaegen, Martin E. van Royen, and Georg Schitter. Automated adjustment of aberration correction in scanning confocal microscopy. in 2012 IEEE International Instrumentation and Measurement Technology Conference Proceedings, page 1083–1088. IEEE, May 2012. doi:10.1109/i2mtc.2012.6229195.

[53] Kai Wang, Daniel E Milkie, Ankur Saxena, Peter Engerer, Thomas Misgeld, Marianne E Bronner, Jeff Mumm, and Eric Betzig. Rapid adaptive optical recovery of optimal resolution over large volumes. Nature Methods, 11(6):625–628, April 2014.ISSN 1548-7105. doi:10.1038/nmeth.2925.

[54] Tsung-Li Liu, Srigokul Upadhyayula, Daniel E. Milkie, Ved Singh, Kai Wang, Ian A. Swinburne, Kishore R. Mosaliganti, Zach M. Collins, Tom W. Hiscock, Jamien Shea, Abraham Q. Kohrman, Taylor N. Medwig, Daphne Dambournet, Ryan Forster, Brian Cunniff, Yuan R, and Eric Betzig. Observing the cell in its native state: Imaging subcellular dynamics in multicellular organisms. Science (New York, N.Y.), 360: 1–53, 2018.

[55] Faris Abouakil, Huicheng Meng, Marie-Anne Burcklen, Hervé Rigneault, Frédéric Galland, and Loïc LeGoff. An adaptive microscope for the imaging of biological surfaces. Light: Science & Applications, 10(1), October 2021.ISSN 2047-7538. doi:10.1038/s41377-021-00649-9.

[56] Martin Weigert, Uwe Schmidt, Tobias Boothe, Andreas Müller, Alexandr Dibrov, Akanksha Jain, Benjamin Wilhelm, Deborah Schmidt, Coleman Broaddus, Siân Culley, Mauricio Rocha-Martins, Fabián Segovia-Miranda, Caren Norden, Ricardo Henriques, Marino Zerial, Michele Solimena, Jochen Rink, Pavel Tomancak, Loic Royer, Florian Jug, and Eugene W. Myers. Content-aware image restoration: pushing the limits of fluorescence microscopy. Nature Methods, 15(12):1090–1097, November 2018.ISSN 1548-7105. doi:10.1038/s41592-018-0216-7.

[57] Cassandra Tong Ye, Jiashu Han, Kunzan Liu, Anastasios Angelopoulos, Linda Griffith, Kristina Monakhova, and Sixian You. Learned, uncertainty-driven adaptive acquisition for photon-efficient scanning microscopy. Optics Express, 33(6):12269, March 2025.ISSN 1094-4087. doi:10.1364/oe.542640.

[58] Maxim A Ziatdinov, Ayana Ghosh, and Sergei V Kalinin. Physics makes the difference: Bayesian optimization and active learning via augmented gaussian process. Machine Learning: Science and Technology, 3(1):015003, February 2022.ISSN 2632-2153. doi:10.1088/2632-2153/ac4baa.

[59] Maxim A. Ziatdinov, Yongtao Liu, Anna N. Morozovska, Eugene A. Eliseev, Xiaohang Zhang, Ichiro Takeuchi, and Sergei V. Kalinin. Hypothesis learning in automated experiment: Application to combinatorial materials libraries. Advanced Materials, 34(20), April 2022.ISSN 1521-4095. doi:10.1002/adma.202201345.

[60] Juliane Liepe, Sarah Filippi, Micha Komorowski, and Michael P. H. Stumpf. Maximizing the information content of experiments in systems biology. PLoS Computational Biology, 9(1): e1002888, January 2013.ISSN 1553-7358. doi:10.1371/journal.pcbi.1002888.

[61] M F Bolus, A A Willats, C J Rozell, and G B Stanley. State-space optimal feedback control of optogenetically driven neural activity. Journal of Neural Engineering, 18(3):036006, March 2021.ISSN 1741-2552. doi:10.1088/1741-2552/abb89c.

[62] Xavier Moreno, Staffan Al-Kadhimi, Jonatan Alvelid, Andreas Bodén, and Ilaria Testa. Imswitch: Generalizing microscope control in python. Journal of Open Source Software, 6 (64):3394, August 2021.ISSN 2475-9066. doi:10.21105/joss.03394.

[63] Xavier Casas Moreno, Mariline Mendes Silva, Johannes Roos, Francesca Pennacchietti, Nils Norlin, and Ilaria Testa. An open-source microscopy framework for simultaneous control of image acquisition, reconstruction, and analysis. HardwareX, 13:e00400, March 2023.ISSN 2468-0672. doi:10.1016/j.ohx.2023.e00400.

[64] Arthur D. Edelstein, Mark A. Tsuchida, Nenad Amodaj, Henry Pinkard, Ronald D. Vale, and Nico Stuurman. Advanced methods of microscope control using µmanager software. Journal of Biological Methods, 1(2):1, November 2014.ISSN 2326-9901. doi:10.14440/jbm.2014.36.

[65] Arthur Edelstein, Nenad Amodaj, Karl Hoover, Ron Vale, and Nico Stuurman. Computer control of microscopes using µmanager. Current Protocols in Molecular Biology, 92(1), October 2010.ISSN 1934-3647. doi:10.1002/0471142727.mb1420s92.

[66] Johannes Roos, Stéphane Bancelin, Tom Delaire, Alexander Wilhelmi, Florian Levet, Maren Engelhardt, Virgile Viasnoff, Rémi Galland, U. Valentin Nägerl, and Jean-Baptiste Sibarita. Arkitekt: streaming analysis and real-time workflows for microscopy. Nature Methods, 21 (10):1884–1894, September 2024.ISSN 1548-7105. doi:10.1038/s41592-024-02404-5.

[67] Ome-zarr: a cloud-optimized bioimaging file format with international community support. Histochemistry and Cell Biology, 160(3):223–251, July 2023.ISSN 1432-119X. doi:10.1007/s00418-023-02209-1.

[68] Antonio Z Politi, Yin Cai, Nike Walther, M Julius Hossain, Birgit Koch, Malte Wachsmuth, and Jan Ellenberg. Quantitative mapping of fluorescently tagged cellular proteins using fcs-calibrated four-dimensional imaging. Nature Protocols, 13(6):1445–1464, May 2018.ISSN 1750-2799. doi:10.1038/nprot.2018.040.

[69] Talley Lambert, Federico Gasparoli, Ian Hunt-Isaak, Gabriel Selzer, Willi L Stepp, and Lucien Hinderling. pymmcore-plus/useq-schema: v0.7.3, 2025. doi:10.5281/ZENODO.15692749.

[70] Carsen Stringer and Marius Pachitariu. Cellpose3: one-click image restoration for improved cellular segmentation. Nature Methods, 22(3):592–599, February 2025.ISSN 1548-7105. doi:10.1038/s41592-025-02595-5.

[71] Lucien Hinderling, Guillaume Witz, Roman Schwob, Ana Stojiljkovic, Maciej Dobrzynski, Mykhailo Vladymyrov, Joel Frei, Benjamin Graedel, Agne Frismantiene, and Olivier Pertz. Convpaint - universal framework for interactive pixel classification using pretrained neural networks. bioRxiv, September 2024. doi:10.1101/2024.09.12.610926.

[72] Daniel B. Allan, Thomas Caswell, Nathan C. Keim, Casper M. van der Wel, and Ruben W. Verweij. soft-matter/trackpy: v0.6.4, 2024. doi:10.5281/ZENODO.12708864.

[73] Kristina Ulicna, Giulia Vallardi, Guillaume Charras, and Alan R. Lowe. Automated deep lineage tree analysis using a bayesian single cell tracking approach. Frontiers in Computer Science, 3:92, 2021.ISSN 2624-9898. doi:10.3389/fcomp.2021.734559.

[74] Benedict Diederich, René Lachmann, Swen Carlstedt, Barbora Marsikova, Haoran Wang, Xavier Uwurukundo, Alexander S. Mosig, and Rainer Heintzmann. A versatile and customizable low-cost 3d-printed open standard for microscopic imaging. Nature Communications, 11(1), November 2020.ISSN 2041-1723. doi:10.1038/s41467-020-19447-9.

[75] Niamh Burke, Gesine Müller, Vittorio Saggiomo, Amy Ruth Hassett, Jérôme Mutterer, Patrick Ó Súilleabháin, Daniel Zakharov, Donal Healy, Emmanuel G Reynaud, and Mark Pickering. EnderScope: a low-cost 3D printer-based scanning microscope for microplastic detection. Philos. Trans. A Math. Phys. Eng. Sci., 382(2274):20230214, July 2024.

[76] Sirine Gharbi, Emeric Poiraud, Hugo Le Guenno, Erwan Grandgirard, Charly Rousseau, Niamh Burke, Jerome Mutterer, and David Rousseau. Enderscope.py: A library for computational imaging using the EnderScope automated microscope. SoftwareX, 31(102210): 102210, September 2025.

[77] Talley Lambert, Ian Hunt-Isaak, Federico Gasparoli, Willi L Stepp, Mark Tsuchida, Alex Landolt, Curtis Rueden, and Lucien Hinderling. pymmcore-plus/pymmcore-plus: v0.15.0, 2025. doi:10.5281/ZENODO.15634905.

[78] Henry Pinkard, Nico Stuurman, Ivan E. Ivanov, Nicholas M. Anthony, Wei Ouyang, Bin Li, Bin Yang, Mark A. Tsuchida, Bryant Chhun, Grace Zhang, Ryan Mei, Michael Anderson, Douglas P. Shepherd, Ian Hunt-Isaak, Raymond L. Dunn, Wiebke Jahr, Saul Kato, Loïc A. Royer, Jay R. Thiagarajah, Kevin W. Eliceiri, Emma Lundberg, Shalin B. Mehta, and Laura Waller. Pycro-manager: open-source software for customized and reproducible microscope control. Nature Methods, 18(3):226–228, March 2021.ISSN 1548-7105. doi:10.1038/s41592-021-01087-6.

[79] Mathias Hammer, Maximiliaan Huisman, Alessandro Rigano, Ulrike Boehm, James J. Chambers, Nathalie Gaudreault, Alison J. North, Jaime A. Pimentel, Damir Sudar, Peter Bajcsy, Claire M. Brown, Alexander D. Corbett, Orestis Faklaris, Judith Lacoste, Alex Laude, Glyn Nelson, Roland Nitschke, Farzin Farzam, Carlas S. Smith, David Grunwald, and Caterina Strambio-De-Castillia. Towards community-driven metadata standards for light microscopy: tiered specifications extending the ome model. Nature Methods, 18(12): 1427–1440, December 2021.ISSN 1548-7105. doi:10.1038/s41592-021-01327-9.

[80] Alessandro Rigano, Shannon Ehmsen, Serkan Utku Öztürk, Joel Ryan, Alexander Balashov, Mathias Hammer, Koray Kirli, Ulrike Boehm, Claire M. Brown, Karl Bellve, James J. Chambers, Andrea Cosolo, Robert A. Coleman, Orestis Faklaris, Kevin E. Fogarty, Thomas Guilbert, Anna B. Hamacher, Michelle S. Itano, Daniel P. Keeley, Susanne Kunis, Judith Lacoste, Alex Laude, Willa Y. Ma, Marco Marcello, Paula Montero-Llopis, Glyn Nelson, Roland Nitschke, Jaime A. Pimentel, Stefanie Weidtkamp-Peters, Peter J. Park, Burak H. Alver, David Grunwald, and Caterina Strambio-De-Castillia. Micro-meta app: an interactive tool for collecting microscopy metadata based on community specifications. Nature Methods, 18(12):1489–1495, December 2021.ISSN 1548-7105. doi:10.1038/s41592-021-01315-z.

[81] Danny Driess, Fei Xia, Mehdi S. M. Sajjadi, Corey Lynch, Aakanksha Chowdhery, Brian Ichter, Ayzaan Wahid, Jonathan Tompson, Quan Vuong, Tianhe Yu, Wenlong Huang, Yevgen Chebotar, Pierre Sermanet, Daniel Duckworth, Sergey Levine, Vincent Vanhoucke, Karol Hausman, Marc Toussaint, Klaus Greff, Andy Zeng, Igor Mordatch, and Pete Florence. Palm-e: An embodied multimodal language model. in arXiv preprint arXiv:2303.03378, 2023.

[82] Loïc A. Royer. Omega — harnessing the power of large language models for bioimage analysis. Nature Methods, 21(8):1371–1373, June 2024.ISSN 1548-7105. doi:10.1038/s41592-024-02310-w.

## BIBLIOGRAPHY

[1] Max Heydasch et al. “GTPase activating protein DLC1 spatio-temporally regulates Rho signaling.” In: eLife (Dec. 2023). DOI: 10.7554/elife.90305.1. URL: http://dx.doi.org/10.7554/eLife.90305.1.

[2] Lucien Hinderling et al. “Real-time feedback control microscopy for automation of optogenetic targeting.” In: (Aug. 2025). DOI: 10.1101/2025.08.17.670729. URL: http://dx.doi.org/10.1101/2025.08.17.670729.

## BIBLIOGRAPHY

[1] Alexander Kirillov et al. Segment Anything. 2023. DOI: 10.48550/ARXIV.2304.02643. URL: https://arxiv.org/abs/2304.02643.

[2] Dominik Niopek et al. “Optogenetic control of nuclear protein export.” In: Nature Communications 7.1 (Feb. 2016). ISSN: 2041-1723. DOI: 10.1038/ncomms10624. URL: http://dx.doi.org/10.1038/ncomms10624.

[3] Katsuhiko Ogata. Modern control engineering. 4th ed. Upper Saddle River, NJ: Pearson, Nov. 2001.

[4] Josiah B. Passmore et al. “Outcome-Driven Microscopy: Closed-Loop Optogenetic Control of Cell Biology.” In: bioRxiv (Dec. 2024). DOI: 10.1101/2024.12.12.628240. URL: http://dx.doi.org/10.1101/2024.12.12.628240.

[5] Nicholas Sofroniew et al. napari: a multi-dimensional image viewer for Python. 2025. DOI: 10.5281/ZENODO.3555620. URL: https://zenodo.org/doi/10.5281/zenodo.3555620.

## BIBLIOGRAPHY

[1] Andreas Brunner et al. “Quantitative imaging of loop extruders rebuilding interphase genome architecture after mitosis.” In: Journal of Cell Biology 224.3 (Jan. 2025). ISSN: 1540-8140. DOI: 10.1083/jcb.202405169. URL: http://dx.doi.org/10.1083/jcb.202405169.

[2] Marco Raffaele Cosenza et al. “Origins ofde novochromosome rearrangements unveiled by coupled imaging and genomics.” In: bioRxiv (Aug. 2024). DOI: 10.1101/2024.08.15.607890. URL: http://dx.doi.org/10.1101/2024.08.15.607890.

[3] Timothy Fuqua et al. “An open-source semi-automated robotics pipeline for embryo immunohistochemistry.” In: Scientific Reports 11.1 (May 2021). ISSN: 2045-2322. DOI: 10.1038/s41598-021-89676-5. URL: http://dx.doi.org/10.1038/s41598-021-89676-5.

[4] Timothy Fuqua et al. “Dense and pleiotropic regulatory information in a developmental enhancer.” In: Nature 587.7833 (Oct. 2020), pp. 235–239. ISSN: 1476-4687. DOI: 10.1038/s41586-020-2816-5. URL: http://dx.doi.org/10.1038/s41586-020-2816-5.

[5] Tobias Kletter et al. “Cell State-Specific Cytoplasmic Material Properties Control Spindle Architecture and Scaling.” In: bioRxiv (July 2024). DOI: 10.1101/2024.07.22.604615. URL: http://dx.doi.org/10.1101/2024.07.22.604615.

[6] Antonio Z Politi et al. “Quantitative mapping of fluorescently tagged cellular proteins using FCS-calibrated four-dimensional imaging.” In: Nature Protocols 13.6 (May 2018), pp. 1445–1464. ISSN: 1750-2799. DOI: 10.1038/nprot.2018.040. URL: http://dx.doi.org/10.1038/nprot.2018.040.

[7] Sanjana Singh et al. “Phenotype-based single-cell transcriptomics reveal compensatory pathways involved in Golgi organization and associated transport.” In: bioRxiv (Dec. 2022). DOI: 10.1101/2022.12.02.518815. URL: http://dx.doi.org/10.1101/2022.12.02.518815.

## BIBLIOGRAPHY

[1] Jonathan U Harrison et al. “Kinetochore tracking in 3D from lattice light-sheet imaging data with KiT.” In: Bioinformatics 38.12 (May 2022). Ed. by Jonathan Wren, pp. 3315–3317. ISSN: 1367-4811. DOI: 10.1093/bioinformatics/btac330. URL: http://dx.doi.org/10.1093/bioinformatics/btac330.

[2] Glenn Jocher, Jing Qiu, and Ayush Chaurasia. “Ultralytics YOLO.” Version 8.0.0. In: GitHub (Jan. 2023). URL: https://github.com/ultralytics/ultralytics.

[3] Li Ren Kong et al. “A glycolytic metabolite bypasses “two-hit” tumor suppression by BRCA2.” In: Cell 187.9 (Apr. 2024), 2269–2287.e16. ISSN: 0092-8674. DOI: 10.1016/j.cell.2024.03.006. URL: http://dx.doi.org/10.1016/j.cell.2024.03.006.

[4] Yian Zhao et al. “DETRs Beat YOLOs on Real-time Object Detection.” In: bioRxiv (2023). DOI: 10.48550/ARXIV.2304.08069. URL: https://arxiv.org/abs/2304.08069.

## BIBLIOGRAPHY

[1] Sirine Gharbi et al. “Enderscope.py: A library for computational imaging using the EnderScope automated microscope.” In: SoftwareX 31 (Sept. 2025), p. 102210. ISSN: 2352-7110. DOI: 10.1016/j.softx.2025.102210. URL: http://dx.doi.org/10.1016/j.softx.2025.102210.

## BIBLIOGRAPHY

[1] Elliot Dine et al. “Protein Phase Separation Provides Long-Term Memory of Transient Spatial Stimuli.” In: Cell Systems 6.6 (June 2018), 655–663.e5. ISSN: 2405-4712. DOI: 10.1016/j.cels.2018.05.002. URL: http://dx.doi.org/10.1016/j.cels.2018.05.002.

[2] Arthur Edelstein et al. “Computer Control of Microscopes Using µManager.” In: Current Protocols in Molecular Biology 92.1 (Oct. 2010). ISSN: 1934-3647. DOI: 10.1002/0471142727.mb1420s92. URL: http://dx.doi.org/10.1002/0471142727.mb1420s92.

[3] Payam E Farahani et al. “pYtags enable spatiotemporal measurements of receptor tyrosine kinase signaling in living cells.” In: eLife 12 (May 2023). ISSN: 2050-084X. DOI: 10.7554/elife.82863. URL: http://dx.doi.org/10.7554/eLife.82863.

[4] Nury Kim et al. “Spatiotemporal Control of Fibroblast Growth Factor Receptor Signals by Blue Light.” In: Chemistry and Biology 21.7 (July 2014), pp. 903–912. ISSN: 1074-5521. DOI: 10.1016/j.chembiol.2014.05.013. URL: http://dx.doi.org/10.1016/j.chembiol.2014.05.013.

[5] Harrison R. Oatman, Beena C. Lad, and Jared Toettcher. “PyCLM: programming-free, closed-loop microscopy for real-time measurement, segmentation, and optogenetic stimulation.” In: bioRxiv (Sept. 2025). DOI: 10.1101/2025.08.29.673155. URL: http://dx.doi.org/10.1101/2025.08.29.673155.

[6] Sergi Regot et al. “High-Sensitivity Measurements of Multiple Kinase Activities in Live Single Cells.” In: Cell 157.7 (June 2014), pp. 1724–1734. ISSN: 0092-8674. DOI: 10.1016/j.cell.2014.04.039. URL: http://dx.doi.org/10.1016/j.cell.2014.04.039.

[7] Jared E. Toettcher, Orion D. Weiner, and Wendell A. Lim. “Using Optogenetics to Interrogate the Dynamic Control of Signal Transmission by the Ras/Erk Module.” In: Cell 155.6 (Dec. 2013), pp. 1422–1434. ISSN: 0092-8674. DOI: 10.1016/j.cell.2013.11.004. URL: http://dx.doi.org/10.1016/j.cell.2013.11.004.

## BIBLIOGRAPHY

[1] Oscar André et al. “Data-driven microscopy allows for automated context-specific acquisition of high-fidelity image data.” In: Cell Reports Methods 3.3 (Mar. 2023), p. 100419. ISSN: 2667-2375. DOI: 10.1016/j.crmeth.2023.100419. URL: http://dx.doi.org/10.1016/j.crmeth.2023.100419.

